# DiffDomain enables identification of structurally reorganized topologically associating domains

**DOI:** 10.1101/2022.12.05.519135

**Authors:** Dunming Hua, Ming Gu, Xiao Zhang, Yanyi Du, Hangcheng Xie, Li Qi, Xiangjun Du, Zhidong Bai, Xiaopeng Zhu, Dechao Tian

**Author notes:** These four authors contributed equally (Dunming Hua, Ming Gu, Xiao Zhang, Yanyi Du).

## Abstract

Topologically associating domains (TADs) are critical structural units in three-dimensional genome organization of mammalian genome. Dynamic reorganizations of TADs between health and disease states are associated with transcription and other essential genome functions. However, computational methods that can identify reorganized TADs are still in the early stages of development. Here, we present DiffDomain, an algorithm leveraging high-dimensional random matrix theory to identify structurally reorganized TADs using chromatin contact maps. Method comparison using multiple real Hi-C datasets reveals that DiffDomain outperforms alternative methods for FPRs, TPRs, and identifying a new subtype of reorganized TADs. The robustness of DiffDomain and its biological applications are demonstrated by applying on Hi-C data from different cell types and disease states. Identified reorganized TADs are associated with structural variations and changes in CTCF binding sites and other epigenomic changes. By applying to a single-cell Hi-C data from mouse neuronal development, DiffDomain can identify reorganized TADs between cell types with reasonable reproducibility using pseudo-bulk Hi-C data from as few as 100 cells per condition. Moreover, DiffDomain reveals that TADs have differential cell-to-population variability and heterogeneous cell-to-cell variability. Therefore, DiffDomain is a statistically sound method for better comparative analysis of TADs using both Hi-C and single-cell Hi-C data.

## Introduction

Recent development of mapping technologies such as Hi-C [1] that probes the 3D genome organization reveals that a chromosome is divided into topologically associating domains (TADs) [2, 3]. TADs are genomic regions where chromatin loci are more frequently interacting with other chromatin loci within the TAD than with those from outside of the TAD. TADs are functional units for transcriptional regulation by constraining interactions between enhancers and promoters [4], for example. Although TADs are stable between cell types as revealed by earlier studies [2, 5], there are growing evidence for TAD reorganization in diseases [6–8], cell differentiation [5, 9, 10], somatic cellular reprogramming [11], between neuronal cell types [12], and between species [13, 14]. For example, extensive reorganizations of TADs are observed during somatic cell reprogramming, associating with dynamics of transcriptional regulation and changes in cellular identity [11]. TADs are also variable among individual cells as revealed by single-cell studies [15–20] and live-cell imaging [21]. Thus, it is important to identify reorganized TADs through comparative analysis to further understand the functional relevance of 3D genome organization, a major priority of current work in the field [22].

The majority of current methods call a reorganized TAD if at least one of its two boundaries changed between two conditions [11, 12, 23–28]. These methods enable easy integration with other analysis pipelines and identified reorganized TADs have clear biological interpretation. However, they fail in identifying reorganized TADs without changes in boundaries, in addition to lacking statistical tests to differentiate random perturbations and significant structural reorganization of a TAD. Only a few non-parametric statistical methods are proposed to call TAD reorganization [29–32]. These methods define the structural similarity of a TAD by statistics from two Hi-C contact matrices, such as the stratum-adjusted correlation coefficient used by DiffGR [30]. Distributions of the statistics on pairs of simulated Hi-C matrices are then used to compute empirical *P* values. However, these nonparametric statistical methods are conservative (see our own comparison later). TADs in high-resolution Hi-C data are relatively small. The median size of TADs is 185 kb [33]. The small size feature of TADs poses another computational challenge for identifying structurally rewired TADs using low-resolution Hi-C data. Importantly, identifying reorganized TADs using emerging single-cell Hi-C (scHi-C) data is largely under-explored. Other methods are developed for comparing Hi-C matrices at different scales and for different purposes: quantifying similarities of genome-wide Hi-C contact matrices [34, 35], identifying differential A/B compartments [36], and identifying differential chromatin interactions [37–39]. However, these methods are not tailored to compare Hi-C contact matrices at TAD-level, not optimal for identifying reorganized TADs (see our own comparison later). Therefore, new algorithms are needed to fill these gaps.

Here, we develop DiffDomain, a new parametric statistical method for identifying reorganized TADs. Its inputs are two Hi-C contact matrices from two biological conditions and a set of TADs called in biological condition 1. This setting enables straightforward integration of DiffDomain with other analysis pipelines of Hi-C data such as TAD calling and integrative analysis of multi-omics data. For each TAD, DiffDomain *directly* computes a difference matrix and then normalizes it properly, skipping the challenging normalization steps for individual Hi-C contact matrices. DiffDomain then borrows well-established theoretical results in random matrix theory to compute a *P* value. We show that the assumptions of DiffDomain are reasonable. Method comparisons on real data reveal that DiffDomain has substantial advantages over alternative methods in false positive rates and accuracy in identifying truly reorganized TADs. Reorganized TADs identified by DiffDomain are biologically relevant in different human cell lines and disease states. Application to scHi-C data reveals that DiffDomain can identify reorganized TADs between cell types and TADs with differential variabilities among individual cells within the same cell type. Moreover, DiffDomain can quantify cell-to-cell variability of TADs between individual cells. Together, these analyses demonstrate the power of DiffDomoain for better identification of structurally reorganized TADs using both bulk Hi-C and single-cell Hi-C data.

## Results

### Overview of DiffDomain

The workflow of DiffDomain is illustrated in Figure 1. Its input are a set of TADs called in biological condition 1 and their corresponding Hi-C contact matrices from biological conditions 1 and 2 (Fig. 1A). In this paper, TADs are called by Arrowhead [33] and Hi-C contact matrices are KR-normalized, unless otherwise stated. Our goal is to test if each TAD identified in biological condition 1 has significant structural reorganization in biological condition 2. The core of DiffDomain is formulating the problem as a hypothesis testing problem, where the null hypothesis is that the TAD doesn’t undergo significant structural reorganization in condition 2. To achieve this goal, for each TAD with *N* bins, DiffDomain extracts the *N ×N* KR-normalized Hi-C contact matrices specific to the TAD region from the two biological conditions, which are denoted as ***A***_1_ and ***A***_2_ (Fig. 1A). Note that ***A***_1_ and ***A***_2_ are *N ×N* submatrices of the genome-wide Hi-C contact matrices. DiffDomain first log-transform them to adjust for the exponential decay of Hi-C contacts with increased 1D distances between chromosome bins. Their difference log(***A***_1_) −log(***A***_2_) is calculated and denoted by ***D*** (Fig. 1B). ***D*** is further normalized by a 1D distance-stratified standardization procedure, similar to the procedures in HiC-DC+ [38] and SnapHiC [40]. Specifically, each *d*-off diagonal part of ***D*** is subtracted by its sample mean and divided by its sample standard deviation (Fig. 1C), −*N* + 2 ≤*d ≤N −*2, reducing 1D distance-dependence of values in ***D*** and differences caused by variation in read depths between two biological conditions (see Supplementary Fig. S1 for two more detailed visualization). Intuitively, if a TAD is not significantly reorganized, normalized ***D*** would resemble a white noise random matrix, enabling us to borrow theoretical results in random matrix theory. Under the null hypothesis, DiffDomain assumes that 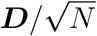 is a generalized Wigner matrix (Fig. 1D), a well-studied random matrix model. Its largest eigenvalue *λ*_*N*_ is proved to be fluctuating around 2. Armed with the fact, DiffDomain reformulates the reorganized TAD identification problem into the hypothesis testing problem:

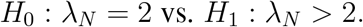

The key theoretical results empowering DiffDomain is that *θ*_*N*_ = *N* ^2*/*3^(*λ*_*N*_ *−*2), a normalized *λ*_*N*_, asymptotically follows a Tracy-Widom distribution with *β* = 1, denoted as *TW*_1_. Thus, *θ*_*N*_ is chosen as the test statistic and the one-sided *P* value is calculated as 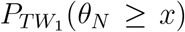. *H*_0_ is rejected if the *P* value is less than a predefined significant level *α*, which is 0.05 in this paper (Fig. 1E). The pseudocode is shown in Supplementary Methods A.1. For a set of TADs, *P* values are adjusted for multiple comparisons using Benjamini-Hochberg (BH) method as the default. Once DiffDomain identifies the subset of reorganized TADs, it further classifies them into six subtypes based on changes in their boundaries, beneficial for downstream biological analyses and interpretations (Fig. 1F). TAD reorganization subtypes are verified by aggregation peak analyses (APA) on multiple real datasets (Supplementary Fig. S2). A few reorganized TADs in real Hi-C data are shown in Fig. 1G. Details are described in Methods.

**Figure 1:**
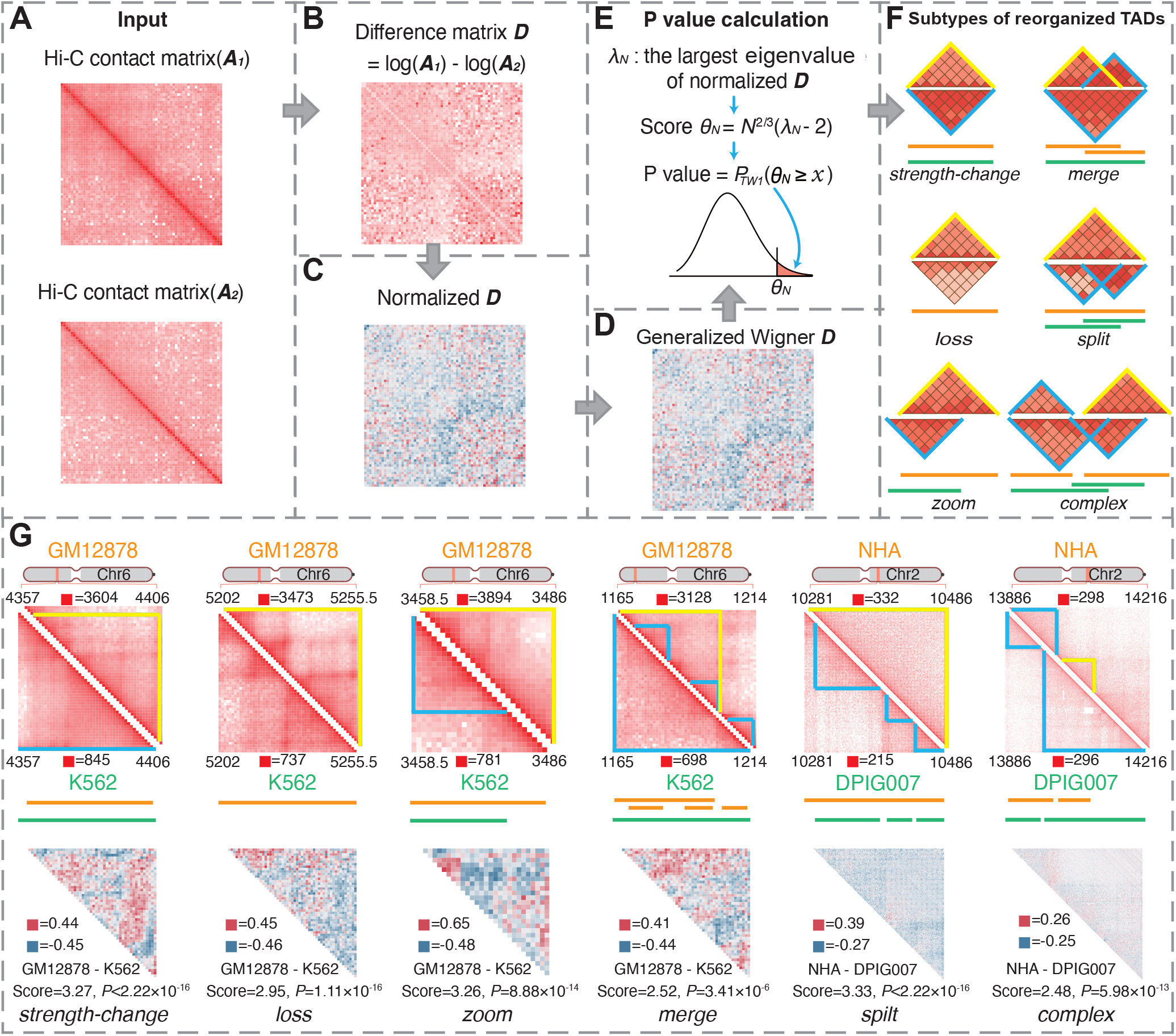
DiffDomain workflow and example outputs. (***A***) Input are a TAD in condition 1 and its two Hi-C contact matrices (***A***_1_ and ***A***_2_) in two biological conditions 1 and 2. (***B***) Difference between log-transformed ***A***_1_ and ***A***_2_, which is denoted as ***D***. (***C***) Normalization of ***D*** by a 1D distance-stratified standardization procedure. Its *d*-off diagonal part is normalized by *d*-off diagonal part-specific sample mean and sample standard deviation. (***D***) ***D*** is transformed by dividing 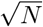. Under the null hypothesis, it is assumed to be a generalized Wigner matrix. (***E***) *P* value is calculated based on the fact that *θ*_*N*_, normalized largest eigenvalue of ***D***, follows Tracy-Widom distribution with *β* = 1 (denoted as *TW*_1_ distribution). A TAD is identified as a reorganized TAD if *P* value ≤0.05. (***F***) Reorganized TADs are classified into six subtypes based on changes in TAD boundaries. The heatmap diagram illustrates TADs in condition 1 (*Top*) and condition 2 (*Bottom*), with lines representing TAD regions of the same condition sharing the same color. (***G***) Example of the subtypes of reorganized TADs. Data are from two studies [33, 68]. *Top:* upper and lower triangular matrices represent Hi-C data in conditions 1 and 2, with blue triangles representing TADs and yellow triangles representing reorganized TADs; *Middle:* TAD regions from the same condition are represented by lines of the same color; *Bottom:* upper triangular section of the normalized difference matrix 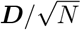 computated from the two Hi-C matrices in the *Top* section.

Note that, although each biological condition may have multiple Hi-C replicates, DiffDomain takes the combined Hi-C contact matrix from the replicates as the input which is a common practice to generate a large number of Hi-C interactions [33]. Correlations among Hi-C interactions lead to correlations among entries in the 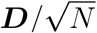 matrix, violating the independent assumption among upper diagonal entries of generalized Wigner matrix. However, the violation of the independence assumption does not substantially alter the properties of 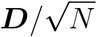 and DiffDomain based on empirical analysis results, suggesting that assumptions of DiffDomain are appropriate (see Methods section, Supplementary Results B.1, Supplementary Fig. S3). When *N* = 10, the Tracy-Widom distribution *TW*_1_ is an adequate approximation of the exact distribution of *θ*_*N*_ [41]. Under common 10 kb resolution Hi-C data, *N* = 10 refers to TADs with 100 kb in length, much smaller than the median TAD length 185 kb [33]. Thus DiffDomain only computes the *P* value for TADs with at least 10 chromosome bins, a practical constraint. DiffDomain is robust to a varied number of sequencing reads, Hi-C resolution, and different TAD callers (Supplementary Results B.2, Supplementary Fig. S4, S5).

### DiffDomain consistently outperforms alternative methods in multiple aspects

First, we assess the false positive rate (FPR) which is the ratio of the number of false positives to the number of true negatives. A smaller FPR means that the identified significantly reorganized TADs are more likely to be true. Due to the lack of gold-standard data, we resort to analyzing the proportions of significantly reorganized TADs between five different Hi-C replicates from the GM12878 cell line. These Hi-C replicates are generated by different experimental procedures and have a highly varied total number of Hi-C contacts (Supplementary Table S1). But the TADs are expected to have few structural changes between these Hi-C replicates. The proportion of identified reorganized TADs is treated as an estimate of FPR (Supplementary Methods A.2) and is expected to be small. The Hi-C resolution is chosen as 10 kb. Comparing GM12878 Hi-C replicates *combined* and *primary*, we find that DiffDomain, TADCompare [31], and HiCcompare [42] have FPRs that are smaller than the given significant level 0.05, suggesting good controls of FPR. In contrast, DiffGR [30], DiffTAD [29], and HiC-DC+ [38] have inflated FPRs (more than two-fold higher than 0.05), indicating poor controls of FPR (Fig 2A). Similar results are observed by repeating the above analysis to other GM12878 Hi-C replicates and 25 kb resolution Hi-C data (Supplementary Fig. S6).

**Figure 2:**
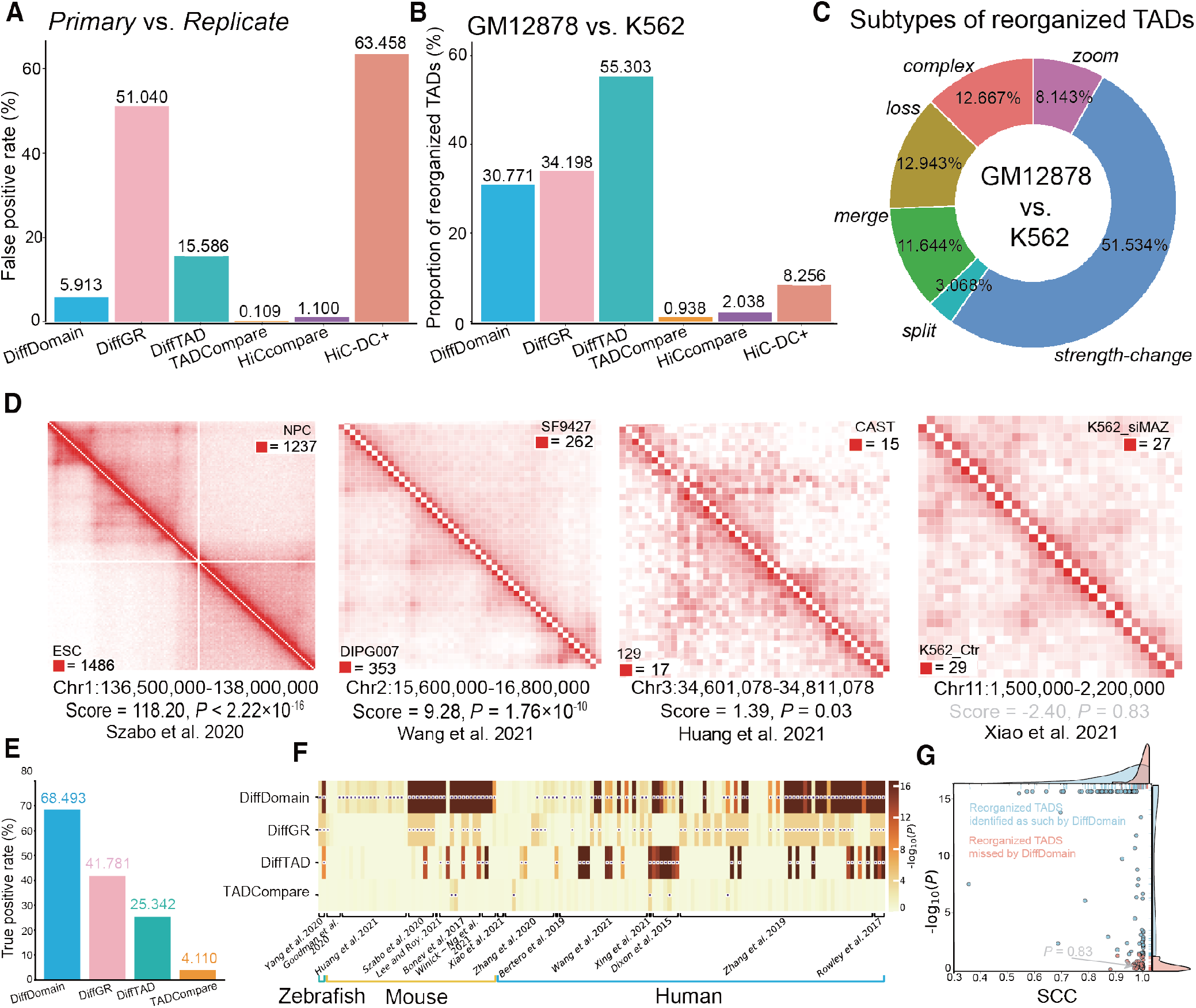
DiffDomain has better performance than alternative methods. (***A***) FPRs of DiffDomain and alternative methods in comparing two GM12878 Hi-C replicates (*combined* and *primary*). FPRs in comparing other Hi-C replicates of GM12878 are shown in Supplementary Fig. S6. FPR equals the ratio of the number of identified reorganized TADs to the number of TADs in GM12878. (***B***) Proportions of identified reorganized TADs by DiffDomain and alternative methods when comparing blood-related cell lines GM12878 and K562. TADs are GM12878 TADs. Results on other pairs of human cell lines are reported in Supplementary Fig. S7. (***C***) Percentages of the subtypes of reorganized TADs when comparing GM12878 and K562. Results on other pairs of human cell lines are reported in Supplementary Fig. S9. (***D***) Heatmaps showing Hi-C contact matrices of four truly reorganized TADs. Upper and lower triangular matrices represent Hi-C data in conditions 1 and 2. The scores and unadjusted *P* value below the heatmap are computed by DiffDomain Among them, three are correctly identified as such by DiffDomain (unadjusted *P ≤*0.05, true positives) and one is missed by DiffDomain (unadjusted *P >* 0.05, false negative). Truly reorganized TADs are manually collected and treated as the gold standard positives (see Supplementary Methods A.3 for more details). (***E***) Barplot showing TPRs of DiffDomain and alternative methods. TPR equals the ratio of the number of reorganized TADs that are identified as such (true positives) to the number of reorganized TADs (positives). (***F***) Heatmap showing unadjusted *P* values for testing truly reorganized TADs. (***G***) Scatter points of unadjusted *P* values by DiffDomain (*Y* -axis) and SCCs [43] (*X*-axis) when testing the truly reorganized TADs. *P* values smaller than 2.22*×*10^*−*16^ are substituted with 2.22*×*10^*−*16^.

Good control in FPR does not necessarily represent high power in detecting reorganized TADs between biological conditions. To investigate this, we compare TADs between two blood cell lines GM12878 and K562, DiffDomain identifies that 30.771% of GM12878 TADs are reorganized in K562. In contrast, TADCompare, HiCcompare, and HiC-DC+ only identify ≤8.256% of GM12878 TADs that are reorganized in K562 (Fig 2B), suggesting that they are too conservative to identify reorganized TADs between biological conditions. Similar results are observed by repeating the above analysis to other human cell lines and 25 Kb resolution Hi-C data (Supplementary Table S2, Supplementary Fig. S7), demonstrating the robustness of the observations. Conservation of TADCompare is because it is designed to scan every chromatin loci for potential reorganized TAD boundaries. But this analysis uses a given list of TADs, a common practice in Hi-C data analysis, which sharply narrows down the search space of TADCompare. Conservations of HiCcompare and HiC-DC+ are because they are designed for detecting differential chromatin interactions, not specifically tailored for identifying reorganized TADs.

Compared with TADsplimer which specifically identifies *split* and *merge* TADs [25], DiffDomain identifies similar numbers of *split* and *merge* TADs between multiple pairs of human cell lines (Supplementary Fig. S8). Importantly, DiffDomain identifies that the majority (minimum 43.137%, median 81.357%, maximum 98.022%) of the identified reorganized TADs are the other four subtypes (Supplementary Fig. S9), which can not be detected by TADsplimer. For example, among the GM12878 TADs that are identified as reorganized in K562 by DiffDomain, *strength-change* is the leading subtype of re-organized TADs, consistent with the fact that both GM12878 and K562 are blood cell lines (Fig 2C). These results demonstrate that DiffDomain has substantial improvements over TADsplimer.

We next investigate the true positive rate (TPR). A higher TPR means that more truly reorganized TADs are correctly identified as reorganized TADs. Through an extensive literature search, we collect 65 TADs that are reorganized between 146 pairs of biological conditions in 15 published papers (Supplementary Table S3, Supplementary Methods A.3). We use these TADs as the gold standard data to compute the TPR (Supplementary Methods A.2) and also call these TADs as truly reorganized TADs. Four truly reorganized TADs, either correctly identified or missed by DiffDomain, are shown in Fig. 2D. HiCcompare and HiC-DC+, designed for identifying differential chromatin interactions, are not directly applicable to only testing reorganization of one single TAD and thus are excluded from the analysis. We find that the TPR of DiffDomain is 68.493%, which is 1.639, 2.703, and 16.665 times higher than that of alternative methods DiffGR, DiffTAD, and TADCompare, respectively (Fig. 2E). Compared with DiffDomain, DiffGR, DiffTAD, and TADCompare only uniquely identify 11, 10, and 1 truly reorganized TADs, respectively (Supplementary Fig. S10). Closer examination shows that DiffDomain has much smaller *P* values than other methods (Wilcoxon rank-sum test, *P ≤*2.31 ×10^*−*8^, Fig. 2F), demonstrating that DiffDomain has stronger statistical evidence in favor of truly reorganized TADs. Based on the depictions of TAD changes reported in the publications, the truly reorganized TADs are broadly categorized into three groups: domain-level change, boundary-level change, and loop-level change (Supplementary Methods A.3). These groups have decreased reorganization levels with increased SCC scores between biological conditions (Supplementary Fig. S11A). Across the groups, DiffDomain consistently achieves the highest TPRs, while the second-best method varies (Supplementary Fig. S11B), further demonstrating the advantages of DiffDomain over alternative methods.DiffDomain still misses 31.507% of possible pairwise comparisons of truly reorganized TADs. One reason is that some of the missed truly reorganized TADs have highly similar Hi-C contact matrices between biological conditions. For example, the missed truly reorganized TAD, chr11:1500000-2200000 (Fig. 2D), has the stratum-adjusted correlation coefficient (SCC) [43] at 0.998. Generally, missed truly reorganized TADs have significantly (*P* = 8.26 × 10^*−*6^) higher SCC scores than those correctly identified reorganized TADs by DiffDomain (Fig. 2G). Similar results are observed when stratifying by the groups of truly reorganized TADs (Supplementary Fig. S11C). Because DiffGR uses SCC as the test statistic, these results also partially explain the low TPR (41.781%) of DiffGR and highlight that SCC alone is not optimal for identifying reorganized TADs. Another reason is that the resolutions of some Hi-C data are low since *P* values are moderately negatively associated with the maximum values of Hi-C contact matrices (Spearman’s rank correlation coefficient *ρ* = −0.534).

Additionally, DiffDomain is efficient in memory usage and acceptable in computation time compared with alternative methods (Supplementary Results B.3, Supplementary Fig. S12).

In summary, compared with alternative methods, DiffDomain has multiple improvements including FPRs, proportions of identified reorganized TADs between different biological conditions, subtypes of reorganized TADs, and TPRs.

### Reorganized TADs are associated with epigenomic changes

Armed with the advantages of DiffDomain over alternative methods, we explore the connections between TAD reorganization and epigenomic dynamics. We first showcase a GM12878 TAD that is significantly reorganized in K562 and classified as a *strength-change* TAD by DiffDomain (Fig. 3). The TAD covers a 445 kb region on chromosome 6. The TAD structural changes involve the vascular endothelial growth factor gene *VEGFA*, which is a major tumor angiogenic gene that is over-expressed in leukemia (See reviews [44, 45] for more details), consistent with the fact that K562 cells are chronic myelogenous leukemia cells. We find that the reorganized TAD has K562-specific functional annotations. The genomic region covered by the TAD is more accessible (1.71 times higher DNase peak coverage) in K562 than in GM12878 (Fig. 3B). The H3K27ac and H3K4me1 peak coverages of the TAD region in K562 are 3.24 and 3.28 times higher than the coverages in GM12878, respectively. In contrast, the H3K4me3 and H3K36me3 peak coverages of the TAD in K562 are only 1.31 and 1.25 times higher than the coverages in GM12878, respectively. Four regions in the TAD are annotated as super-enhancers [46] only in K562 (Fig. 3B). Note that, the TAD region is in A compartments in both cell types, suggesting that A/B compartments switch is not the reason for the gain in accessibility and histone modifications that are associated with gene activation. The normalized difference matrix ***D*** between Hi-C contact matrices of the TAD highlights that super-enhancer SE2 has increased Hi-C contacts with the *VEGFA* gene in K562 (Fig. 3C). To gain further insights into structural differences of the TAD in the two cell types, we compare the 3D structural representations of the TAD region. We run Chrom3D [47] 100 times to construct 100 possible 3D structures in each cell type for statistical comparisons. Two possible 3D structures with each per cell type illustrate the 3D structural differences of the TAD between GM12878 and K562 (Fig. 3D-E). Overall, the super-enhancer SE2, but not SE3, (Fig. 3B) is much spatially closer (*P<*2.22 ×10^*−*16^) to VEGFA in K562 than in GM12878 (Supplementary Fig. S13). These results show that the reorganized GM12878 TAD in K562 has K562-specific chromatin organization and potential biological functions.

**Figure 3:**
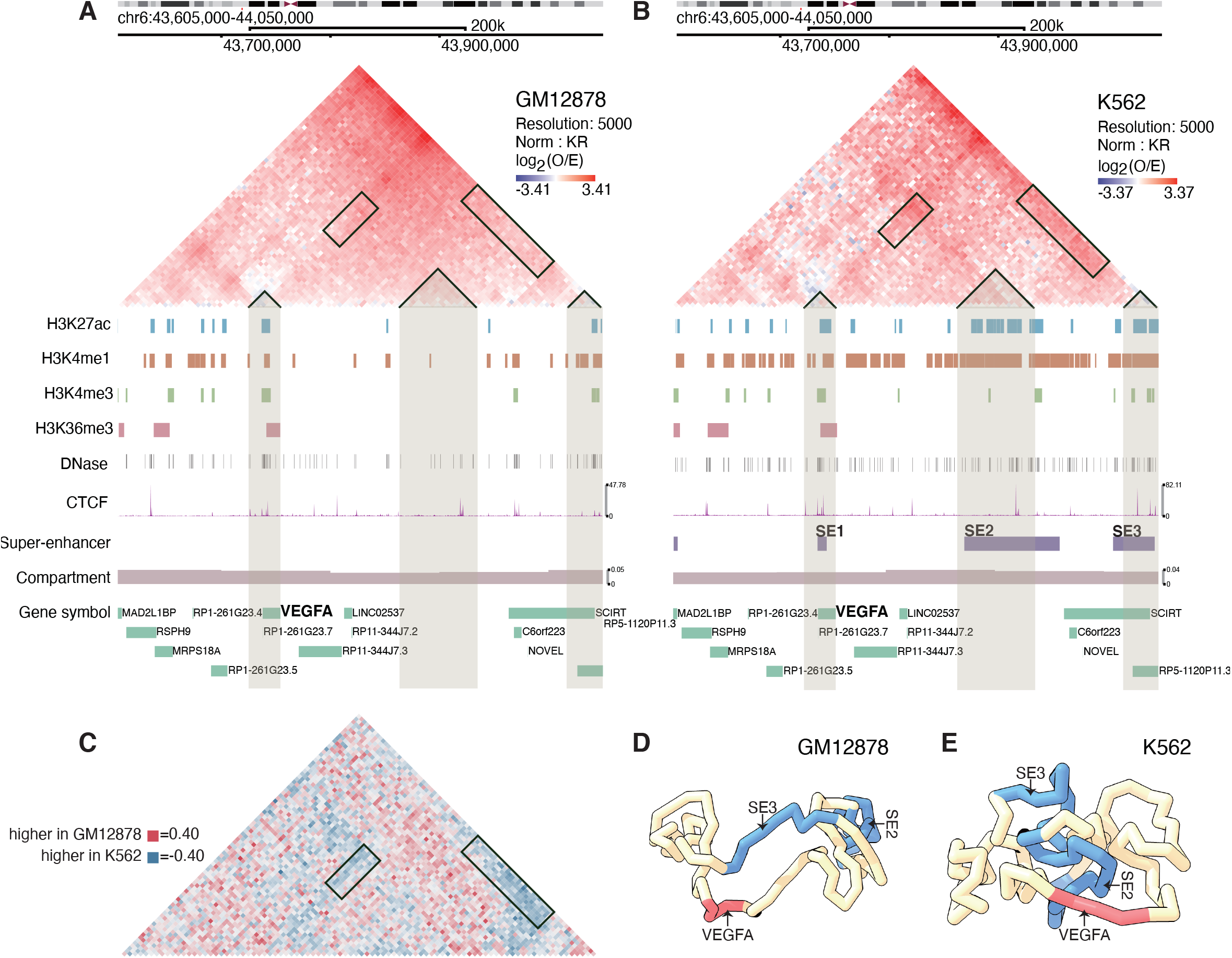
A reorganized TAD involves the angiogenic gene *VEGFA*. Hi-C contact matrices and 1D genomic tracks of the TAD region (Chr6:43605000-44050000) in GM12878 (***A***) and K562 (***B***) cell lines that show differences in both Hi-C data and epigenomics profiles involving *VEGFA* gene. (***C***) The normalized difference matrix ***D*** between the two cell lines highlights the differences in Hi-C contact maps. Rectangular boxes highlight the increased Hi-C interactions in K562. Rectangular boxes in panels (***A-B***) highlight corresponding sections in Hi-C contact maps. (***D-E***) Potential 3D structures of the TAD in GM12878 and K562. They are estimated by Chrom3D and demonstrate the TAD structural differences between GM12878 and K562.

Generally, comparative analyses across multiple pairs of human normal and disease cell lines reveal that *strength-change* reorganized TADs with increased contact frequencies have significant increase in the number of CTCF binding sites at TAD boundaries compared with other TADs, whereas lost TAD boundaries associated with *loss, zoom*, and *merge* TADs have significantly fewer number of CTCF binding sites, consisting with enrichment of CTCF binding sites in TAD boundaries (Supplementary Results B.4, Supplementary Fig. S14). Across diverse human cell lines, while TADs remain relatively stable, their proportions of reorganized TADs vary depending on the cell type, and these variations can cluster cell lines with similar cell identities (Supplementary Results B.5, Supplementary Fig. S7, S15). GM12878 (normal lymphoblastoid cell line) TADs that are reorganized in K562 (chronic myeloid leukemia cell lines) are enriched (*P* =0.01) in disease genes in chronic myelogenous leukemia. Across pairs of cell types, reorganized TADs tend to gain in chromatin accessibility and active transcription signals H3K27ac and K3K4me1. Particularly, TAD reorganization subtypes have distinct associations with chromatin accessibility as well as histone modifications. Specifically, TAD reorganization subtypes *strength-change-up, zoom, split*, and *complex* are associated with increased chromatin accessibility and histone modifications signals marking active transcription activities. Conversely, TAD reorganization subtypes *loss, strength-change-down* and *merge* are associated with decreased histone modifications signals marking active transcription activities, emphasizing the importance of TAD reorganization subtypes in investigating genome activity and functionality (Supplementary Results B.6, Supplementary Fig. S16, S17, S18). Compared to normal human astrocytes (NHA), patient-derived diffuse intrinsic pontine glioma cell lines DIPG007 and DIPGXIII share a substantial (73.46%) of reorganized TADs, some harboring potential oncogenes and super-enhancers, while dBET6 treatment demonstrates a stronger effects on TAD reorganization than BRD4 inhibition. (Supplementary Results B.7, Supplementary Fig. S19, S20, S21, Supplementary Table S4). Together, these results demonstrate the functional relevance of reorganized TADs in multiple human normal and disease cell lines.

### Reorganized TADs are enriched with structural variations (SVs)

SVs can contribute to diseases by rewiring 3D genome organization. To further demonstrate the biological relevance of reorganized TADs, we systematically investigate the associations between SVs and reorganized TADs. High-resolution SVs, including deletions and duplications, from erythroleukemia (K562 cell line) and pediatric high-grade glioma (DIPG007 and DIPGXIII cell lines) are downloaded from Wang et al. [48]. Because Arrowhead TADs does not necessarily cover the whole genome, SVs are filtered by keeping only those with their genomic regions overlapping with TADs (illustrated by two examples in Fig. 4A). The number of SVs and paired normal Hi-C data are summarized in Supplementary Table S5.

**Figure 4:**
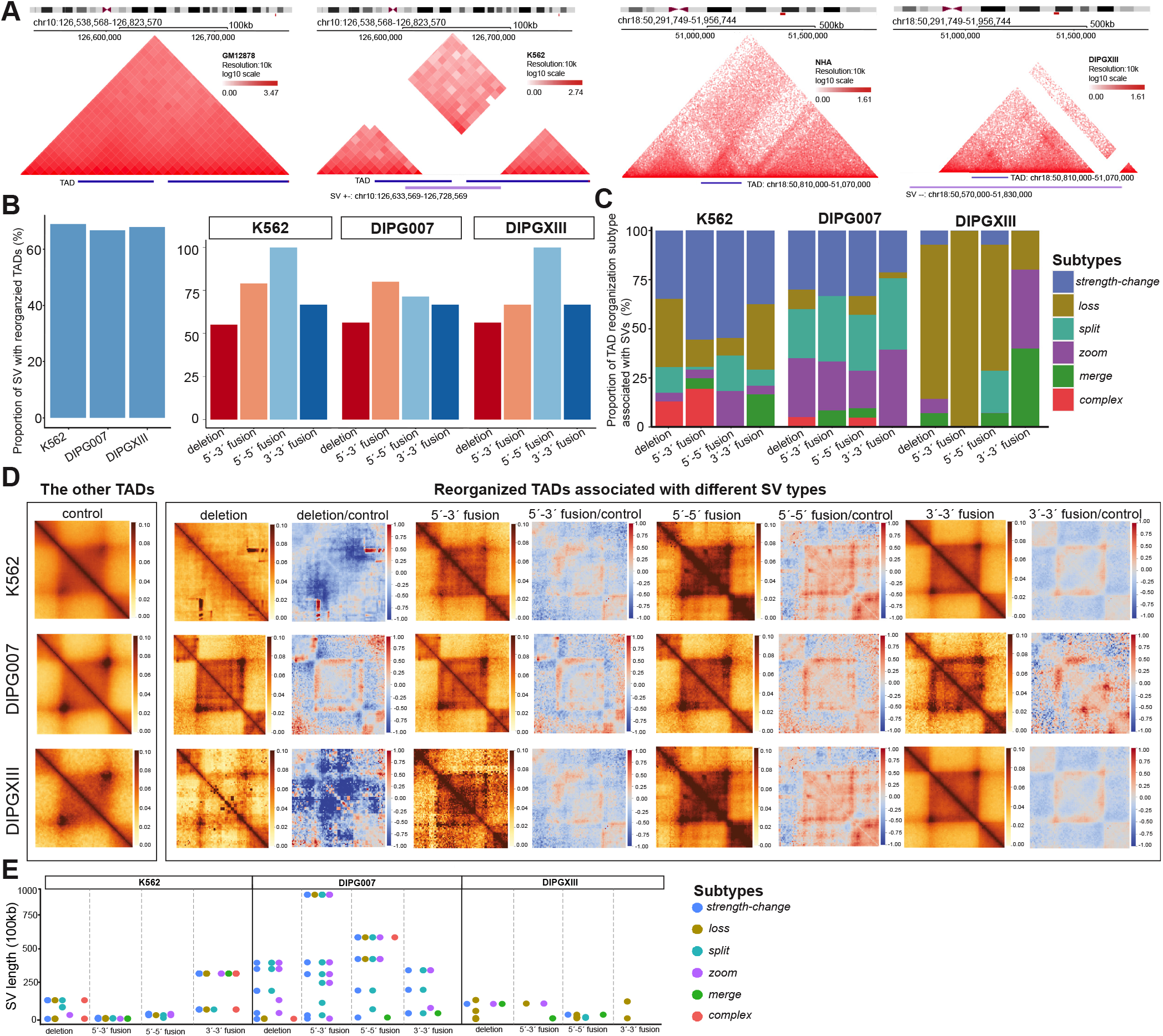
Associations between SVs and reorganized TADs in cell lines with disease. (***A***) Heatmaps showing two example SVs with associated reorganized TADS. *Left* two heatmaps showing the Hi-C contact maps from GM12878 and K562 for a specific K562 SV region, while *Right* two heatmaps showing the Hi-C contact maps from NHA and DIPGXIII for a specific DIPGXIII SV region. The first track below the heatmaps outlines the reorganized TAD regions, and the second track shows the SV regions. (***B***) Barplot showing the proportions of SVs with associated reorganized TADs across the three cell lines. *Left* barplot includes all SVs; *Right* barplots are stratified by the four types of SV, which include deletion and duplications such as 5^*1*^ to 3^*1*^ fusion, 5^*1*^ to 5^*1*^ fusion, 3^*1*^ to 3^*1*^ fusion. (***C***) Stacked barplot showing the proportions of reorganized TAD subtypes associated with each SV type. (***D***) APA plot summarizing the aggregated changes in reorganized TADs associated with each type of SV. *First column* APA plot summarizing the other TADs (not reorganized) using Hi-C data from condition 2 as a control. *Second column* APA plot summarizing reorganized TADs associated with deletion SVs using Hi-C data from condition 2. *Third column* APA plot showcasing log2-transformed fold-change of APA matrices, using APA matrices from the *second column* and the *first column*. Subsequent columns are APA plots summarizing reorganized TADs associated with 5^*1*^ to 3^*1*^ fusion (*fourth and fifth column*), 5^*1*^ to 5^*1*^ fusion, and 3^*1*^ to 3^*1*^ fusion, using Hi-C data from condition 2. Rows represent comparisons including GM12878 vs. K562, NHA vs. DIPG007, and NHA vs. DIPGXIII. (***E***) Jitterplot showing the length of SVs associated with each reorganized TAD subtype across the three cell lines. The APA matrices are based on 25 kb resolution Hi-C data produced by FAN-C using the command ‘fanc aggregate -m -p –pixels 90 -r -e –rescale’. The APA plots are generated using python function ‘sns.heatmap’. Abbreviations: ‘+-’, deletion;’-+’, 5^*1*^ to 3^*1*^ fusion; ‘--’, 5^*1*^ to 5^*1*^ fusion; ‘++’, 3^*1*^ to 3^*1*^ fusion. NHA, normal human astrocytes; DIPG007 and DIPGXIII, pediatric high-grade glioma cell lines.

If an SV region overlaps with one reorganized TAD, we consider the SV to have an associated re-organized TADs. We find that the majority (72.2%) of the SVs have such associations (Fig 4B), with proportions significantly higher than those of randomly sampled, equal-numbered reorganized TADs (Supplementary Fig. S22). Furthermore, the majority (*>*55%) of SVs with the same type also have such connections (Fig 4B). Each type of SVs has distinct associations with the subtypes of reorganized TADs. For example, in the comparison between GM12878 and K562 cell lines, the reorganized TADs associated with the four types of K562 SVs have differential distributions over their subtypes (Fig. 4C). The leading subtype of reorganized TADs, *strength-change*, is consistently observed across the four types of SVs. However, the second leading subtype of reorganized TADs varies among the four types of SVs (Fig. 4C). This observation is further emphasized by evident differences in the APA plots (Fig. 4D). Importantly, the association between SV type and TAD reorganization subtype is disease-specific, supported by the clear distinctions in both the APA plots and the subtype distributions of reorganized TADs across K562, DIPG007, and DIPGXIII cell lines (Fig. 4C-D). Particularly, leading reorganized TAD subtypes associated with SVs vary among cell types, with *strength-change* and *loss* in K562; *strength-change, zoom*, and *split* in DIPG007; and *loss* in DIPGXIII (Fig. 4C). This variability may be due to the substantial differences in SV lengths among these cell types (Fig. 4E). Notably, small proportions (17.4-27.7%, 8 in K562, 10 in DIPG007, and 4 in DIPGXIII) of SVs lack associated reorganized TADs (Fig. 4B). Upon visual examination through the Nucleome Browse, in total, 13 SVs have reorganized TADs that are not detected by DiffDomain, implying that the remaining 9 SVs in the three cell types may lack associated TAD reorganization (Supplementary Fig. S23-S25). Nevertheless, these results significantly enhance our understanding of the relationship between SVs and TADs compared to the previous study [48], further highlighting the biological relevance of reorganized TADs.

### DiffDomain improves profiling of TAD reorganization related to SARS-CoV-2 infection

Severe acute respiratory syndrome coronavirus 2 (SARS-CoV-2) causes over 640 million confirmed coronavirus disease 2019 (COVID-19) cases, including over 6.6 million deaths, worldwide as of December 2, 2022 (https://covid19.who.int), posing a huge burden to global public health. Wang et al. [49] is the first Hi-C study into the effects of SARS-CoV-2 infection on host 3D genome organization, finding a global pattern of TAD weakening after SARS-CoV-2 infection. However, the analysis uses aggregation domain analyses which cannot directly identify individual weakened TADs, in addition to missing other subtypes of TAD reorganization.

To further demonstrate the biological applications of DiffDomain, we reanalyze the data. We find that 20.58% (840 in 4082) mock-infected A549-ACE2 TADs are reorganized in SARS-CoV-2 infected A549-ACE2 cells. Among the reorganized TADs, *strength-change* TADs are the leading subtype (64.64%) (Fig. 5A) which is consistent with the global pattern of TAD weakening [49], verifying the reorganized TADs identified by DiffDomain. The following most frequent subtypes are *merge* TADs and *complex* TADs (18.45% and 10.95%, Fig. 5A), refining the characterization of TAD reorganization after SARS-CoV-2 infection. These reorganized TADs also enable refined profiling of transcriptional regulation in response to SARS-CoV-2 infection. Compared with the other TADs, the reorganized TADs have significantly higher numbers of up-regulated genes and down-regulated genes (*P ≤*1.27 ×10^*−*4^, Fig. 5B, Supplementary Fig. S26A). Similar significant patterns are observed comparing *strength-change* TADs and *split* TADs with the other TADs (*P ≤*8.07 ×10^*−*3^, Supplementary Fig. S26B), highlighting that the two subtypes have stronger connections with differentially expressed genes than other subtypes of reorganized TADs. In contrast, compared to the other TADs, the six subtypes of reorganized TADs have significantly higher numbers of both enhanced and weakened peaks of H3K27ac, SMC3, and RAD21 where H3K27ac is a marker for active enhancers and SMC3 and RAD21 are two critical cohesin subunits that regulate 3D genome organization (Supplementary Results B.8, Fig. 5C-E, Supplementary Fig. S26C-E). Gene-centric analysis shows that differentially expressed genes in reorganized TADs have stronger connections with differential chromatin interactions than in other TADs. In particular, the *strength-change* subtype has a 3-fold higher proportion (9.73%) of down-regulated genes with both enhanced and weakened chromatin interactions compared to other TADs (Supplementary Results B.9, Supplementary Fig. S27). These results suggest that, after SARS-CoV-2 infection, the subtypes of reorganized TADs all have strong associations with epigenome reprogramme, and *strength-change* TADs and *split* TADs have strong associations with deregulation of gene expression, highlighting the importance of subtypes of reorganized TADs identified by DiffDomain.

**Figure 5:**
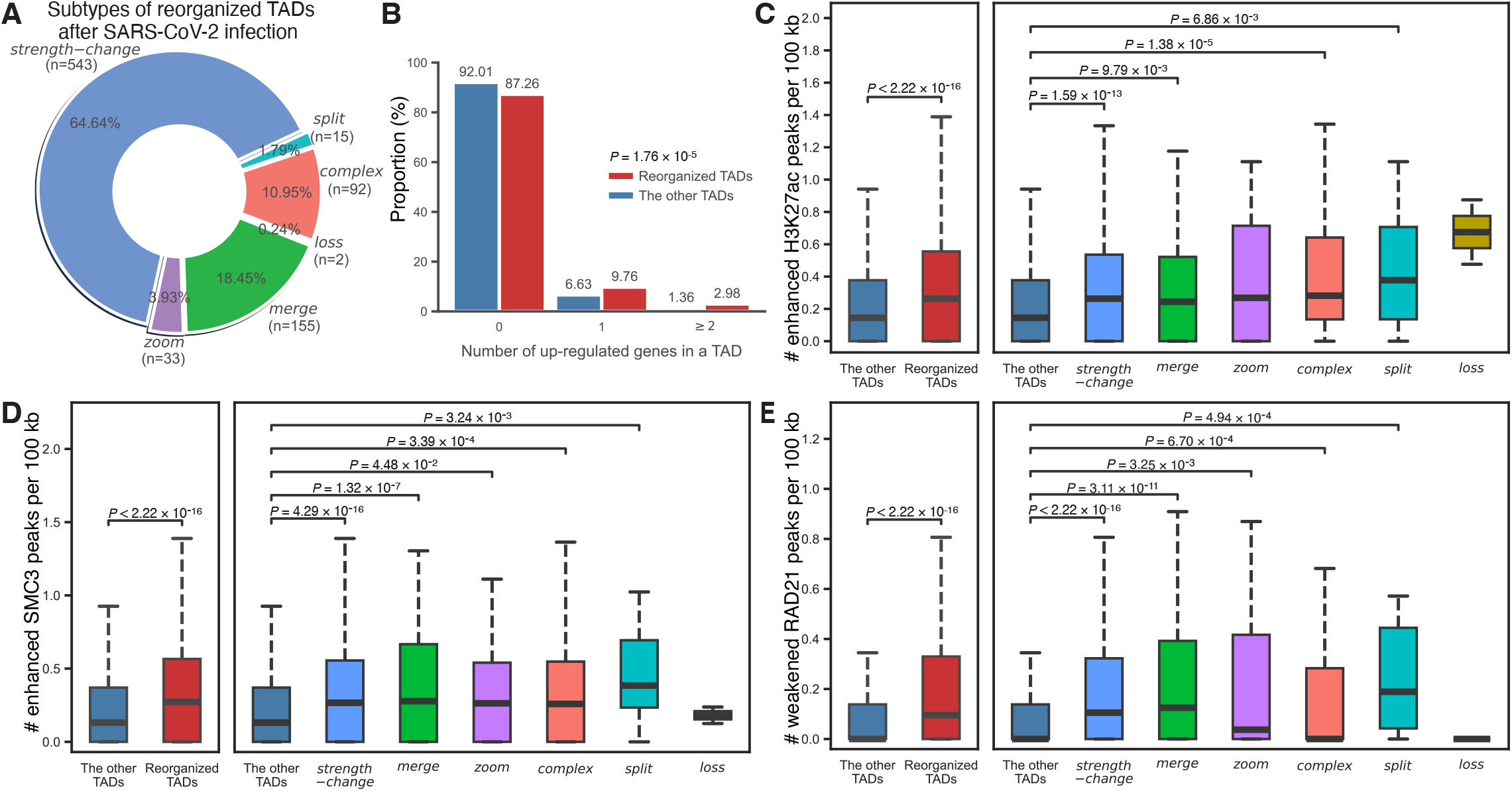
Mock-infected A549-ACE2 TADs that are reorganized in SARS-CoV-2 infected A549-ACE2. (***A***) Pie chart showing the percentages of subtypes of reorganized TADs in A549-ACE3 after SARS-CoV-2 infection. *Strength-change* TADs are the leading subtype. (***B***) Barplot comparing the number of up-regulated genes in the reorganized TADs and the other TADs. X-axis represents TADs categorized based on the number of up-regulated genes located within them, Y-axis represents the proportion of TADs. Boxplots comparing the numbers of enhanced H3K27ac peaks (***C***), enhanced SMC3 peaks (***D***), and weakened RAD21 peaks (***E***) per 100 kb. *Left* : comparing reorganized TADs with the other TADs (X-axis); *right* : comparison stratified by the subtypes of reorganized TADs (X-axis). Y-axis represents the number of differential peaks per 100 kb within TADs. *P* values are computed using one-sided MannWhitney U test, a nonparametric test dealing with asymmetric distributions. In the box plots, the middle line represents the median; the lower and upper lines correspond to the first and third quartiles; and the upper and lower whiskers extend to values no farther than 1.5IQR.

### DiffDomain characterizes cell-to-population and cell-to-cell variability of TADs using scHi-C data

Recent advances in scHi-C sequencing methods enable profiling of 3D genome organization in individual cells, revealing intrinsic cell-to-cell variability of TADs among individual cells. However, quantifying the variability is challenging due to the properties of scHi-C data such as high sparsity, low genome coverage, and heterogeneity [50, 51]. As a proof-of-concept, we apply DiffDomain to a moderate-sized scHi-C dataset from mouse neuronal development (median number of contacts per cell at 400,000) [52].

We first ask how many individual cells are sufficient to identify reorganized TADs between cell types with high reproducibility using pseudo-bulk Hi-C data (Supplementary Methods A.5.1). To do this, we design a sampling experiment to gradually increase the number of used individual cells and the reproducibility in identified reorganized TADs is quantified using the Jaccard index (Splementary Methods A.5.2). We find that DiffDomain can identity reorganized TADs between cell types with reasonable reproducibility (average Jaccard index ≥0.104) using as few as one hundred sampled cells (Fig. 6B-C). For example, DiffDomain consistently identifies that a neuronal TAD, harboring neuronal marker genes *GM24071, LRFN2, MOCS1*, and *1700008K24RIK* [52], is reorganized in oligodendrocytes with numbers of cells starting at 100 (Fig. 6A). Consistent of DiffDomain on other example genomic regions are shown in Supplementary Fig. S28. On average, DiffDomain identifies that 19.25% neuronal TADs are reorganized in oligodendrocytes using only 250 sampled cells from each cell type, consistent (average Jaccard index at 0.49) with the identified reorganized TADs using all available cells in both cell types (Fig. 6B). Similar results are observed when identifying neonatal neuron 1 (the youngest structure type) TADs that are reorganized in cortical L2-5 pyramidal cells (adult type) (Fig. 6C-D). Jointly increasing the numbers of sampled cells in both cell types improves the performance of DiffDomain, as expected (Fig. 6B). In contrast, only increasing the number of sampled cells in one cell type has a limited boost in performance. For example, oligodendrocytes have only 257 cells but neurons have 1380 cells. Further increasing the number of sampled neurons from 250 to 500 has a slight performance improvement (Fig. 6B). The observation is further confirmed when comparing neuronal subtypes neonatal neuron 1 and neonatal cortical L2-5 pyramidal cells, in which the number of sampled cells in the latter subtype is no more than 150 (Fig. 6C) or 228 (Fig. 6D). Repeating the analysis by using bulk Hi-C data [53, 54] to create gold-standard reorganized TADs, we observed similar patterns in neuronal TADs that are identified in astrocytes. For example, sampling 150 cells in each cell type identify 12.40% neuronal TADs that are reorganized in astrocytes on average. Among the reorganized TADs, 62.55% are also identified as reorganized TADs when bulk Hi-C data are used (Fig. 6E). Considering the medium number of contacts per cell at 400000, the merged Hi-C data from hundreds of cells are ultra-sparse pseudo-bulk Hi-C data. These results demonstrate that DiffDomain can work with ultra-sparse Hi-C data.

**Figure 6:**
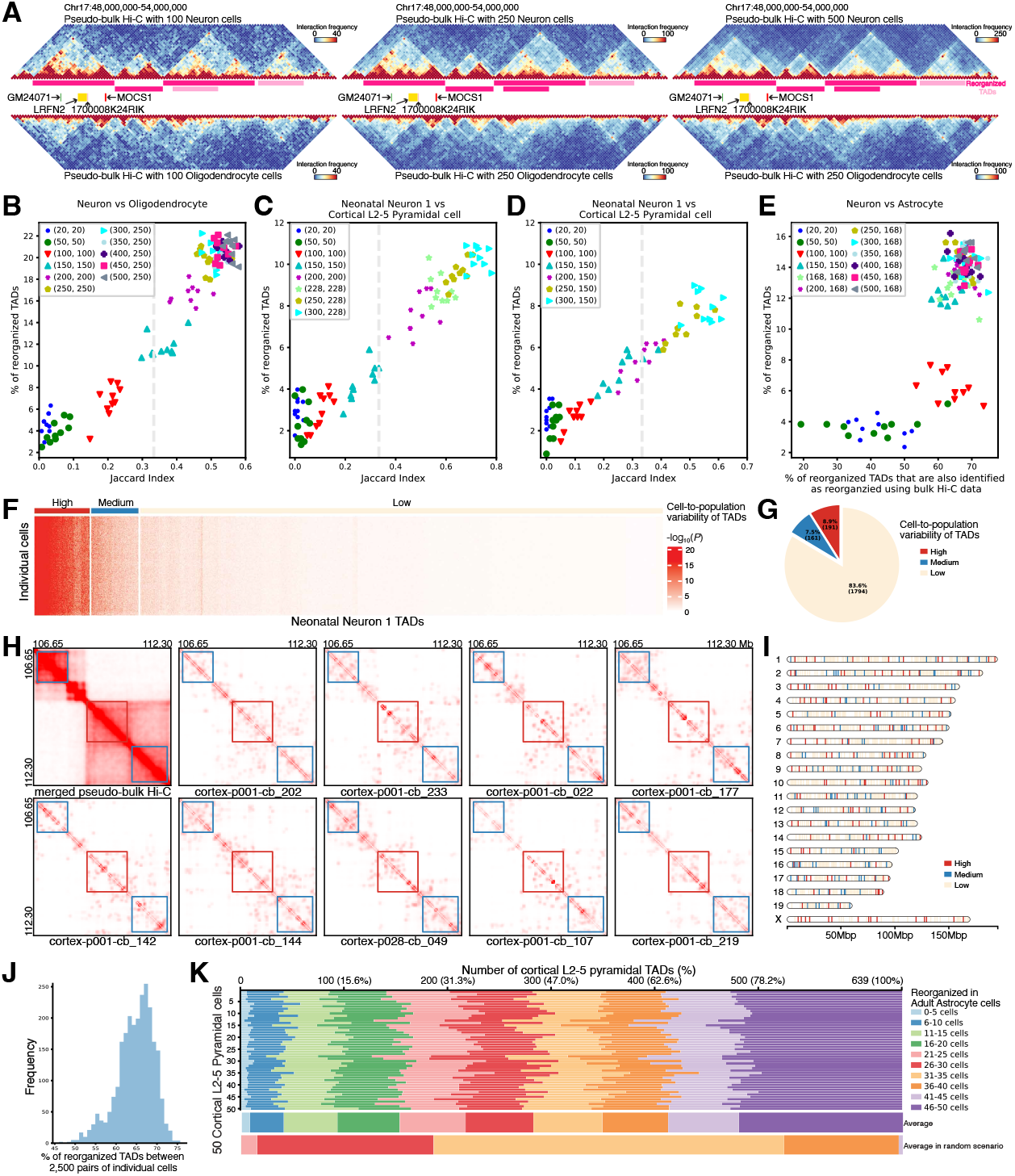
Application of DiffDomain on scHi-C data to identify reorganized TADs between cell types and characterize cell-to-population and cell-to-cell variability of TADs. (***A***) Visualization of the pseudo-bulk Hi-C contact maps and the identified neuronal TADs that are reorganized in oligodendrocytes (dark pink horizontal bars) using varied numbers of randomly sampled individual cells. Gene track shows four neuronal marker genes. (***B***) Scatter plot showing the proportions of neuronal TADs that are reorganized in oligodendrocytes using varied numbers (*k*_1_, *k*_2_) of randomly sampled individual cells from the two cell types. Their agreements with the set of reorganized TADs identified using all cells in each cell type are quantified by the Jaccard index (*X*-axis). The vertical dashed line is *JI* = 1*/*3, representing that two equal-sized sets share half of reorganized TADs. (***C-D***) Scatter plots showing the proportions of neonatal neuron 1 TADs that are reorganized in cortical L2-5 pyramidal cells. Up to 228 and 150 cortical L2-5 pyramidal cells are randomly sampled, respectively. (***E***) Scatter plot showing the agreements between sets of reorganized TADs that are identified using (1) pseudo-bulk Hi-C data and (2) bulk Hi-C data. (***F***) Heatmap showing high, medium, and low cell-to-population variational TADs. *P* value is from comparing scHi-C contact map of a TAD in an individual cell to the pseudo-bulk Hi-C contact map that represents population average. Classification of TADs is done by hierarchical clustering. (***G***) Pie chart showing the percentages of the high, medium, and low cell-to-population variational TADs. (***H***) Heatmaps visualizing one high cell-to-population variational TAD (middle red rectangular box) and two medium cell-to-population variational TADs (top-left and bottom-right blue rectangular boxes). (***I***) Chromosome map showing the genomic locations of high, medium, and low cell-to-population variational TADs. (***J***) Histogram showing the percentages of reorganized TADs in 2,500 pairwise comparisons of 50 cortical L2-5 pyramidal cells and 50 adult astrocytes. (***K***) Stacked bar graph showing the number (the percentage) of the cortical L2-5 pyramidal TADs that are reorganized in a varied number (0 to 50) of adult astrocytes. Y-axis represents the selected 50 cortical L2-5 pyramidal cells, indexed from 1 to 50. When comparing a specific cortical L2-5 pyramidal cell, such as cell 1, with 50 adult astrocytes, each cortical L2-5 pyramidal TAD is reorganized in a different number of adult astrocytes. X-axis (top of the plot) is the number (proportion) of the cortical L2-5 pyramidal TADs that are reorganized in 0 to 5 adult astrocytes, 6 to 10 adult astrocytes, and subsequent ranges (legend). Averaging the data in the stacked bar graph across the 50 cortical L2-5 pyramidal cells (Y-axis) results in the first stacked bar graph below, labeled as “Average” on the right. The subsequent stacked bar graph below is the average computed in random scenarios (Supplementary Fig. S30), labeled as “Average in random scenarios” on the right.

Next, we move to quantify the cell-to-population variability of TADs, that is, comparing TADs in individual cells to the population average. To do this, scHi-C data with 50 kb resolution from neonatal neuron 1 cells is imputed by scHiCluster [55] For each TAD, DiffDomain compares the imputed Hi-C contact map of the TAD in each cell to the pseudo-bulk Hi-C contact map. Resulted *P* values reflect cell-to-population variability of TADs and thus are used by hierarchical clustering to divide TADs into three categories: high, medium, and low cell-to-population variational TADs (Fig. 6F). We find that TADs have clear differential cell-to-population variability. One example high cell-to-population variational TADs and their adjacent medium cell-to-population variational TADs in 9 cells are shown in Fig. 6H. Among the 2146 neonatal neuron 1 TADs, 8.90% (191) are high cell-to-population variational TADs, 7.50% (161) and 83.60% (1794) are medium and low cell-to-population variational TADs (Fig. 6G). They are distributed across chromosomes (Fig. 6I). Similar results are observed in other cell types (Supplementary Fig. S29). These results demonstrate that TADs have clear differential variability between individual cells and the population average, consistent with earlier observations [26, 56].

Next, we move to investigate the cell-to-cell variability of TADs. Requiring only one Hi-C contact matrix from each condition, DiffDomain can directly quantify cell-to-cell variability of TADs between individual cells using imputed scHi-C data. Note that, similar to other methods, pairwise comparison of TADs using scHi-C data from thousands of individual cells leads to exponential growth in runtime and thus is computationally-expensive [50]. As a proof-of-concept, we apply DiffDomain to scHi-C data from randomly selected 50 cortical L2-5 pyramidal cells and 50 adult astrocytes. We find that the cell-to-cell variability of TADs is heterogeneous. The heterogeneity is consistent among the pairwise comparisons but quite different from those from random scenarios in which equal-numbered reorganized TADs are randomly assigned in pairwise comparisons. The proportion of reorganized TADs is consistent among the 2,500 pairs of individual cells, ranging from 46.0% to 75.7% (Fig 6J). Across the 50 cortical L2-5 pyramidal cells, the proportions of TADs that are reorganized in a varied number of adult astrocytes are fairly consistent (Fig. 6K). Moreover, the proportions of TADs that are either low in cell-to-cell variability (reorganized in no more than 10 adult astrocytes) or high in cell-to-cell variability (reorganized in more than 40 adult astrocytes) are much higher than those from random scenarios (Fig. 6K, Supplementary Fig. S30), consistent with differential cell-to-population variability of TADs as reported in the previous paragraph. This observation is also in concordance with the randomized placement of TAD-like blocks in individual cells but with a strong preference for TAD boundaries observed in bulk Hi-C data [18], further demonstrating the utilization of DiffDomain.

In summary, DiffDomain works on scHi-C data to identify reorganized TADs between cell types, identify TADs with differential cell-to-population variability, and characterize cell-to-cell variability of TADs.

## Conclusion and Discussion

In this work, we present a statistical method, DiffDomain, for comparative analysis of TADs using a pair of Hi-C datasets. Extensive evaluation using real Hi-C datasets demonstrates clear advantages of DiffDomain over alternative methods for controlling false positive rates and identifying truly reorganized TADs with much higher accuracy. Applications of DiffDomain to Hi-C datasets from different cell lines and disease states demonstrate that reorganized TADs are enriched with structural variations and associated with CTCF binding site changes and epigenomic changes, revealing their condition-specific biological relevance. By applying to a scHi-C dataset from mouse neuronal development, DiffDomain can identify reorganized TADs between cell types with considerable reproducibility using pseudo-bulk Hi-C data from as few as a hundred cells. Moreover, DiffDomain can reliably characterize the cell-to-population and cell-to-cell variability of TADs using scHi-C data.

The major methodological contribution of DiffDomain is directly characterizing the differences between Hi-C contact matrices using high-dimensional random matrix theory. First, DiffDomain makes no explicit assumption on the input chromatin contact matrices, directly applicable to both bulk and single-cell Hi-C data. Second, DiffDomain computes the largest eigenvalue *λ*_*N*_ of a properly normalized difference contact matrix ***D***, enabling the quantification of the differences of a TAD using all chromatin interactions within the TAD. Third, leveraging the asymptotic distribution of *λ*_*N*_, DiffDomain computes theoretical *P* values which is much faster in computation than simulation methods used in alternative methods. The model assumptions are realistic (Methods). Last but not the least, the normalized difference contact matrix ***D*** can help pinpoint genomic regions with increased or decreased chromatin interactions within the reorganized TAD, enabling model interpretation and refined integrative analysis with other genomic and epigenomic data.

There is room for improvement. First, DiffDomain has the highest accuracy in detecting truly re-organized TADs, but it misses some truly reorganized TADs that only show subtle structural changes (see example in Fig. 2D, G). Developing more powerful model-based methods is future work. The manually created list of gold-standard reorganized TADs is deposited in the GitHub repository that hosts the source code, which would benefit the research community for better method development. Second, because of the hierarchy of TADs and sub-TADs [2, 33], generalizations of DiffDomain to explicitly consider dependencies among TADs to further refine reorganized TADs identification and classification is future work. Third, it would be desirable to generalize DiffDomain to compare other TAD-like domains [57–62]. scHi-C data is imputed by scHiCluster [55] for the characterization of cell-to-population and cell-to-cell variability of TADs. Benchmarking the effects of different imputation algorithms, including Higashi [26, 63] and scVI-3D [64], on quantifying cell-to-population and cell-to-cell variability of TADs is future work.

As a subset of TADs, the identified reorganized TADs could be a critical unit for refined integrative analyses of multi-omics data. We demonstrate this type of application on different human cell lines and diseases states including SARS-CoV-2-infected A549-ACE2 cells. Future work integrating multiple types of omics data and functional perturbation experiments [65] is necessary to elucidate the causal relationships between TAD reorganization and disease.

DiffDomain is an interpretable statistical method for enhanced comparative analysis of TADs and it works for both bulk and single-cell Hi-C data. The accelerated application of Hi-C and scHi-C mapping technologies would generate ever-growing numbers of bulk and single-cell Hi-C data from different health and disease states. DiffDomain and its future generalizations would be an essential part of the Hi-C analysis toolkit for the emerging comparative analysis of TADs, which in turn would advance understanding of the genome’s structure-function relationship in health and disease.

## Methods

In this section, we first introduce the first part of DiffDomain: a model-based method to identify reorganized TADs. We state the model assumptions and their verification using real Hi-C data before reporting the second part of DiffDomain: classification of reorganized TADs into six subtypes. Last is missing value imputation.

### Model-based method to identify reorganized TADs

In this paper, our aim is to identify a subset of TADs that are reorganized between two biological conditions, such as a pair of healthy and disease cell lines/tissues. Specifically, given a set of TADs identified in one biological condition, we aim to identify the subset of TADs that are reorganized in another biological condition. To achieve this goal, we develop DiffDomain that takes a set of TADs and their Hi-C contact matrices as the input. The TADs are identified using the Arrowhead method [33] (Supplemental Methods A.4) and Hi-C contact matrices specific to each TAD region are extracted from the genome-wide KR-normalized Hi-C contact maps, unless specified otherwise. The core of DiffDomain is converting the comparison of Hi-C contact matrices into a hypothesis testing problem on their difference matrix. This difference matrix is modeled as a symmetric random matrix, enabling DiffDomain to borrow well-established theoretical results in high-dimensional random matrix theory.

Before explaining the hypothesis testing problem, we first introduce some mathematical notations and normalization operations. For each TAD in biological condition 1, let *N* denote the number of consecutive and equal-length chromosome bins within the genomic region covered by the TAD. Let 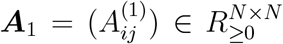 represent the symmetric KR-normalized Hi-C contact matrix, where 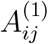 represents the non-negative Hi-C contact frequency between chromosome bins *i* and *j* (1≤ *i, ≤ j N*) in the TAD region in condition 1. In other words, ***A***_1_ serves as the Hi-C contact matrix specific to the TAD region in biological condition 1, forming a submatrix within the genome-wide Hi-C contact matrix. Similarly, 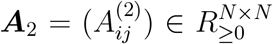 denotes the KR-normalized Hi-C contact matrix corresponding to the same TAD region, but in biological condition 2. It is well-known that the Hi-C contact frequency *A*_*ij*_ exponentially decreases with an increased linear distance between bins *i* and *j*. We first log-transform the Hi-C contact matrices ***A***_1_ and ***A***_2_ and compute their entry-wise differences, denoted by ***D***, as shown in Eq.(1).

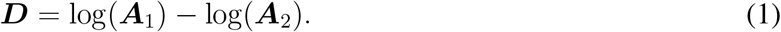

Values in Hi-C contact matrices ***A***_1_ and ***A***_2_ could have large differences because of variations in read depths. For example, the GM12878 Hi-C experiment has 4.76 times more Hi-C contacts than the K562 Hi-C experiment (Table S2). Among the 889 GM12878 TADs on Chromosome 1, the averages in the 889 ***D***s range from 1.479 to 2.611, with a median at 2.229. To adjust for the differences due to variations in read depths, we normalize 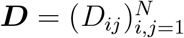 by standardizing each of its *k*-off diagonal blocks by

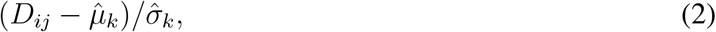

where *k* = *j − i*, 2 *− N ≤ k ≤ N −* 2,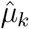 and 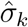 are the sample mean and standard deviation of (*D*_*mn*_)_1*≤m,n≤N,n−m*=*k*_. Here, without abuse of notations, we continue to use ***D*** to denote the resulted normalized difference matrix. Note that the normalization is TAD-specific because two different TADs most likely have different 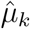 and 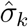, 2 *N ≤k ≤N −*2. Besides visualization in Fig. 1A-C, the effects of the above procedures for both bulk and single-cell Hi-C matrices from the same TAD are also visualized in Supplementary Fig. S1.

Intuitively, if a TAD does not undergo structural reorganization from biological condition 1 to biological condition 2, the differences between ***A***_1_ and ***A***_2_ are caused by multiple factors including variations in read depths and random perturbations of 3D genome organization. Thus, we assume that entries in ***D*** follow a standard Gaussian distribution, resulting in ***D*** being a symmetric random noise matrix with entries that follow a standard Gaussian distribution. Scaling ***D*** by 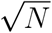, where *N* represents the number of bins in the TAD, results in 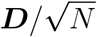 exhibiting characteristics typical of a well-studied random matrix known as a generalized Wigner matrix. With the justifications presented in the next subsection, we assume that 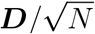 is a generalized Wigner matrix. The problem of identifying reorganized TADs is reformulated as the following hypothesis testing problem:

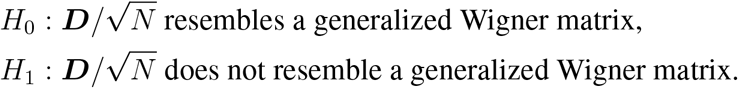

The largest eigenvalue of 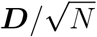, denoted by *λ*_N_, converges to 2 with increased *N*. This result helps us to reformulate the hypothesis testing problem as the following:

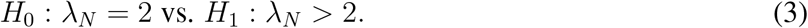

Under *H*_0_, *θ*_*N*_ = *N* ^2*/*3^(*λ*_*N*_ *−*2), a normalized *λ*_*N*_, converges in distribution to Tracy-Widom distribution with index *β* = 1, denoted as *TW*_1_.

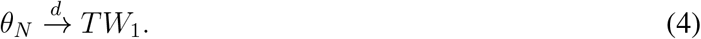

In other words, under *H*_0_, a TAD does not undergo structural reorganization in biological condition 2. Then the fluctuations of *θ*_*N*_ is governed by Tracy-Widom distribution *TW*_1_. Thus, we choose *θ*_*N*_ as the test statistic and compute *P* value by

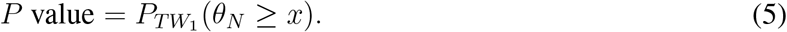

A smaller *P* value means that the TAD is more likely to be reorganized in condition 2.

For a set of TADs, *P* values are adjusted for multiple comparisons by a few methods with BH method as the default. The pseudocode of our DiffDomain algorithm is presented in Supplemental Methods A.1.

### Model assumptions and their verifications

Given two KR-normalized Hi-C contact matrices, DiffDomain computes the normalized difference matrix ***D***, bypassing complicated further normalization of individual Hi-C contact matrices [66]. Thus, DiffDomain makes no explicit assumptions on the individual Hi-C contact matrices. DiffDomain only makes assumptions on the normalized difference matrix ***D***. First, under *H*_0_, DiffDomain assumes that 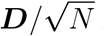 *N* is a generalized Wigner matrix: a symmetric random matrix with independent mean zero upper diagonal entries. Symmetry is satisfied by 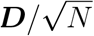 because Hi-C contact matrices are symmetric. The independence assumption on the upper diagonal entries is violated by 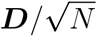 considering the well-known fact that Hi-C contact frequencies positively correlate with each other among nearby chromosome bins. However, the violation of the independence assumption does not substantially alter the properties of 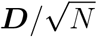 and DiffDomain (Supplementary Results B.1). Briefly, the empirical properties of 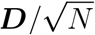 and DiffDomain resemble the following theoretical properties: (i) empirical spectral distribution of a generalized Wigner matrix converging to the well-established semicircle law, (ii) *λ*_*N*_ *→*2 (Supplementary Results B.1, Supplementary Fig. S3A-D), (iii) unadjusted *P* values following a uniform distribution when *H*_0_ is true and model assumptions are satisfied (Supplementary Results B.1, Supplementary Fig. S3E-F). The key result (4) requires one more assumption. It holds under the condition that the distributions of entries in generalized Wigner matrices have vanishing third-moments as *N* tends to infinity [67]. After the standardization procedure (2), 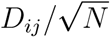 approximately follows a Gaussian distribution *N* (0, 1*/N*) whose third moment is 0, satisfying the vanishing third moment assumption. Taken together, assumptions of DiffDomain are appropriate.

### Reorganized TAD classification

Once a subset of reorganized TADs is identified, the classification of reorganized TADs is critical to interpret TAD reorganization and link them to the dynamics of genome functions. Motivated by classifications in previous studies [10, 30, 31], DiffDomain classifies reorganized TADs into six subtypes: *strength-change, loss, split, merge, zoom*, and *complex* (depending on the dynamics of TAD boundaries, Fig. 1F). Briefly, given a condition 1 TAD that is identified as reorganized in condition 2,

1. *Strength-change* represents that the boundaries of the TAD are the same in both conditions, but the Hi-C contact frequencies within the TAD either increased or decreased in biological condition 2, after proper normalization on the differences in total sequenced reads. Subsequently, a *strength-change* TAD can be classified into a *strength-change up* TAD or a *strength-change down* TAD. Before explaining the classification, a few mathematical notions are introduced. Given a matrix ***A***, Let *m*(***A***) be the medium value of the elements in ***A***, *s*(***A***) be the sum of the elements in ***A***. Let ***A***_1_ be the Hi-C contact matrix of the TAD in condition 1, ***A***_2_ the Hi-C contact matrix in condition 2. If a *strength-change* TAD satisfies 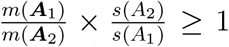, the *strength-change* TAD is classified as a *strength-change up* TAD. Otherwise, it is classified as a *strength-change down* TAD.
2. *Lost* represents that the genomic region covered by the TAD in condition 1 is not assigned to any TAD in condition 2.
3. *Split* represents that a TAD in condition 1 is split into at least two TADs in biological condition 2.
4. *Merge* represents that at least two TADs in condition 1 are merged into one TAD in condition 2. *Merge* is the opposite of *split* when condition 2 is treated as condition 1.
5. *Zoom* represents that the TAD in condition 1 intersects with exactly one TAD in condition 2, without being completely overlapping.
6. *Complex* represents the other reorganized TADs.

### Missing value imputation

Missing values may be exist in Hi-C contact matrices ***A***_1_ or ***A***_2_ for a specific TAD region, and their origin can vary. DiffDomain distinguishes between missing values caused by SVs and those caused by other factors such as low sequencing depth. When SVs are present, say in condition 2, DiffDomain first checks if a *m ×m* submatrix, *m ≥*3, of ***A***_2_ contains exclusively missing values. In such cases, the *m ×m* submatrix is imputed with a constant, with the default value of 1. Otherwise, if any row and column with a proportion of missing values greater than a given threshold, with default value of 0.5, DiffDomain removes the corresponding row/column from both ***A***_1_ and ***A***_2_. Subsequently, remaining missing values are imputed by the median contact frequency of interactions at the same distance within the corresponding contact matrix.

## Code availability

The software is published under the GNU GPL v3.0 license. The source code of DiffDomain is available at https://github.com/Tian-Dechao/diffDomain.

## Data availability

The study uses publicly available data from repositories including ENCODE portal, 4DN, and GEO. The accession codes for these datasets can be found in Supplementary Methods A.3 of the paper.

### Acknowledgement

This work was supported by the National Natural Science Foundation of China grant 12271536 (D.T.), National Key Research and Development Program of China grant 2021YFC2300102 (D.T.), GuangDong Basic and Applied Basic Research Foundation grant 2022A1515010043 (D.T.), Shenzhen Sustainable Research grant KCXFZ20211020172545006 (D.T.), National Natural Science Foundation of China grant 12171198 (Z.B.), and Jilin Provincial Foundation grant 20210101147JC (Z.B.). We thank Jiang Hu for helpful discussion on the theoretical properties of the proposed method, Jun Ding and Yang Zhang for help discussion that improved the manuscript, Jian Ma for helpful comments to improve the manuscript.

## Author contributions

Conceptualization: X.Z. and D.T.; Methodology: M.G. and D.T.; Software: M.G., D.H., X.Z., and D.T.; Investigation: D.H., M.G., X.Z, Y.D., H.X. and D.T.; Writing-Original Draft: D.T.; Writing-Review and Editing: L.Q., X.D., Z.B., X.Z. and D.T. Funding Acquisition: Z.B. and D.T.

## Supplemental Information

### A Supplementary Methods

#### A.1. Pseudocode of DiffDomain

The pseudocode of DiffDomain is presented in Algorithm 1.

##### Algorithm 1 Hypothesis-testing based method for identifying reorganized TADs.

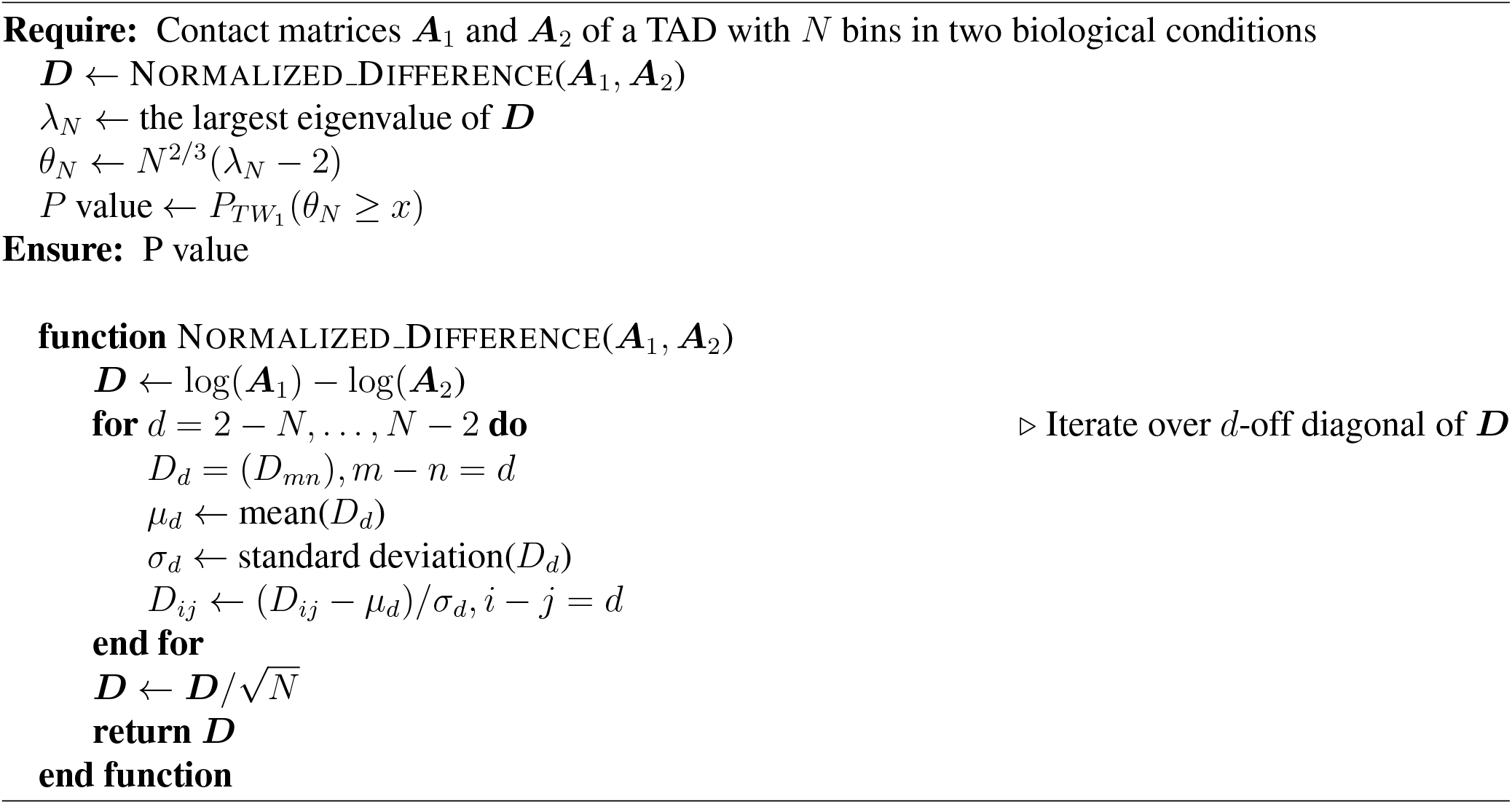

#### A.2. Method comparison

For a fair comparison, we choose alternative methods that calculate *P* -values: TADCompare [31], DiffGR [30], DiffTAD [29], TADsplimer [25], HiCcompare [42] and HiC-DC+ [38]. Note that, HiCcompare and HiC-DC+ are designed for detecting differential chromatin interactions. To run both methods, a pseudo chromatin interaction is created for each TAD where the TAD boundaries are the two loci and the sum of Hi-C contact map of the TAD is the Hi-C contact frequency of the pseudo chromatin interaction. We use false positive rate (FPR), true positive rate (TPR) and Jaccard Index (JI) for method evaluation.

FPR (Eq. 6) is used to quantify performance on controlling false positives (type I errors).

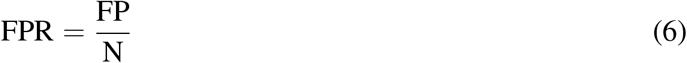

We assume biological replicates do not have reorganized TADs. FP stands for the number of identified reorganized TADs. N stands for the number of TADs. A lower FPR means a better control of false positives.

TPR (Eq. 7) is used to quantify performance on identifying truly reorganized TADs (power).

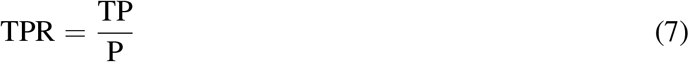

Here, TP is the number of identified reorganized TADs. P is the number of reorganized TADs in the manually collected gold standard data from a diverse set of publications. Note that a TAD might be counted multiple times if is is reorganized between multiple pairs of conditions. A higher TPR means a higher accuracy in identifying truly reorganized TADs. Combing FPR and TPR, a method with lower FPR and higher FPR is desirable.

JI, 0 ≤JI ≤1, is a metric quantifying the degree of overlap between two sets *A* and *B* as defined in Eq. (8).

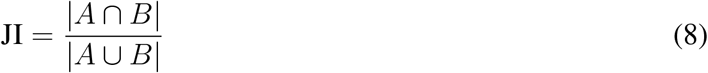

JI closer to 1 represents that two sets *A* and *B* share more elements. JI is used to compare the sets of identified reorganized TADs by DiffDomain with varied experimental settings. High values of JI indicate that DiffDomain is robust to experimental settings.

#### A.3. Collection of data used in this study

##### Bulk and scHi-C data

We use the following publicly available data: Hi-C of 8 human cell lines and multiple replicates of GM12878 cell lines from Rao et al. [33] (GEO:GSE63525); down-sampled Hi-C data of GM12878 from Xiong and Ma [69]; Hi-C of patient-derived DIPG, NHA and GBM cell line and DIPG frozen tissue specimens from Wang et al. [68] (GEO: GSE162976); Hi-C of two cell lines from Yang et al. [70] (GEO: GSE134055); Hi-C of two cell lines from Huang et al. [71] (4DN Data Portal: 4DNFIO21YDCV, 4DNFIUE5RAS6); Hi-C of two cell lines from Goodman et al. [72] (GEO: GSE138822); Hi-C of two cell lines in Szabo et al. [20], Lee and Roy [73], Bonev et al. [74] (4DN Data Portal: 4DNFI4OUMWZ8, 4DNFIHW8NTQX); Hi-C of two cell lines from Xiao et al. [75] (GEO: GSE143937); Hi-C of three cell lines from Zhang et al. [76] (GEO: GSE137376); Hi-C of two cell lines from Bertero et al. [77] (4DN Data Portal: 4DNFIN39NO4O, 4DNFINLW374M); Hi-C of seven cell types under different conditions from Wang et al. [68] (GEO: GSE162976); Hi-C of two cell lines from Xing et al. [78] and four cell lines from Rowley et al. [79] (GEO: GSE63525); Hi-C of H1 embryonic stem cells and H1 Mesenchymal stem cells from Dixon et al. [5] (GEO: GSE52457); Eight cell types under different conditions from Zhang et al. [80] (4DN Data Portal: 4DNFII6AN691, 4DNFINEQY95T, 4DNFIJ8JKKWJ); Hi-C of neuronal cell type from Jiang et al. [53] (SRR: SRR5617731, SRR5617733) and astrocytes from Sofueva et al. [54] (SRR: SRR941305). Hi-C of mock-infected and SARS-CoV-2 infected A549-ACE2 cells from Wang et al. [81].

Processed single-cell Dip-C of multiple cell types in mouse brains are from Tan et al. [52] (GEO: GSE162511).

##### Gold standard reorganized TADs

To create gold-standard reorganized TADs, a TAD is called as a reorganized TAD if it meets the following inclusion criteria: (1) The TAD is stated as having structural differences between a pair of conditions in a publication; (2) Chromatin contact map of the TAD is visualized in the main or supplementary figures; (3) Genome coordinate of the reorganized TAD can be explicitly inferred from the publication. In total, the manually collected gold standard data has 65 reorganized TADs that are reported in 15 publications (Supplementary Table S3). For each of such TAD, the Hi-C resolution is one of 5 kb, 10 kb, 25 Kb, 40 kb, 50kb, and 100 kb. Note that we do not choose a universal Hi-C resolution for these gold standard reorganized TADs for two reasons. First, those TADs have highly different lengths (minimum, median, and maximum lengths are 100 kb, 1.3 Mb, and 4.0 Mb, respectively). Second, the Hi-C data have different read depths.

The collected TADs are called as truly reorganized TADs in the main text. These TADs have a varied degree of reorganization and thus a varied difficulty in correctly identifying them. To further testing the performance of DiffDomain and alternative methods, we classify them into different groups. Note that, they are studied in a case-by-case manner and their adjacent TADs are not well described in the original publications. Thus, classifying them through DiffDomain is not suitable. Instead, they are classified according to their original definitions and are broadly categorized into three distinct groups: domain-level change, boundary-level change, and loop-level change. Domain-level change represents widespread changes in interactions within the TAD. Boundary-level change represents TAD reorganization due to changes in TAD boundaries, including boundary strength changes, boundary gain/loos, and boundary shifts. Loop-level change represents that TAD reorganization results from alterations in loops within the TAD, including loop gain/loss or increased/decreased loop contact frequencies. In total, there are 26 pairs of comparisons involving TADs with domain-level changes, 77 pairs involving TADs with boundary-level changes, and 43 pairs involving TADs with loop-level changes.

##### Epigenomic data for cell lines in Rao et al. [33]

We use the following publicly available data: RNA-seq in K562 from ENCODE[82] (ENCODE: ENCSR000CPZ); Super-enhancers of GM12878 and K562 from Hnisz et al. [46]; DNase peaks and histone modification profiles of multiple cell types from ENCODE[82];

##### Super-enhancers and oncogenes for NHA and DIPG cell lines

Super-enhancers in NHA, DIPG007, and DIPGXIII are called using ROSE algorithm [83] with input data downloaded from GEO database (GSE162976) [68]. A super-enhancer is linked to a reorganized TAD if their 1D distance is within 50 kb. Cancer genes are created by intersecting two gene lists: (1) cancer genes downloaded from OncoKB [84]; (2) Genes related to central nervous system that is downloaded from GeneCards with searching keyword [all] (glioma) OR [all] (Pediatric AND Brain AND Tumor) OR [all] (DIPG) OR [all] (central AND nervous AND system). The list of oncogenes is the subset of the above resulted cancer gene list, filtering by the column “Is Oncogene”. Oncogenes are then filtered using gene expression TPM *>* 1, resulting in 61 oncogenes.

#### A.4. TAD calling for bulk Hi-C data

One of the input of DiffDomain is the TAD list. By default, DiffDomain uses TADs called by Arrowhead [33], which can detect hierarchical TADs and are recommended by Zufferey et al. [85]. Arrowhead with a window size of 2000, resolution of 10 kb, and other parameters as the default is used to call TADs within Hi-C replicates from GM12878 [33]. For other cell lines in [33] that are used in this study, their TAD lists are downloaded from GEO database (GSE63525). In the case of NHA, DIPG007, and DIPGXIII cell lines, TADs are called by Arrowhead with a window size of 2000, resolution of 10 kb, and other parameters as the default. However, for the mock-infected A549-ACE2 and SARS-CoV-2 infected A549-ACE2 Hi-C data [49], Arrowhead is not used to call TADs. Instead, insulation score method is used for a better comparison with the original study’s findings [49]. More specifically, TADs are called using the command ‘run-insulator-score-caller.sh’, with a binsize of 10 kb, window size of 200 kb, and cutoff of 0.5 (https://github.com/4dn-dcic/docker-4dn-insulator-score-caller), following standard 4D Nucleome consortium protocol and utilizing the same parameters as the original study [49].

#### A.5. Application to scHi-C data

We briefly describe some methodological details when applying DiffDomain to scHi-C data. Hi-C resolution is chosen at 50 kb for this analysis.

##### A.5.1. Pseudo-bulk Hi-C data creation and TAD calling

For a given cell type, the pseudo-bulk Hi-C data is created by summing the scHi-C contact matrices from all individual cells with the same cell type. This procedure is used to (1) combine raw scHi-C contact matrices in identifying reorganized TADs between cell types, (2) combine imputed scHi-C contact matrices by scHiCluster in characterizing cell-to-population variability of TADs and cell-to-cell variability of TADs. Similar to TAD calling using bulk Hi-C data in other sections of this paper, once the pseudo-bulk Hi-C data is created, Arrowhead algorithm is applied to call TADs for the cell type. The TAD calling procedure is consistently used in identifying reorganized TADs between cell types, characterizing TADs with differential cell-to-population variability, and characterizing cell-to-cell variability of TADs.

##### A.5.2. Sampling individual cells to identify reorganized TADs between cell types

Given a cell type with *n* individual cells, *k* cells are randomly sampled and used to generate pseudo-bulk Hi-C data (details in Supplementary Methods A.5.1), *k* = 20, 50, …, *n*. The sampling procedure is repeated 10 times for each *k*. The pseudo-bulk Hi-C data with *n* cells are used in two ways. First, it is used to identify TADs by the Arrowhead algorithm. The resulting TAD list is used consistently for all *k* = 20, 50, …, *n*. Second, it is used to identify reorganized TADs between a pair of cell types, treated as the gold-standard reorganized TADs for the pair of cell types.

The reproducibility between the set of gold-standard reorganized TADs and the set of reorganized TADs using *k* sampled cells are quantified using the Jaccard index. A Jaccard index greater than 1*/*3 represents that two equal-sized sets share more than half of reorganized TADs.

#### A.6. Statistical analyses and Hi-C contact map visualization

Statistical analyses were conducted using the R version 3.5.1 and Python version 3.7.3. Hypergeometric test is used to test the enrichment of cancer genes in reorganized TADs. The Wilcoxon test is used to compare difference in means between two distributions such as gene expression, histone modifications and chromatin accessibility. Visualization of bulk and single-cell Hi-C contact matrices is done by Nucleome Browser [86] or in-house Python scripts.

### B Supplementary Results

#### B.1. Violation of independence assumption has mild effects on DiffDomain

After computing the normalized difference matrix ***D***, DiffDomain reformulates the problem of identifying reorganized TADs into the following hypothesis testing problem:

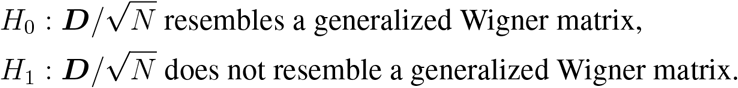

A generalized Wigner matrix is a symmetric random matrix with independent mean zero upper diagonal entries. The independence assumption on the upper diagonal entries is violated by 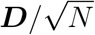 given that Hi-C contact frequencies positively correlate with each other among nearby chromosome bins. Note that DiffDomain does not assume independence among the entries in the individual Hi-C contact matrices.

To investigate the effects of violating the independence assumption on the performance of DiffDomain, we leverage three well-established theoretical results. The first result is the semicircle-law of the empirical spectral distribution of a generalized Wigner matrix ***A***_*N×N*_. Specifically, let *λ*_1_ ≤*λ*_2_ ≤· · · ≤ *λ*_*N*_ be the sorted eigenvalues of ***A***. The empirical spectral distribution of ***A*** is defined as

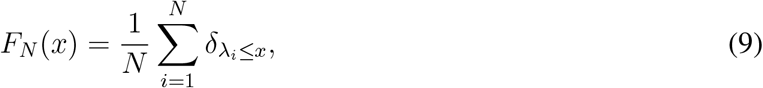

where *δ* is an indicator function. As *N→ ∞*, the discrete empirical spectral distribution *F*_*N*_ (*x*) converges to a continuous distribution function *F* (*x*) with the semicircular density function

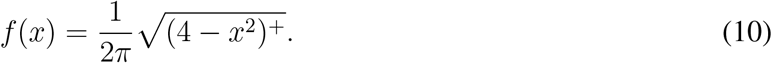

The second result is the largest eigenvalue *λ*_*N*_ of ***A*** converging to 2 in law as *N→ ∞*. The third result is the unadjusted *P* values following a uniform distribution when *H*_0_ is true and model assumptions are satisfied.

To verify the agreements between empirical properties of 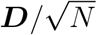 and theoretical properties of generalized Wigner matrices under *H*_0_ and their disagreements under *H*_1_, we choose two sets of real Hi-C data to generate multiple 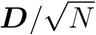 under *H*_0_ and *H*_1_, respectively. The first set consists of the 8261 GM12878 TADs and two GM12878 Hi-C replicates (*combined* and *primary* in Supplementary Table S1). The 8261 GM12878 TADs should not have much structure reorganization between the GM12878 Hi-C replicates. Thus, the corresponding normalized difference matrices 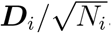, 1 *≤ i ≤* 8261, are treated as generated under *H*_0_. Second set consists of the manually collected 65 reorganized TADs between pairs of biological conditions from 15 published papers (Supplementary Table S3, Supplementary Methods A.3). Their normalized difference matrices 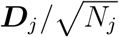, 1 ≤*j ≤*146, are treated as generated under *H*_1_. To verify the distribution of unadjusted *P* values, we use the *P* values from investigating TAD reorganization between multiple pairs of the GM12878 Hi-C replicates (Supplementary Table S1) as the *P* values under *H*_0_ and *P* values from comparing multiple pairs of human cell lines (Supplementary Table S2) as the *P* values under *H*_1_.

The analyses find broad agreements between empirical properties of 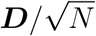 and theoretical properties of generalized Wigner matrices under *H*_0_ and substantial disagreements under *H*_1_. First, the estimated density functions of empirical spectral distributions of 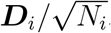, 1 *≤ i ≤* 8261, generated under *H*_0_ are close to the theoretical semicircular density function (Supplementary Fig. S3A). On the other hand, density functions of empirical spectral distributions of 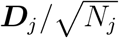, 1 ≤*j ≤*146, generated under *H*_1_ have much heavier tails on both sides and thus are different from the theoretical semicircular density function (Supplementary Fig. S3B). Secondly, only a small proportion (12.42%, 1026 out of 8261) of 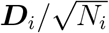, 1 ≤*i ≤*8261 generated under *H*_0_ have the largest eigenvalues that are greater than 2 (Supplementary Fig. S3C). In contrast, a large proportion (70.42%, 100 out of 142 possible comparisons) of 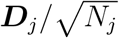, 1 ≤*j ≤*146, generated under *H*_1_ have the largest eigenvalues that are greater than 2 (Supplementary Fig. S3D). Thirdly, the distributions of unadjusted *P* values under *H*_0_ are approximately flattened over the interval (0, 0.6) (Supplementary Fig. S3E), resembling the expected uniform distribution. The distributions of unadjusted *P* values under *H*_1_ have peaks around 0 (Supplementary Fig. S3F), indicating the ability of DiffDomain in identifying reorganized TADs. The distributions of *P* values are much better than those from DiffGR and DiffTAD under *H*_0_ and much better than those HiCcompare and TADCompare under *H*_1_ (Supplementary Fig. S3E-F).

Taken together, these results demonstrate that the normalized difference matrices 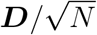 from real Hi-C data and generalized Wigner matrices have similar properties under *H*_0_ and different properties under *H*_1_, highlighting the violation of independence assumption having mild effects on DiffDomain. DiffDomain is appropriate for identifying reorganized TADs using real Hi-C data.

#### B.2. DiffDomain is robust

##### B.2.1. DiffDomain is robust to sequencing depth

To test the effect of sequencing depth on the performance of DiffDomain, we use down-sampled GM12878 datasets (using 2%, 5%, 10%, and 40% reads). First, we observed that lower the sequencing depth generally resulted in the reduced number of identified reorganized TADs. However, even at 5% reads, DiffDomain still identifies 530 GM12878 TADs that are reorganized in K562. At 40% reads, DiffDomain is able to recover 84.79% of reorganized TADs that are identified using 100% reads. More importantly, we observe that identified reorganized TADs are highly concordant across varied sequencing depths. For example, among the 530 reorganized TADs that are identified using 5% sequencing reads, 524 out of them are also identified as reorganized TADs using 10% sequencing reads (Supplementary Fig. S4A). Similar results are observed when comparing GM12878 data against down-sampled GM12878 data (Supplementary Fig. S4B).

##### B.2.2. DiffDomain is robust to Hi-C resolution

To test the effect of Hi-C resolution on the performance of DiffDomain, we apply DiffDomain to compare multiple pairs of human cell lines [33]. Hi-C resolution is chosen at 10 kb and 25 kb. We use two metrics to quantify the consistency of DiffDomain at the above two resolutions. The first metric is Jaccard index of two sets. One set is reorganized TADs identified with 10 kb resolution Hi-C data. Another set is reorganized TADs identified with 25 kb resolution Hi-C data. The second metric is the Pearson correlation coefficient. One vector is log-transformed BH-adjusted *P* values that are computed by DiffDomain with 10 kb resolution Hi-C data. Second vector is log-transformed BH-adjusted *P* values that are computed by DiffDomain with 25 kb resolution Hi-C data. Overall, Jaccard index and Pearson correlation coefficient are high across multiple pairs of cell types (Supplementary Fig. S4C-D), implying that DiffDomain is robust to a varied Hi-C resolution.

##### B.2.3. DiffDomain is robust to TAD callers

DiffDomain does not call TADs itself. It leverages multiple TAD callers developed by the community, facilitating commonly performed integrative analysis of Hi-C data and other genomics data. Among the multiple TAD callers, Arrowhead and TopDom are among the recommended TAD callers [85]. Arrowhead identifies hierarchical TADs, i.e., one TAD might contain multiple smaller TADs. In contrast, TopDom identifies non-hierarchical TADs [87]. Thus, Arrowhead and TopDom are chosen to test the robustness of DiffDomain to TAD callers.

Compared with Arrowhead, TopDom identifies a higher number of TADs and identified TopDom TADs cover a higher fraction of the genome across GM12878 replicates and different cell lines (Supplementary Fig. S5A-B). Regardless of the differences in TopDom TADs and Arrowhead TADs, DiffDomain controls the FPRs using both TopDom TADs and Arrowhead TADs (Supplementary Fig. S5C) where FPRs are estimated as the proportions of reorganized TADs between the Hi-C replicates from GM12878 cell line. Note that, when considering Hi-C replicates “GM12878 primary” and “GM12878 replicate” as condition 1, higher FPRs are observed using Arrowhead TADs than TopDom TADs. This is largely due to higher numbers of Arrowhead TADs than TopDom TADs in both replicates (Supplementary Fig. S5A). Applying to TopDom TADs and Arrowhead TADs in different human cell lines, DiffDomain identifies similar proportions of reorganized TADs between different human cell lines (Supplementary Fig. S5D). These results demonstrate that DiffDomain is robust to TAD callers.

Considering that the hierarchical organization of TADs is a critical feature, we choose the Arrowhead algorithm as the default method to call TADs in this paper.

#### B.3. Computation time and memory usage comparison

We compare DiffDomain with alternative methods in terms of computation time and memory usage. Condition 1 is chosen as GM12878 *combined* Hi-C data which has resolution up to 1 kb. Condition 2 is chosen as GM12878 *primary* Hi-C data, a replicate of GM12878 cell line. GM12878 *combined* Hi-C data has 9275 TADs that are identified by Arrowhead. For a fair comparison, computation time is estimated as CPU time (*user*) and memory usage is estimated as *memory resident set size*. Both statistics are estimated by Linux kernel function */usr/bin/time -f “%M %e”*. The experiment is repeated 10 times and the average is reported. We find that DiffDomain consistently uses the least memory across the varied Hi-C resolution compared with other methods (Supplementary Fig. S12B), demonstrating that DiffDomain is memory efficient. The reason is that DiffDomain iteratively compares Hi-C contact matrices of TADs, only loading Hi-C contact matrices for a given TAD into RAM. In contrast, other methods require loading Hi-C contact matrices of a whole chromosome into RAM. Memory efficiency does not make DiffDomain the slowest in computation time. DiffDomain completes the comparison of 9275 TADs within 1275 seconds (21.25 minutes) under commonly used Hi-C resolutions (10 kb, 25 kb, and 50 kb), which is much faster than DiffGR (Supplementary Fig. S12). Importantly, unlike other methods, DiffDomain has stable memory usage and computation time (Supplementary Fig. S12). Together, these results demonstrate that DiffDomain is memory efficient and reasonably fast.

#### B.4. Reorganized TADs are associated with changes in numbers of CTCF peaks

Earlier studies show that TAD boundaries are enriched with CTCF peaks [2]. To further demonstrate the biological relevance of reorganized TADs identified by DiffDomain, we compare the number of CTCF peaks in the boundaries of reorganized TADs and the other TADs. Regarding the reorganized IMR90 TADs in K562, reorganization of IMR90 TADs in K562 results in three types of TAD boundaries: lost, new, and stable. For simplicity, the K562 TAD boundaries that do not coincide with these TAD boundaries are called as the other TAD boundaries in K562 (Supplementary Fig. S14A). First, lost IMR90 TAD boundaries in K526 are due to *loss, zoom*, and *merge* subtypes of reorganized TADs. These TAD boundaries have significantly (*P ≤*2.22 ×10^*−*16^) fewer CTCF peaks than the other TAD boundaries in K562. Second, new K562 TAD boundaries are created in K562 due to *split* and *zoom* subtypes of reorganized TADs. These TAD boundaries have no significant difference in the number of CTCF peaks compared with the other TAD boundaries in K562. Thirdly, stable TAD boundaries are TAD boundaries of *strength-change* subtype of reorganized TADs. The subset of these TAD boundaries that are from *strength-change* subtype of reorganized TADs with increased contact frequencies (*strength-change up TADs*) have significantly (*P ≤*2.22 ×10^*−*16^) higher number of CTCF peaks than the other K562 TADs. In contrast, the subset of these TAD boundaries that are from *strength-change* subtype of reorganized TADs with decreased contact frequencies has no significant difference in the number of CTCF peaks compared with the other K562 TADs. Importantly, *complex* subtype of reorganized TADs have both lost and new TAD boundaries in K562. These TAD boundaries have no significant difference in the number of CTCF peaks compared with the other K562 TAD boundaries. Note that due to complexity and a small proportion of *complex* reorganized TADs, we do not stratify the comparison by lost and new types of TAD boundaries (Supplementary Fig. S14A). Repeating the analysis to other pairs of cell types find similar patterns (Supplementary Fig. S14B). These observations are consistent with the enrichment of CTCF peaks at TAD boundaries [2]. Additionally, a *merge* TAD represents two or more TADs of condition 1 are merged into one TAD in condition 2. In a *merge* TAD, the boundaries that are lost in condition 2 have fewer CTCF peaks in condition 2. The observation is consistent with the observation by Wang et al. [25], in which TAD splits and TAD merges between a pair of conditions are investigated. These results demonstrate that the reorganized TADs identified by DiffDomain are reasonable and biologically relevant.

#### B.5. Proportion of reorganized TADs is consistent with cell type identities

Armed with superior performance in terms of both FPR and TPR, we next apply DiffDomain to real data sets for biological relevance analyses. First, we investigate the TAD structural changes between different biological conditions by applying DiffDomain to 7 different human cell types (Supplementary Table S2). The proportions of reorganized TADs between cell types are reported in Supplementary Fig. S7. Overall, TADs are stable across cell types. Among the 42 pairwise combinations between the 7 cell types, the median proportion of significantly reorganized TADs is 13.76%. The observation is consistent with the previous study [2]. On the other hand, we do find that the proportion of reorganized TADs has a high variety and is cell type dependent. For example, the K562 TADs have the smallest proportion (11.41%) of reorganized TADs in KBM7 (both K562 and KBM7 are chronic myeloid leukemia). The highest proportion (41.24%) of reorganized K562 TADs is found in IMR90, a normal cell type. The ratio of the highest proportion to the smallest proportion is 3.61. Repeating the analysis to the other cell types, we find that the ratios are no less than 2.74. Similar results are observed at 25 kb resolution (Supplementary Fig. S7). Hierarchical clustering reveals that cell types with similar developmental relationships have smaller proportions of reorganized TADs (Supplementary Fig. S15), further demonstrating the utilization of DiffDomain.

#### B.6. Reorganized TADs are enriched in cancer genes and associated with epigenomic changes

First, we compare human hematopoietic cells: K562 (chronic myeloid leukemia cell lines) and GM12878 (normal lymphoblastoid cell line) cell lines. We downloaded 128 cancer genes that are related to chronic myelogenous leukemia from GeneCards [88]. GM12878 TADs that are reorganized in K562 are enriched (*P* =0.01, 66 out of the 128 cancer genes) with the cancer genes, indicating disease-trait relevance of reorganized TADs.

Second, we investigate the association between reorganized TADs and chromatin accessibility. Among the reorganized GM12878 TADs in K562, 16.25% of them have at least a 2-fold increase in DNase peak coverage in K562 cell type. Generally, the reorganized GM12878 TADs have significantly (*P <* 2.22 ×10^*−*16^) higher fold-change in DNase peak coverage in K562 than the other GM12878 TADs. The pattern for GM12878 TADs is consistently observed in the other cell types. Similar patterns are also observed for HUVEC and K562 TADs (Supplementary Fig. S16). These results suggest that the reorganized TADs tend to gain chromatin accessibility. Noticeably, we found reverse patterns for HMEC TADs. Compared to the other HMEC TADs, the reorganized HMEC TADs have significantly higher loss in DNase peaks in GM12878, HUVEC, and K562 cell types. Taken together, our results suggest that structural changes in TADs are associated with gain/loss of chromatin accessibility in the TAD regions across cell types.

We further ask the connection between TAD structural rewiring and histone modification dynamics in the TAD. It is known that histone modifications of H3K27ac and H3K4me1 are markers of active/poised enhancers and super-enhancers. H3K4me3 and H3K36me3 mark active/poised promoters and actively transcribed regions, respectively [89–91]. We find that reorganized TADs have significantly higher fold-changes in H3K27ac and H3K4me1/2 signals comparing with the other TADs. The significance patterns are much more profound than the patterns for H3K4me3 and H3K36me3 signals (Supplementary Fig. S17). In the subsets of TADs with gain (fold-change *>* 1) in chromatin accessibility or histone modifications signals, TAD reorganization subtypes *strength-change up, zoom, split*, and *complex* have significantly higher proportion of TADs with at least 2-fold gain in DNase peak coverage, H3K27ac, H3K4me1, and H3K4me3 signals (Supplementary Fig. S18A). In contrast, within the subsets of TADs with decrease (fold-change *<* 1) in chromatin accessibility or histone modifications signals, TAD reorganization subtypes *loss, strength-change down* and *merge* have significantly higher proportions of TADs with at least 2-fold loss in H3K27ac, H3K4me1, and H3K4me3 signals (Supplementary Fig. S18B). These results underscore distinct associations between TAD reorganization subtypes and chromatin accessibility as well as histone modifications. Specifically, TAD reorganization subtypes *strength-change up, zoom, split*, and *complex* are associated with increased chromatin accessibility and histone modifications signals marking active transcription activities. Conversely, TAD reorganization subtypes *loss, strength-change down* and *merge* are associated with decreased histone modifications signals marking active transcription activities, emphasizing the importance of TAD reorganization subtypes in investigating genome activity and functionality. The findings are consistent with previous study [91]. The authors discovered that histone modifications on promoters are stable across cell types. While the histone modifications on enhancers are cell type-specific and strongly associated with cell type-specific gene expressions. The findings are also consistent with the findings that *split* and *merge* subtypes of reorganized TADs are associated with changes in chromatin epigenetic state [25].

To summarize, comparative epigenomic analyses reveal that structurally reorganized TADs are enriched in cancer genes, and have associated changes in epigenomic profiles, suggesting the biological relevance of reorganized TADs.

#### B.7. Reorganized TADs are biologically relevant to diffuse intrinsic pontine glioma (DIPG)

DIPG is a pediatric high-grade glioma that is the leading cause of cancer death in children. A recent study generates the first high-resolution Hi-C data of DIPG cell lines and reveals that reorganized TADs are associated with alternations of transcriptional regulation [68]. However, the study uses a boundary-based method to identify reorganized TADs, missing reorganized TADs without boundary changes (*strength-change* TADs) and lacking statistical significance as mentioned in the Introduction section. To demonstrate the utilization of DiffDomain, we reanalyze the data.

We first focus on the NHA TAD that harbors the oncogenes MYCN. The TAD is identified as a *split* TAD in DIPG cell line DIPG007 and as a *zoom* TAD in DIPG cell line DIPGXIII (Supplementary Table S4), consistent with the previous observations that chromatin interactions in the TAD are strengthened in DIPG007 and DIPGXIII (Fig. 3A in [68]). Genome-wide, DiffDomain identifies that 14.89% (352) and 15.78% (373) of NHA TADs are reorganized in DIPG007 and DIPGXIII, respectively. The two sets of reorganized TADs share more than 73.46% of reorganized TADs (Supplementary Fig. S19A), consistent with the fact that both DIPG007 and DIPGXIII are patient-derived cell lines. Among the reorganized TADs, 26.99% and 26.27% are *strength-change* TADs (Supplementary Fig. S20A,C), refining the TAD reorganization analysis as reported in Wang et al. [68]. Although these reorganized TADs are not significantly enriched in oncogenes in DIPG007 and DIPGXIII, multiple subtypes of reorganized TADs including *strength-change* TADs, *split* TADs, and *merge* TADs, harbor at least one oncogenes (Supplementary Table S4). Four examples are visualized in Supplementary Fig. S21. Among the subsets of oncogenes (Supplementary Table S4), *MYCN, CCND1* genes are well-known functional genes in DIPG; *SOX2, VEGFA* and other oncogenes are under-explored in DIPG, which may have therapeutic implications for DIPG. Stratifying the results by super-enhancers, two reorganized TADs harbor both oncogenes and DIPG007 super-enhancers (Supplementary Methods A.3), two reorganized TADs harbor both onco-genes and DIPGXIII super-enhancers (see Supplementary Fig. S21C-D for two examples), expanding the analysis from enhancers [68] to super-enhancers. Additionally, a higher number of DIPG007-DMSO TADs are reorganized after the BRD degrader treatment (dBET6) compared with BET BRD inhibitor treatment (BRD4i) (41 in dBET6 vs. 9 in BRD4i, 5 are shared, 10 kb Hi-C resolution, Supplementary Fig. S19D). The preliminary results suggest stronger effects of dBET6 on TADs, consistent with dBET6 having stronger effects on chromatin loops and A/B compartments [68]. Hi-C resolution at 10 kb and 25 kb provide similar results (Supplementary Fig. S19B-C, S20B, D). These results further demonstrate the utilization of DiffDomain and the functional relevance of identified reorganized TADs.

#### B.8. Reorganized TADs are associated with epigenome reprograme after SARS-CoV-2 infection

Regarding epigenome reprogramme, the reorganized TADs have significantly (*P<* 2.22 ×10^*−*16^) higher numbers of enhanced and weakened peaks of H3K27ac (Fig. 5C, Supplementary Fig. S26C), a marker for active enhancers. The reorganized TADs have significantly (*P<* 2.22 ×10^*−*16^) higher numbers of enhanced and weakened peaks of SMC3 (Fig. 5D, Supplementary Fig. S26D), a cohesin subunit that regulates 3D genome organization. Similar patterns are observed for another key cohesin subunit RAD21 (Fig. 5E, Supplementary Fig. S26D), but to a lesser degree than SMC3 because a much smaller number of differential peaks are called by MAnorm2. Stratifying the above comparative analysis by the subtypes of reorganized TADs, similar patterns are observed (Fig. 5C-E, Supplementary Fig. S26C-E), demonstrating that enrichments of differential peaks of H3K27ac, SMC3, and RAD21 have no strong preference over the subtypes of reorganized TADs.

#### B.9. Distinct associations of differentially expressed genes with differential chromatin interactions in reorganized TADs after SARS-CoV-2 infection

To explore the connection between differentially expressed genes and their potential differential Hi-C interactions, we use HiC-DC+ to identify differential chromatin interactions. HiC-DC+ is run using Hi-C resolution of 10 kb, ssiz of 0.1, and other parameters with default values. Given the relatively low resolution of Hi-C data, only a limited number of significantly differential chromatin interactions are identified with an adjusted *P* value threshold of 0.05, Thus, the original *P* value with threshold of 0.05 is used to identify significantly differential chromatin interactions. First, we study down-regulated genes. A down-regulated gene is called as having a differential chromatin interaction if its transcription start site is located within 50 kb of one of the chromosome bins involved in the differential chromatin interaction. We find that, among the down-regulated genes located in reorganized TADs, 36.9% (62 out of 168) have at least one differential chromatin interactions (Supplementary Fig. S27A). The proportion is 1.74 times higher than that of the down-regulated genes located in the other TADs (21.2%, 92 out of 433, Supplementary Fig. S27A). Due to low number of differential chromatin interactions in down-regulated genes (Supplementary Fig. S27B-C), down-regulate genes are classified into four groups depending on involved differential chromatin interactions: with only enhanced chromatin interaction, with only weakened chromatin interaction, with both enhanced and weakened chromatin interaction, and without differential chromatin interaction. Stratified by the group of down-regulated genes and subtype of reorganized TADs, comparative analysis reveals that down-regulated genes located in *strength-change* TADs have a 3 times higher proportion (9.73%) of down-regulated genes with both enhanced and weakened chromatin interactions compared with down-regulated genes located in the other TADs (Supplementary Fig. S27F). Down-regulated genes in the other subtypes of reorganized TADs also have distinct patterns involving differential chromatin interactions compared to down-regulated genes in the other TADs (Supplementary Fig. S27F). Similar observations are observed when analyzing up-regulated genes (Supplementary Fig. S27A,D-E,G). These observations underscore the role of TAD reorganization in gene expression regulation.

### C Supplementary Tables

**Table S1:** Summary of the multiple Hi-C replicates from GM12878 cell line [33]. The replicates are produced with different settings and have different total Hi-C contacts. The highest total Hi-C contacts (combined) is 18.9 fold higher than the lowest total Hi-C contacts (noXlink). The data are used to quantify the false positive rates of DiffDomain and alternative methods.

| Abbreviation | Restriction enzyme | Crosslinking | # contacts |
| --- | --- | --- | --- |
| Primary | MboI | Yes | 3,587,190,419 |
| Replicate | MboI | Yes | 2,937,330,058 |
| Combined | MboI | Yes | 6,524,520,477 |
| DpnII | DpnII | Yes | 567,550,645 |
| noXlink | MboI | No | 345,058,337 |

**Table S2:** Brief description of the seven cell types [33] used in the study. Two of them are cancer cell lines.

| Cell line | Disease | Cell type | # contacts |
| --- | --- | --- | --- |
| GM12878 | Normal | B-lymphoblastoids | 6,524,520,477 |
| HMEC | Normal | Mammary Epithelial | 538,392,678 |
| HUVEC | Normal | Umbilical Vein Endothelial | 732,108,988 |
| IMR90 | Normal | Lung Fibroblasts | 1,535,222,082 |
| NHEK | Normal | Epidermal Keratinocytes | 1,073,207,572 |
| K562 | Cancer | Erythroleukemia | 1,366,228,845 |
| KBM7 | Cancer | Near Haploid Human Myelogenous Leukemia | 1,247,936,408 |

**Table S3:**
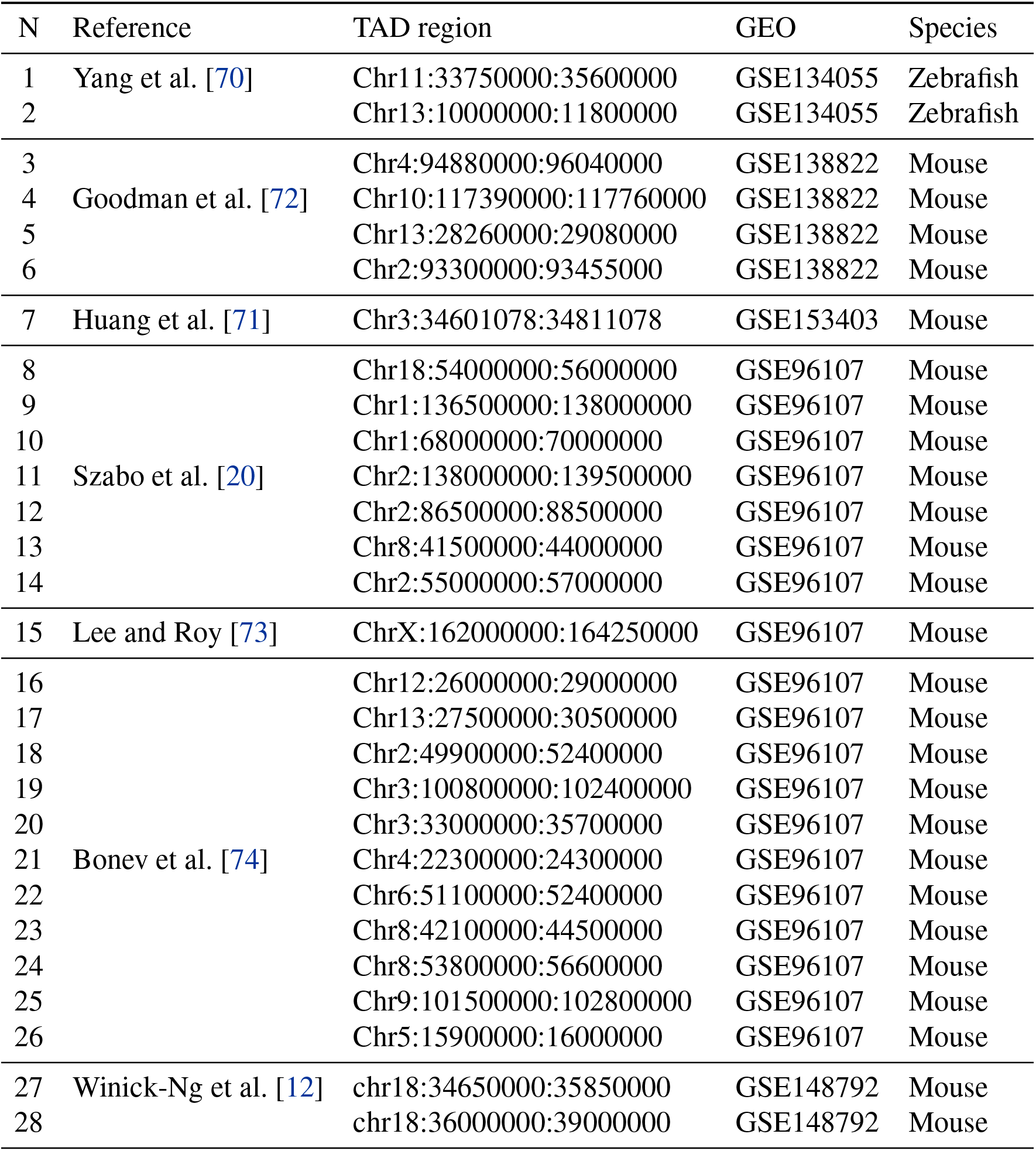
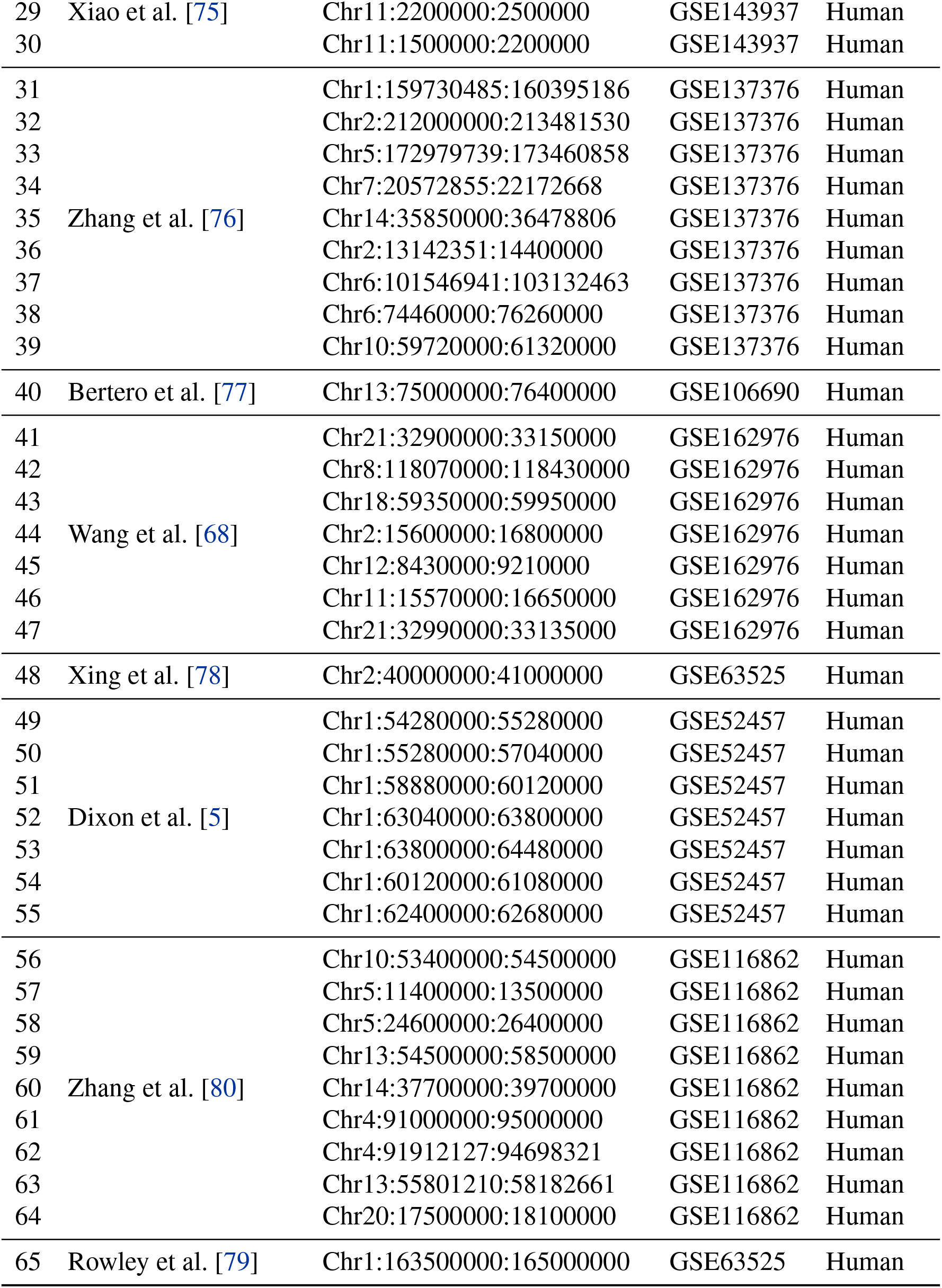
Gold-standard reorganized TADs that are manually collected from multiple studies.

**Table S4:** Oncogenes in the reorganized TADs when comparing NHA with DIPG cell lines (DIPG007 and DIPGXIII). Abbreviation: NHA, normal human astrocytes; DIPG, pediatric high-grade glioma.

| Comparison | Reorganized TAD | Subtype of reorganization | Oncogene |
| --- | --- | --- | --- |
| NHA vs. DIPG007 | chr2:15640000-16580000 | *split* | *MYCN* |
|  | chr3:18170000-182690000 | *split* | *SOX2* |
|  | chr3:189250000-189950000 | *strength-change* | *TP63* |
|  | chr6:43760000-44080000 | *loss* | *VEGFA* |
|  | chr8:127710000-129770000 | *zoom* | *MYC* |
|  | chr11:69100000-69660000 | *loss* | *CCND1* |
| NHA vs. DIPGXIII | chr2:15640000-16580000 | *zoom* | *MYCN* |
|  | chr3:181700000-182690000 | *zoom* | *SOX2* |
|  | chr3:189250000-189950000 | *strength-change* | *TP63* |
|  | chr6:43630000-44080000 | *loss* | *VEGFA* |
|  | chr7:54760000-55250000 | *strength-change* | *EGFR* |
|  | chr8:127710000-129770000 | *split* | *MYC* |
|  | chr10:120940000-121720000 | *strength-change* | *FGFR2* |
|  | chr11:69100000-69660000 | *merge* | *CCND1* |
|  | chr12:102100000-102770000 | *strength-change* | *IGF1* |

**Table S5:** Summary of SVs and Hi-C data used for analyzing the associations between SVs and reorganized TADs identified by DiffDomain. SVs detected by the deep learning method EagleC using bulk and scHi-C data are downloaded from Wang et al. [48]. Condition 1 contains GM12878 and NHA cell lines, representing health conditions. Condition 2 contains K562, DIPG007, and DIPGXIII cell lines, representing disease conditions.

| N | Condition 1 | Condition 2 | # SVs associated with condition 1 TADs | Hi-C data (Ref.) |
| --- | --- | --- | --- | --- |
| 1 | GM12878 | K562 | 39 | [33] |
| 2 | NHA | DIPG007 | 36 | [68] |
| 3 | NHA | DIPGVIII | 23 | [68] |

### D Supplementary Figures

**Figure S1:**
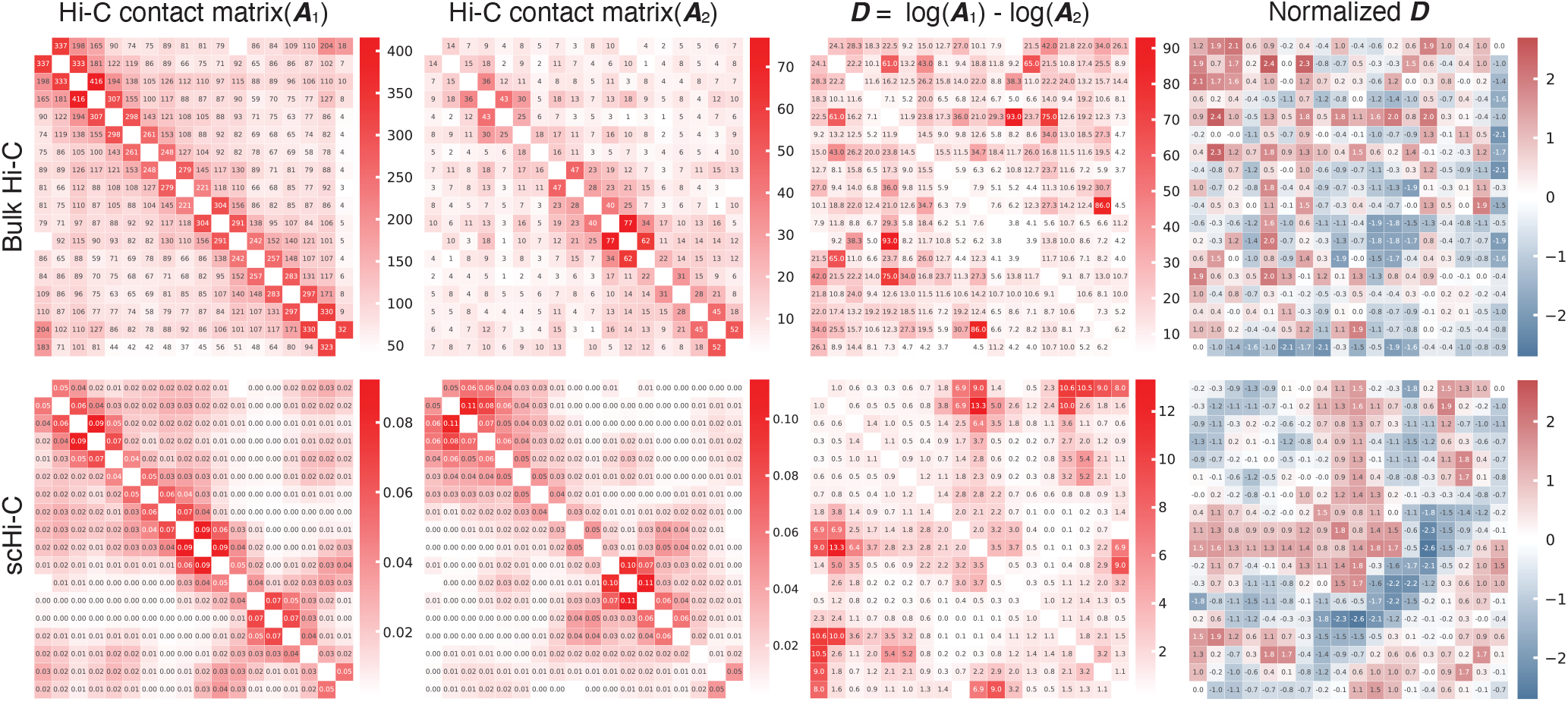
Visualization of the procedures in DiffDomain for both bulk and scHi-C data as the input for the same TAD. *Top;* bulk Hi-C contact matrices as the input (***A***_1_ and ***A***_2_); *Bottom* scHi-C contact matrices imputed by scHiCluster as the input (***A***_1_ and ***A***_2_). Cortical L2-5 pyramidal cells are condition 1. Adult astrocytes are condition 2. The TAD region is Chr3:128,350,000:129,200,000. *First* column is the input contact matrix from condition 1. *Second* column is the input contact matrix from condition 2. *Third* column is log-transformed difference matrix ***D***. *Forth* column is normalized ***D*** by iteratively standardizing its *k*-off diagonal elements, − (*N −*1) ≤*k ≤N −*1. Although the same TAD is used, the normalized ***D*** in the two rows have some differences largely due to high randomness of TAD organization in individual cells. In both bulk Hi-C and scHi-C scenarios, the TAD is identified as a reorganized TADs. This visualization is an addition to the visualization in Fig. 1A-C, highlighting that DiffDomain can work on both bulk and single-cell Hi-C data.

**Figure S2:**
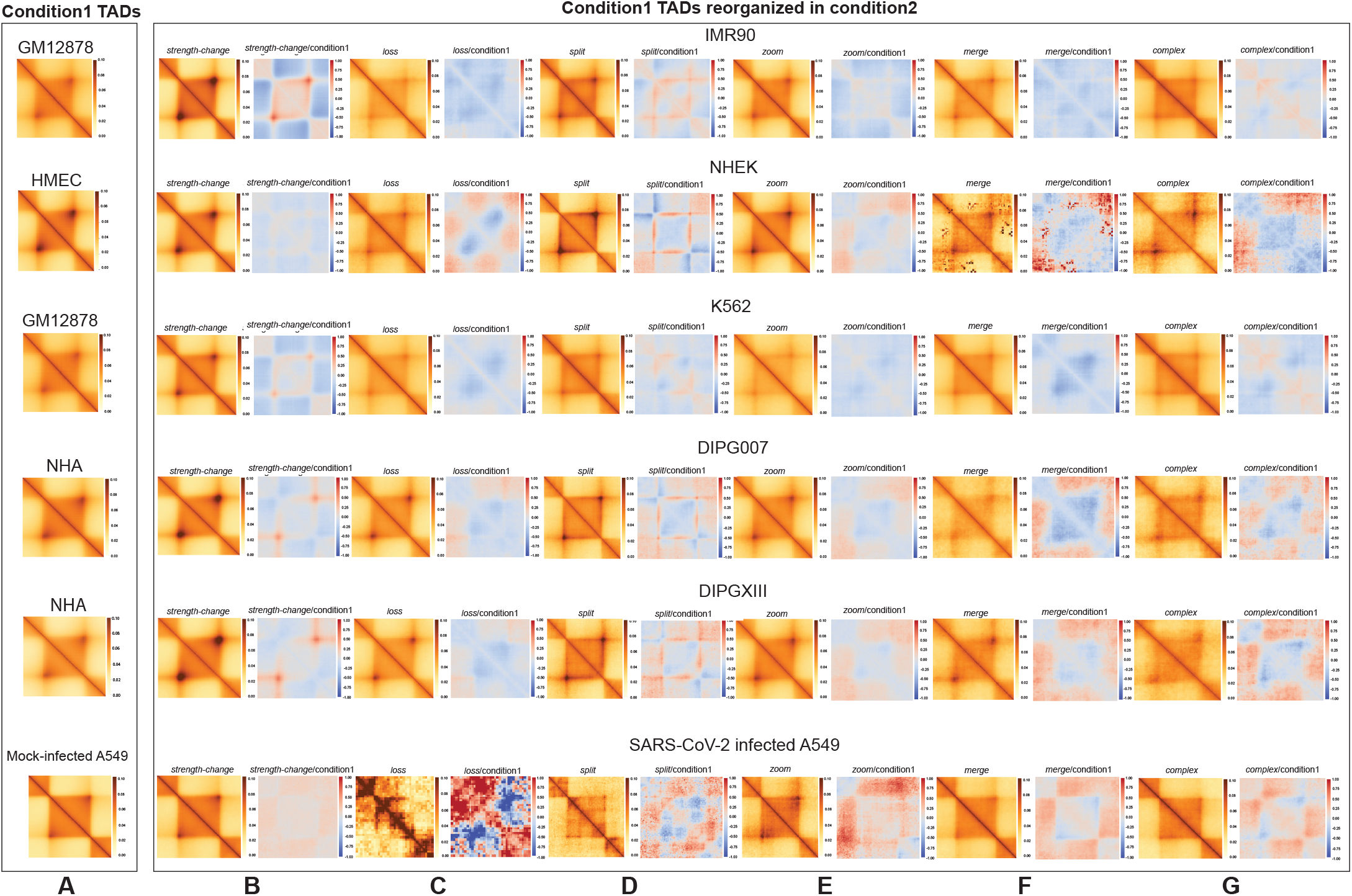
APA plot summarizing subtypes of reorganized TADs. (***A***) APA plot summarizing all TADs in condition 1 using Hi-C data from the same condition. The APA plot is consistently used as a control in the subsequent APA plots to highlight aggregated changes in the subtypes of reorganized TADs. It is referred to as condition 1 in the subsequent APA plots. (***B***) APA plot summarizing aggregated changes in *strength-change* TADs. *Left* APA plot for *strength-change* TADs using Hi-C data from condition 2; *Right* APA plot showcasing log2-transformed fold-change of APA matrices, where the numerator is from the *Left* and the denominator is from (***A***). Decreased/increased interactions within the aggregated TAD region alongside stable interactions around its boundaries support *strength-change* TAD definition. (***C***) APA plot summarizing aggregated changes in *loss* TADs. Decreased interactions within the aggregated TAD region and around its boundaries align with *loss* TAD characterization. (***D***) APA plot summarizing aggregated changes in *split* TADs. Decreased interactions within the aggregated TAD region, particularly around its center, is consistent with the definition of *split* TADs. (***E***) APA plot summarizing aggregated changes in *zoom* TADs. Increased/decreased interactions within the aggregated TAD region and between it and adjacent upstream/downstream regions are typical of *zoom* TADs. (***F***) APA plot summarizing aggregated changes in *merge* TADs. Increased interactions between the aggregated TAD region and its adjacent regions align with *merge* TAD definition. (***G***) APA plot summarizing aggregated changes in *complex* TADs. Across the six pairs of conditions (*Rows*), distinct changes within the aggregated TAD region and between it and its adjacent regions match *complex* TAD description. These pairs of conditions include two pairs of normal cell lines (GM12878 vs. IMR90, HMEC vs. NHEK) and four pairs of normal cell lines and cell lines with disease (GM12878 vs. K562, NHA vs. DIPG007, NHA vs. DIPGXIII, mock-infected vs. SARS-CoV-2 infected A549-ACE2 cells). The APA matrices are based on 25 kb resolution Hi-C data produced by FAN-C using the command ‘fanc aggregate -m -p –pixels 90 -r -e –rescale’. The APA plots are generated using python function ‘sns.heatmap’.

**Figure S3:**
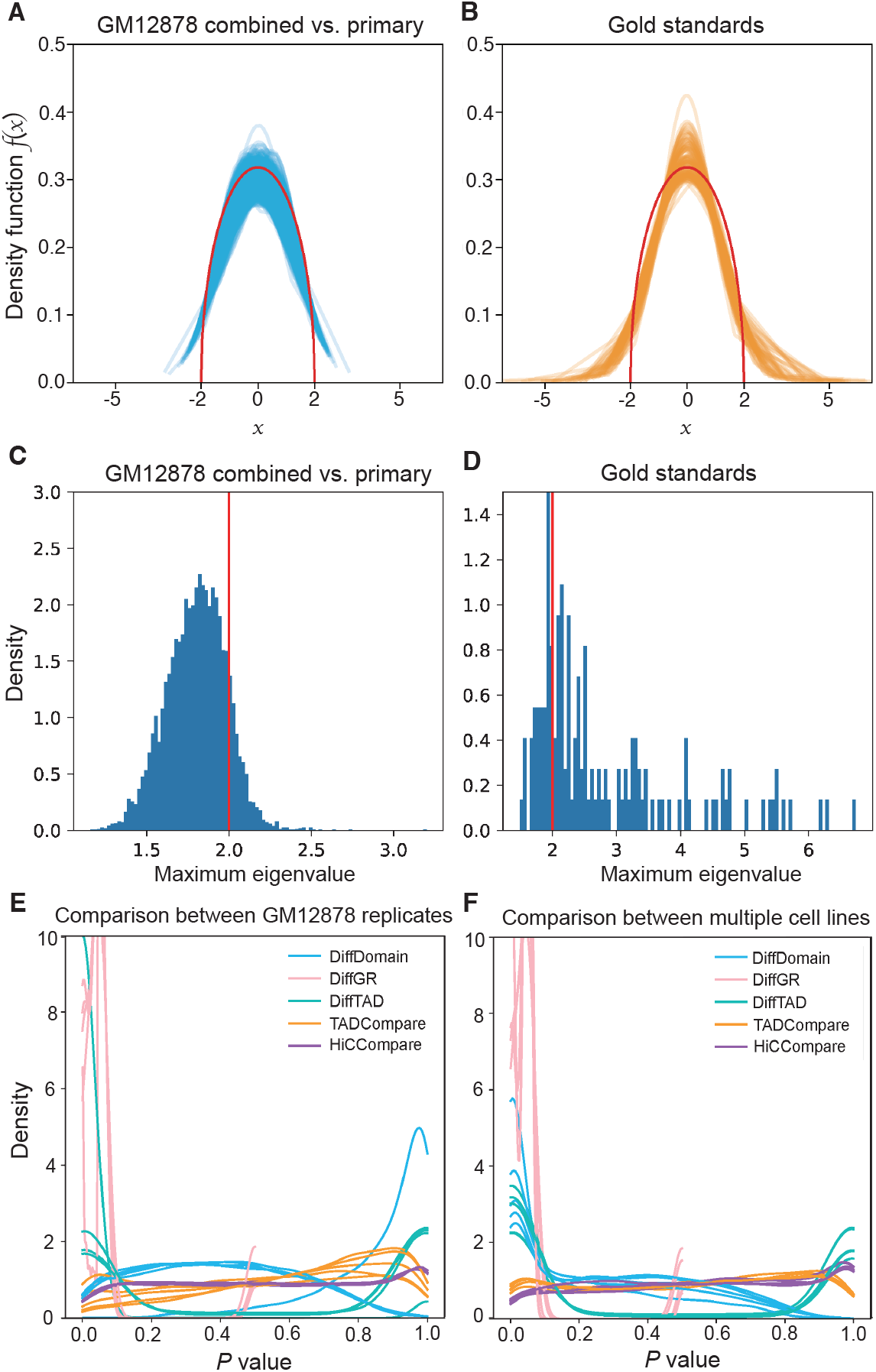
Violation of the independence assumption by 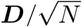 of DiffDomain has modest effects. (***A***) Estimated density functions *f*_*i*_(*x*) of empirical spectral distributions of 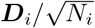, 1 *≤ i ≤* 8261,using GM12878 TADs and two GM12878 Hi-C replicates. Those 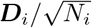, 1 *≤ i ≤* 8261, that are generated are treated as generated under *H*_0_. Red curve is the theoretical semicircular density function of generalize Wigner matrices under *H*_0_. (***B***) Estimated density functions *f*_*j*_(*x*) of empirical spectral distributions of 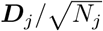, 1 *≤ j ≤* 146, that are generated using manually collected 65 reorganized TADs. Those 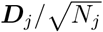, 1 ≤*j ≤*146, are treated as generated under *H*_1_. (***C***) Histogram shows the largest eigenvalues of 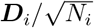, 1 *≤i ≤*8261, same as in (***A***). The vertical red line is the theoretical limit of the largest eigenvalue of a generalized Wigner matrix (*H*_0_). (***D***) Histogram shows the largest eigenvalues of 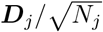, 1 ≤*j ≤*146, same as in (***B***). (***E***) Estimated density curves of unadjusted *P* values that are computed in investigating TAD reorganization between multiple pairs of the GM12878 Hi-C replicates (Supplementary Table S1). Each curve corresponds to the comparison between a pair of GM12878 Hi-C replicates. The unadjusted *P* values are treated as the *P* values under *H*_0_. *P* values follow a uniform distribution when *H*_0_ is true and model assumptions are satisfied. (***F***) Estimated density curves of unadjusted *P* values that are computed in comparing multiple pairs of human cell lines (Supplementary Table S2). The unadjusted *P* values are mixtures of *P* values under *H*_1_ and *P* values under *H*_0_, thus they are not uniformly distributed, as expected. DiffDomain assumes that 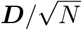 is a generalized Wigner matrix, a symmetric random matrix with independent upper diagonal entries. The independence assumption on the upper diagonal entries is violated by 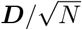 given that Hi-C contact frequencies positively correlate with each other among nearby chromosome bins. The analyses find broad agreements between empirical properties of 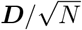 and theoretical properties of generalized Wigner matrices under *H*_0_ and substantial disagreements under *H*_1_, showing violation of independence assumption of DiffDomain has modest effects.

**Figure S4:**
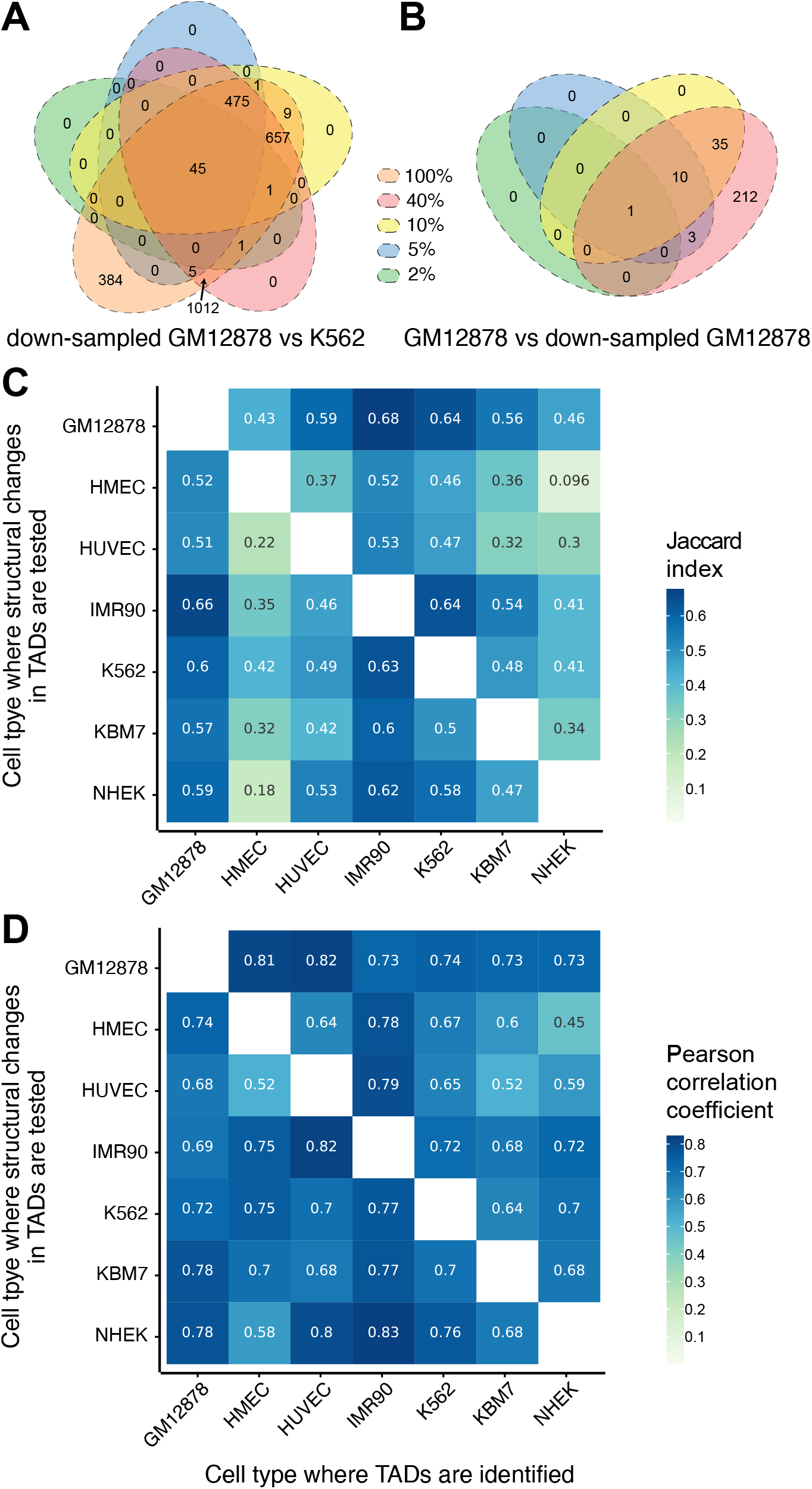
DiffDomain is robust to sequencing depths and Hi-C resolution. (***A***) Venn diagram showing reproducibility of reorganized TADs that are identified by DiffDomain using a varied sequencing depth. (***B***) Venn diagram showing reproducibility between original and down-sampled Hi-C data from GM12878. Sequences were down-sampled at 2, 5, 10, 40% sequencing reads. (***C***) Heatmap showing Jaccard index between reorganized TADs that are identified by DiffDomain using 10 kb and 25 kb resolution Hi-C data. (***D***) Heatmap showing Pearson correlation coefficient between log-transformed BH-adjusted *P* values. One list of *P* values is computed by DiffDomain using 10 kb resolution Hi-C data. Another list of *P* value is computed using 25 kb resolution Hi-C data.

**Figure S5:**
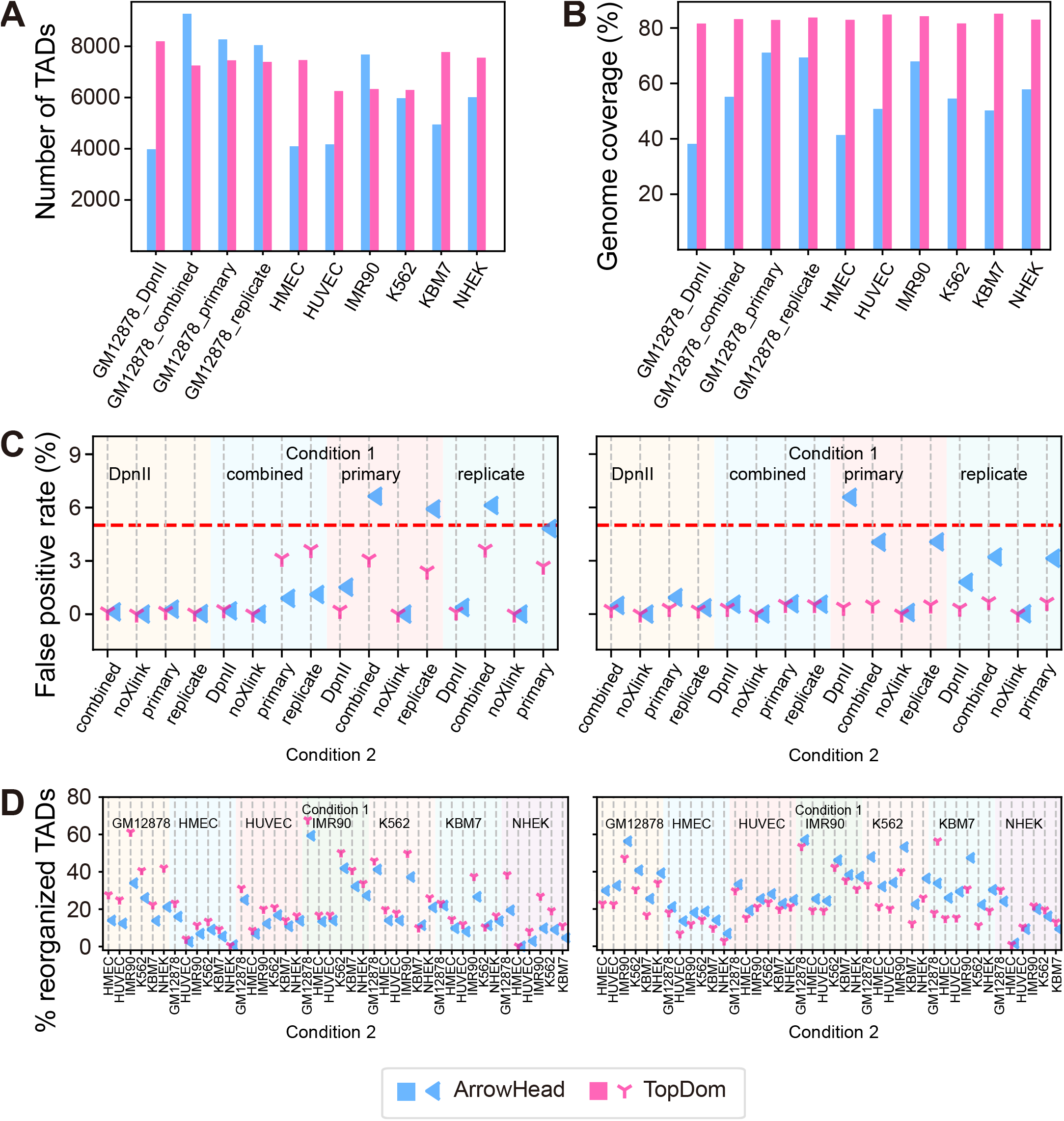
DiffDomain is robust to TAD callers. (***A***) Bar graph showing the number of TADs called by Arrowhead and TopDom across GM12878 replicates and different cell lines. (***B***) Bar graph showing the genome coverage of TADs. (***C***) Dot plot showing the false positive rates of DiffDomain. False positive rate is estimated as the proportion of reorganized TADs between GM12878 replicates. Hi-C resolution is 10 kb and 25 kb, respectively. (***D***) Dot plot showing the proportion of reorganized TADs between different human cell lines. Hi-C resolution is 10 kb and 25 kb, respectively. Overall, although the TADs called by Arrowhead and TopDom have clear differences in the number of TADs and genome coverage, DiffDomain has comparable FPRs and proportions of reorganized TADs between cell lines, suggesting that DiffDomain is robust to TAD callers.

**Figure S6:**
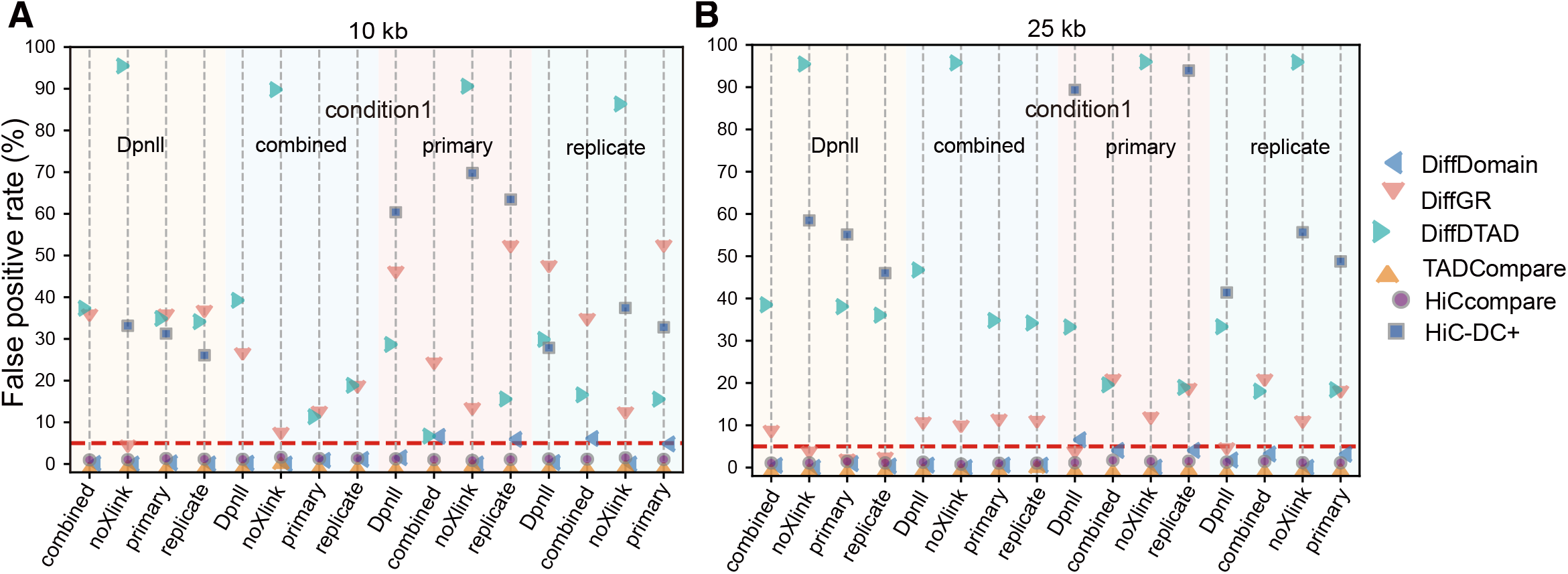
Method comparison using FPR. The resolution of Hi-C data are 10 kb (***A***) and 25 kb (***B***). Given a pair of GM12878 Hi-C replicates, FPR is estimated as the percentage of identified reorganized GM12878 TADs. The GM12878 Hi-C replicate “noXlink” as condition 1 is missing because it does not have enough read coverage to call TADs by Arrowhead algorithm at 10 kb resolution. The horizontal dashed red lines represent significant level *α* = 0.05, which is also the expected FPR under *H*_0_. Note that HiC-DC+ requires at least two Hi-C replicates in each condition. The Hi-C replicate *combined* is created by merging the replicates *primary* and *replicate*. Thus, HiC-DC+ is not applied to compare replicate *combined* with other replicates. These results show that DiffDomain, TADCompare, and HiCcompare have better control of FPR than the other three methods.

**Figure S7:**
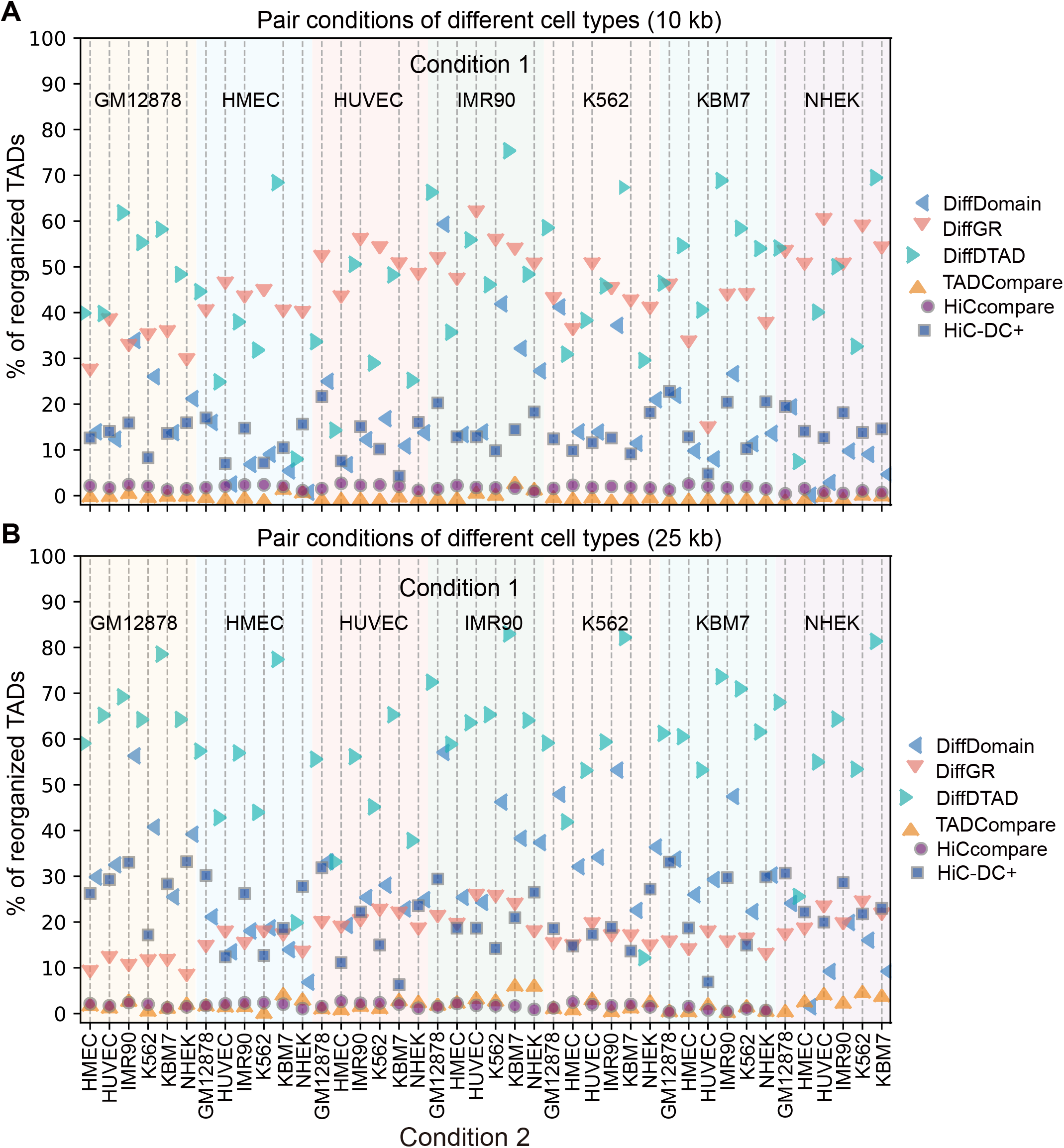
Method comparison using the percentages of identified reorganized TADs between pairs of human cell lines. The resolution of Hi-C data are 10 kb (***A***) and 25 kb (***B***). The proportions of reorganized TADs identified by HiCcompare and TADCompare are below 5%, suggesting that HiCcompare and TADCompare are too conservative. The proportions of reorganized TADs identified by HiC-DC+ are small (below 30%) while the FPRs of HiC-DC+ are high (ranging from 25% to 70% (Supplementary Fig. S6)), suggesting that HiC-DC+ is not optimal for identifying reorganized TADs. These results show that both HiC-DC+ and HiCcompare, specifically designed for identifying differential chromatin interactions, are not optimal for identifying reorganized TADs.

**Figure S8:**
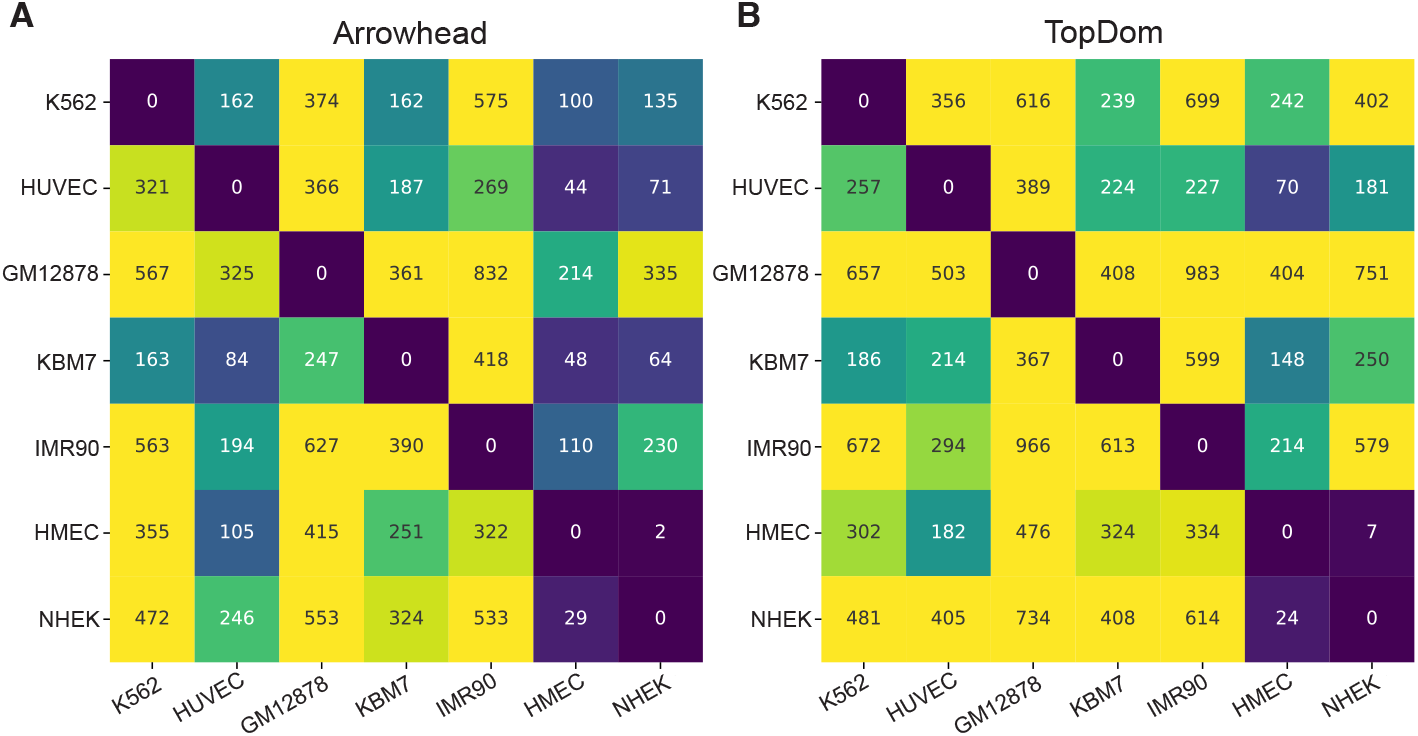
Comparing DiffDomain with TADsplimer [25] in terms of the numbers of *split* and *merge* TADs. (***A***) TADs are called by Arrowhead, then the subset of reorganized TADs is identified by DiffDomain. (***B***) TADs are called by TopDom, then the subset of reorganized TADs is identified by DiffDomain. For each pair of cell types, the number of *split* and *merge* TADs is calculated as the summation of the number of *split* TADs and the number of *merge* TADs, the same as the calculation in Fig. 2E in Wang et al. [25]. Overall, the results in (***B***) are much more similar to the original results in Fig. 2E [25] than the results in (***A***). This is expected because TopDom has similar performance with the built-in TAD caller in TADsplimer.

**Figure S9:**
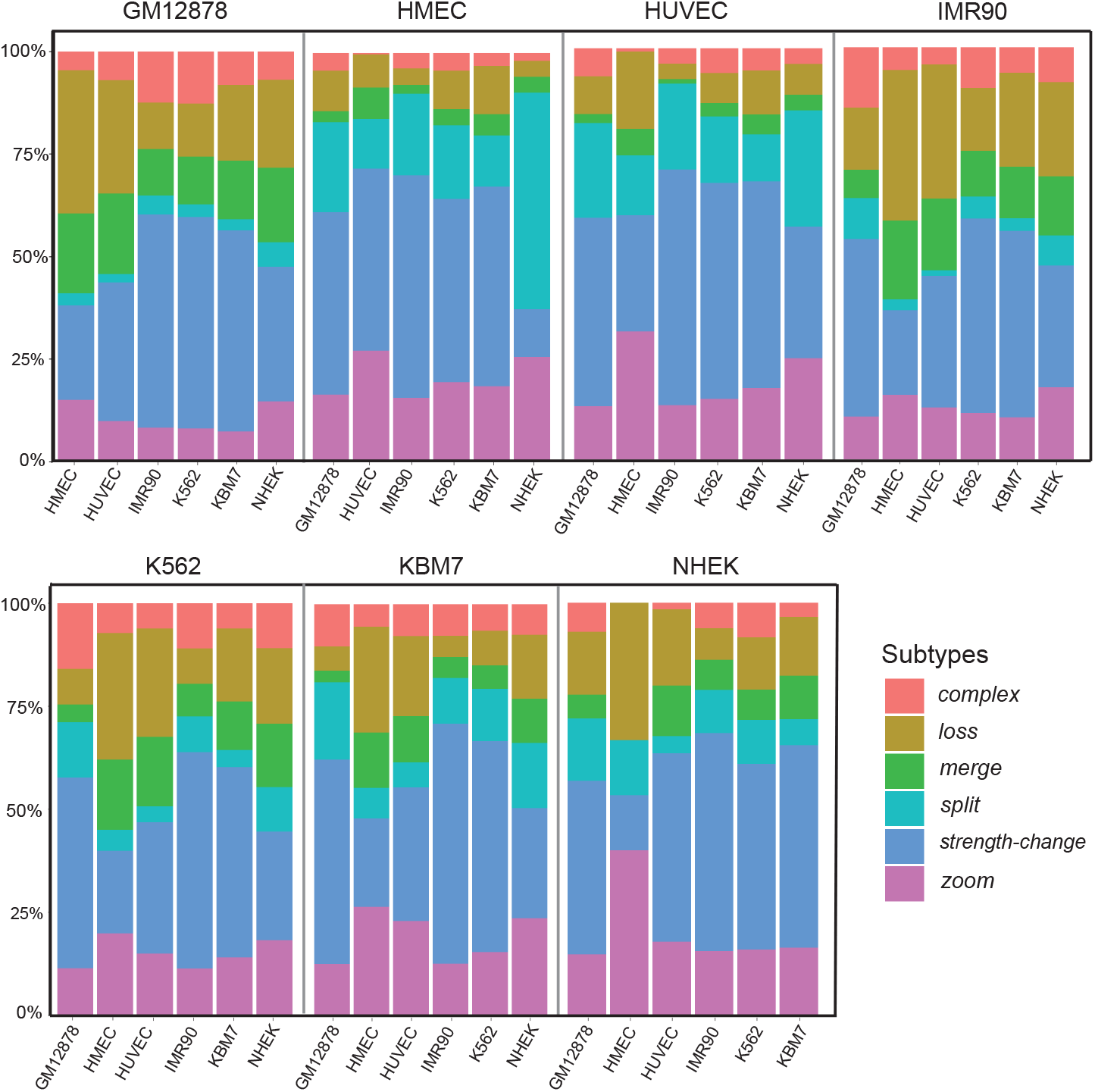
Percentages of subtypes of reorganized TADs between human cell lines that are identified by DiffDomain. Cell lines on the top of stacked barplots are condition 1, cell lines on the *X*-axis are condition 2.

**Figure S10:**
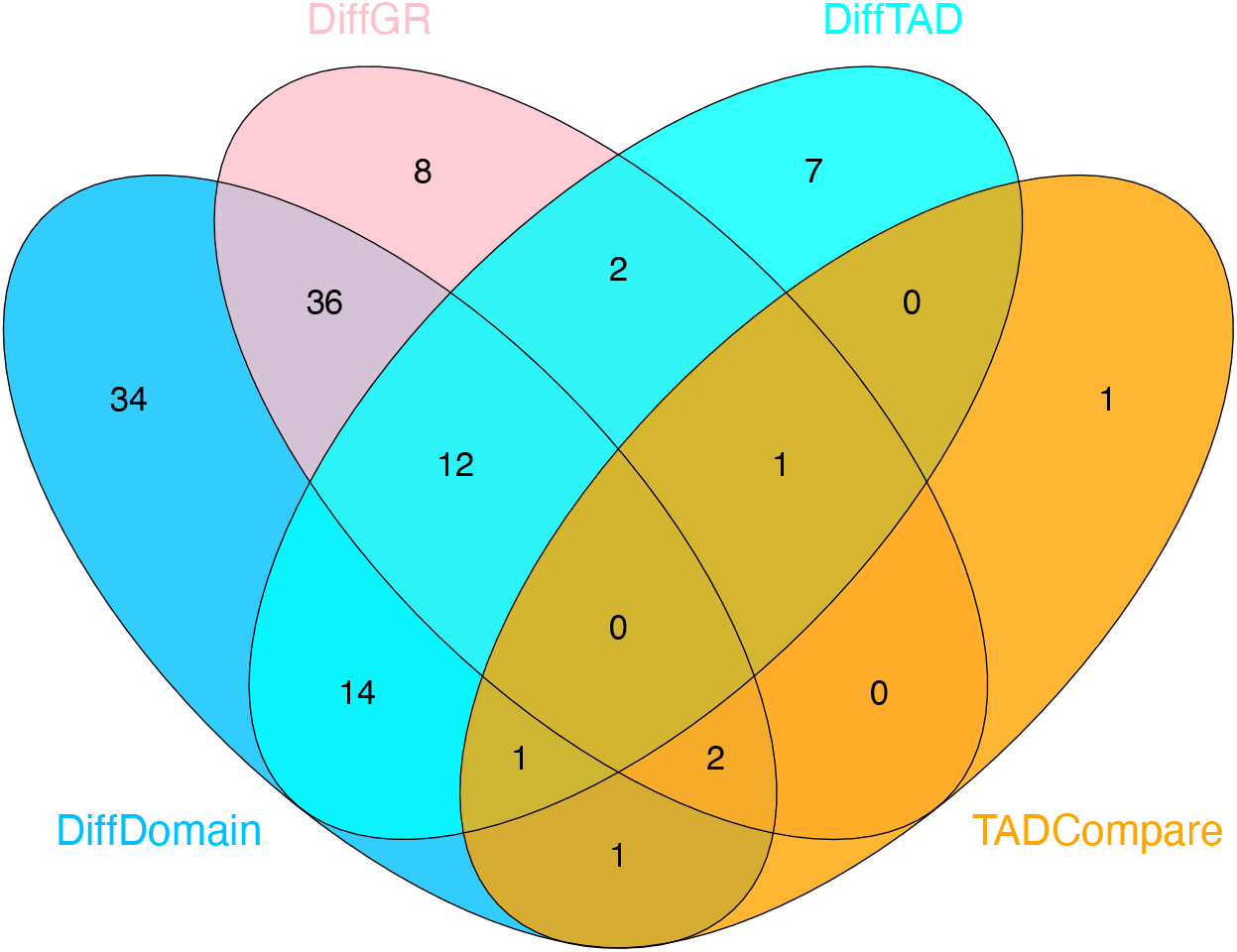
Venn diagram showing the agreements between the lists of correctly identified truly reorganized TADs. There are 34 truly reorganized TADs are only correctly identified as such by DiffDomain. In contrast, only 8, 7, and 1 truly reorganized TADs are specifically correctly identified as such by DiffGR, DiffTAD, and TADCompare, respectively. Compared with DiffDomain, DiffGR, DiffTAD, and TADCompare only uniquely identify 11, 10, and 1 truly reorganized TADs, respectively, demonstrating the advantage of DiffDomain.

**Figure S11:**
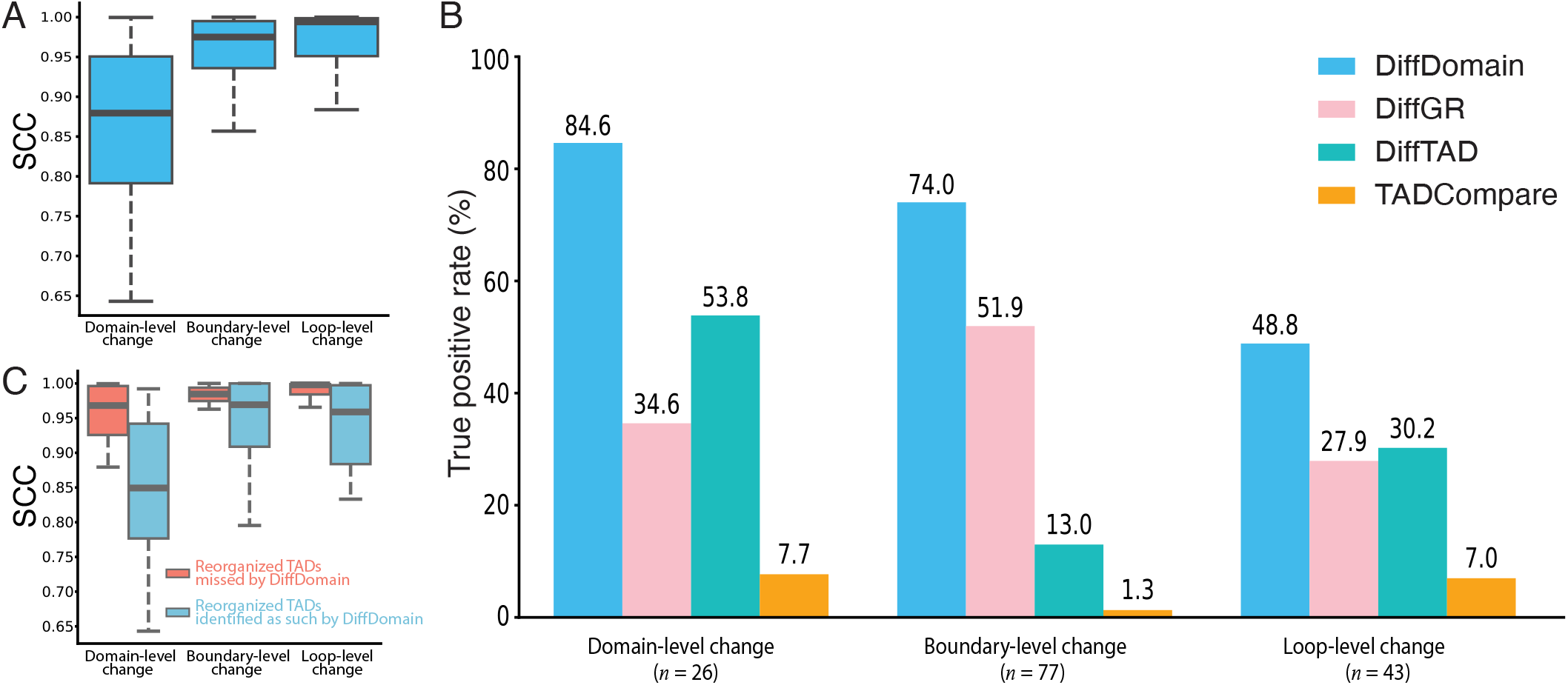
Method comparison based on groups of truly reorganized TADs. (***A***) Boxplot showing the distribution of SCC score of truly reorganized TADs between 146 pairs of comparisons. Each data point represents the SCC score between a pair of Hi-C contact maps for a truly reorganized TAD. (***B***) Barplot showing TPRs of DiffDomain and alternative methods across the three groups of truly reorganized TADs. (***C***) Boxplot comparing the SCC scores between the correctly identified truly reorganized TADs and those missed by DiffDomain. The three groups of truly reorganized TADs-domain-level change, boundary-level change, and loop-level change-are broadly defined based on the original descriptions of changes in these TADs in the literature.

**Figure S12:**
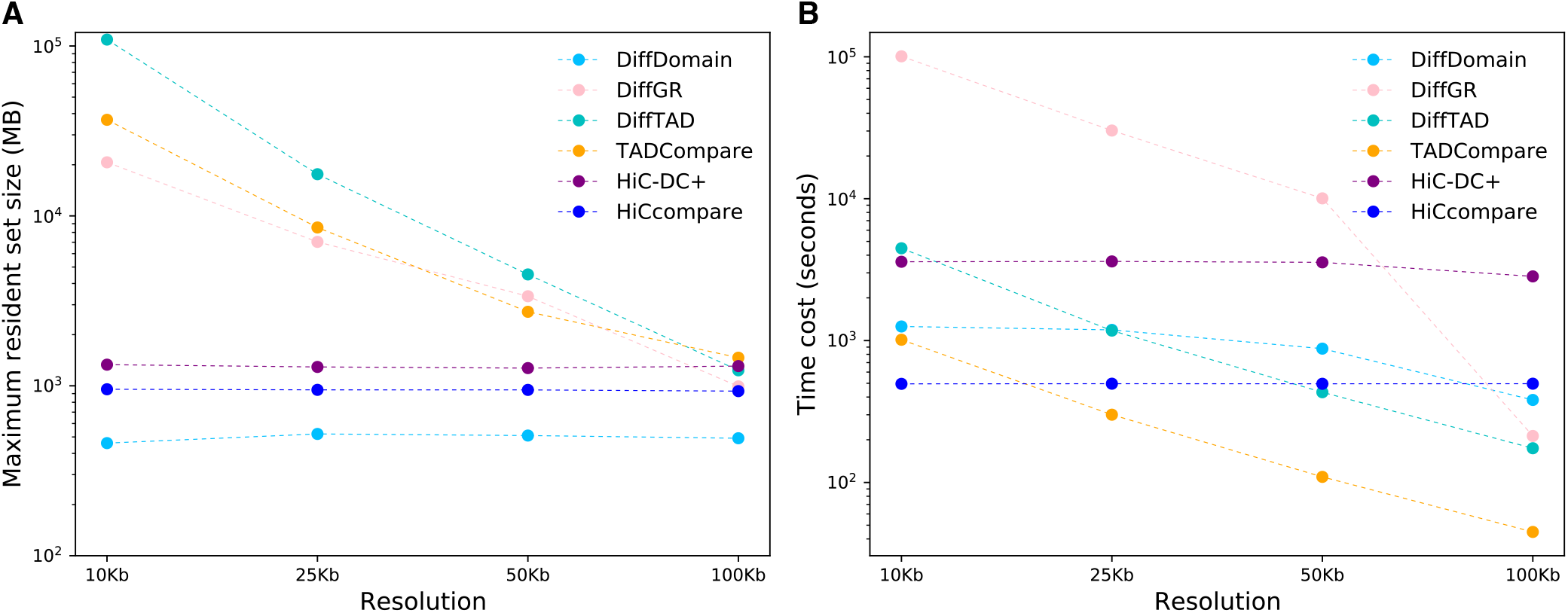
Method comparison by memory usage and computation time. (***A***) Memory usage comparison in terms of maximum resident set size (MB). (***B***) Computation time comparison in terms of time cost (seconds). Statistics are estimated by Linux kernel function */usr/bin/time -f “%M %e”*. Each dot represents the average from 10 repeated experiments. Note that DiffDomain, HiC-DC+, and HiCcompare have stable memory usage and computation time because their bottleneck is extracting contact matrices from .hic files by *straw* package.

**Figure S13:**
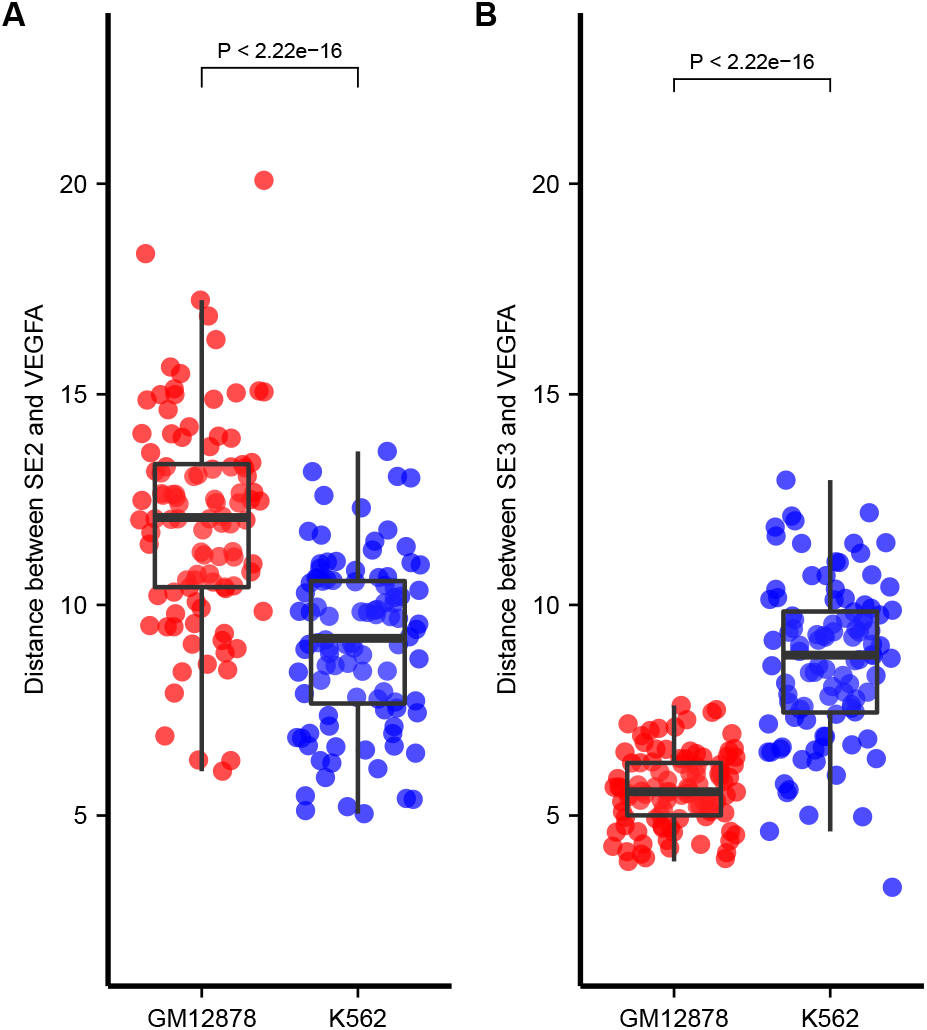
Spatial proximities between VEGFA and super-enhancers in GM12878 and K562. (***A***) Distances between VEGFA and SE2 (super-enhancer Chr6:43871593-43959002). (***B***) Distances between VEGFA and SE3 (super-enhancer Chr6:44008142-44046149). Each dot represents a Euclidean distance between VEGFA and a super-enhancer in a given 3D structure. Totally, 100 3D structures for each cell type are generated by Chrom3D [47]. In the box plots, the middle line represents the median; the lower and upper lines correspond to the first and third quartiles; and the upper and lower whiskers extend to values no farther than 1.5*×*IQR.

**Figure S14:**
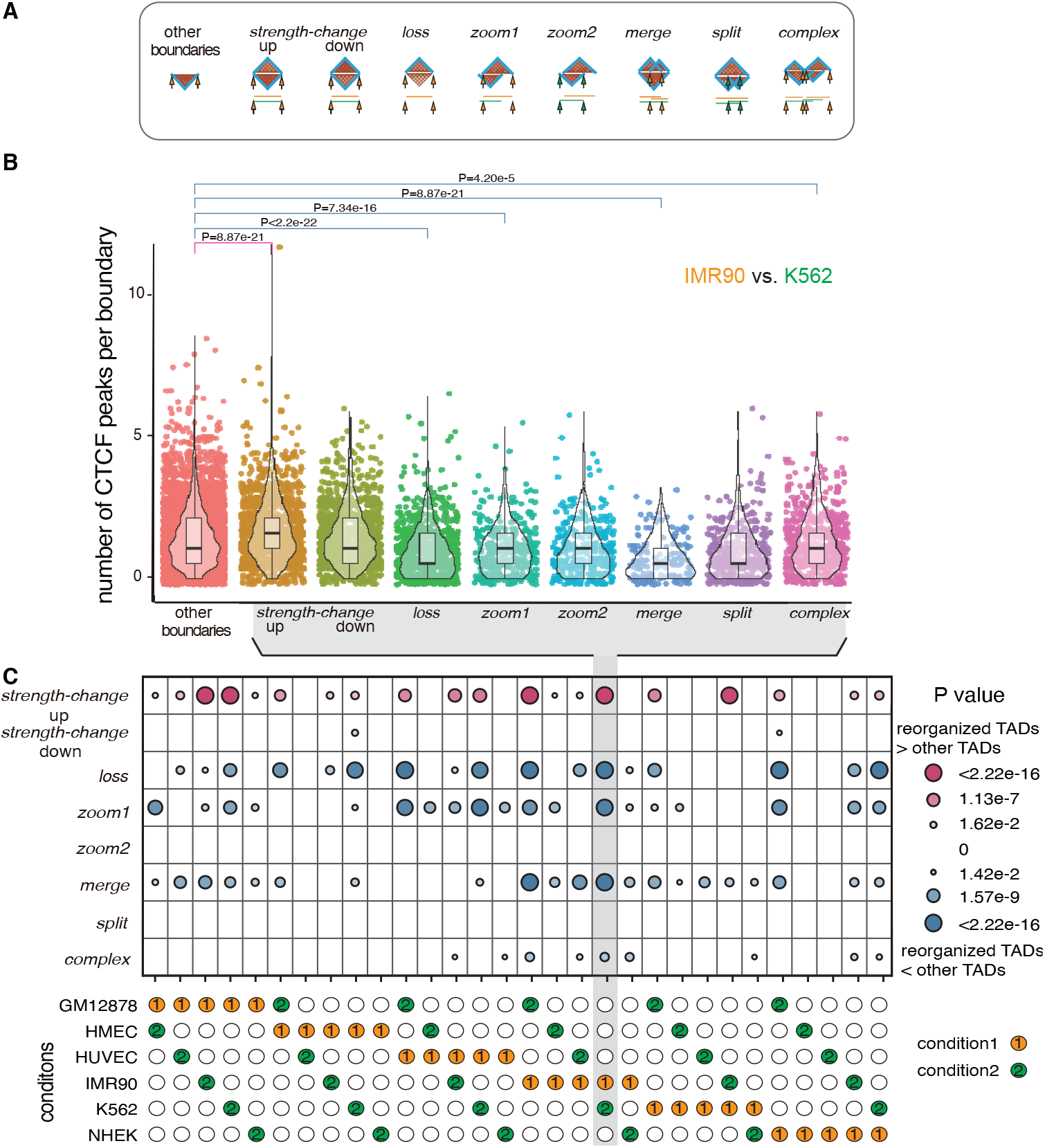
Variations in the number of CTCF peaks related to reorganized TADs. (***A***) Cartoon diagram illustrating the comparison of CTCF peaks between different TAD boundary groups. The heatmap diagram illustrates TADs in condition 1 (top) and condition 2 (bottom), with lines representing TAD regions of the same condition sharing the same color. The first group are TAD boundaries from condition 2 TADs not overlapping with the subset of condition 1 TADs reorganized in condition 2, serving as a control. This group of TAD boundaries is called as other boundaries. Subsequent groups are defined by TAD reorganization subtypes. These include TAD boundaries from *strength-change up* TADs, *strength-change down* TADs, *loss* TADs, and zoom TADs (split into two groups: *zoom 1*, lost in condition 2; *zoom 2*, gained in condition 2). Addition groups include TAD boundaries lost due to TAD merging (*merge*) and TAD boundaries gained due to TAD splitting (*split*). Lastly, TAD boundaries associated with *complex* TADs but existing only in condition 1 are classified as *complex*. These groups are illustrated using vertical arrows. The number of CTCF peaks in each group are all counted using data from condition 2. (***B***) Boxplots showing the number of CTCF peaks across the TAD boundary groups. Here, condition 1 is IMR90, condition 2 is K562. Each dot represents one TAD boundary. The number of CTCF peaks in the group of other boundaries is compared with those from the remaining boundary groups. *P* values are computed using one-sided MannWhitney U test, a nonparametric test dealing with asymmetric distributions of the number of CTCF sites. For the sake of clarity, insignificant *P* values (*P >* 0.05) are excluded from the boxplot. (***C***) Heatmap showing the comparison of the number of CTCF peaks between the group of other boundaries and the remaining boundary groups across pairs of cell types. Each column corresponds to a specific cell type pair, with cell types indicated below the heatmap. Each row represents one TAD group. The heatmap cells represent significant levels (log_10_(*P*)) computed from comparing the corresponding TAD boundary group (row) with the group of other boundaries within a given cell type pair (column). Insignificant *P* values (*P >* 0.05) are left out from the heatmap to enhance clarity. Similar significant results are observed across the various cell type pairs. In the box plots, the middle line represents the median; the lower and upper lines correspond to the first and third quartiles; and the upper and lower whiskers extend to values no farther than 1.5*×*IQR.

**Figure S15:**
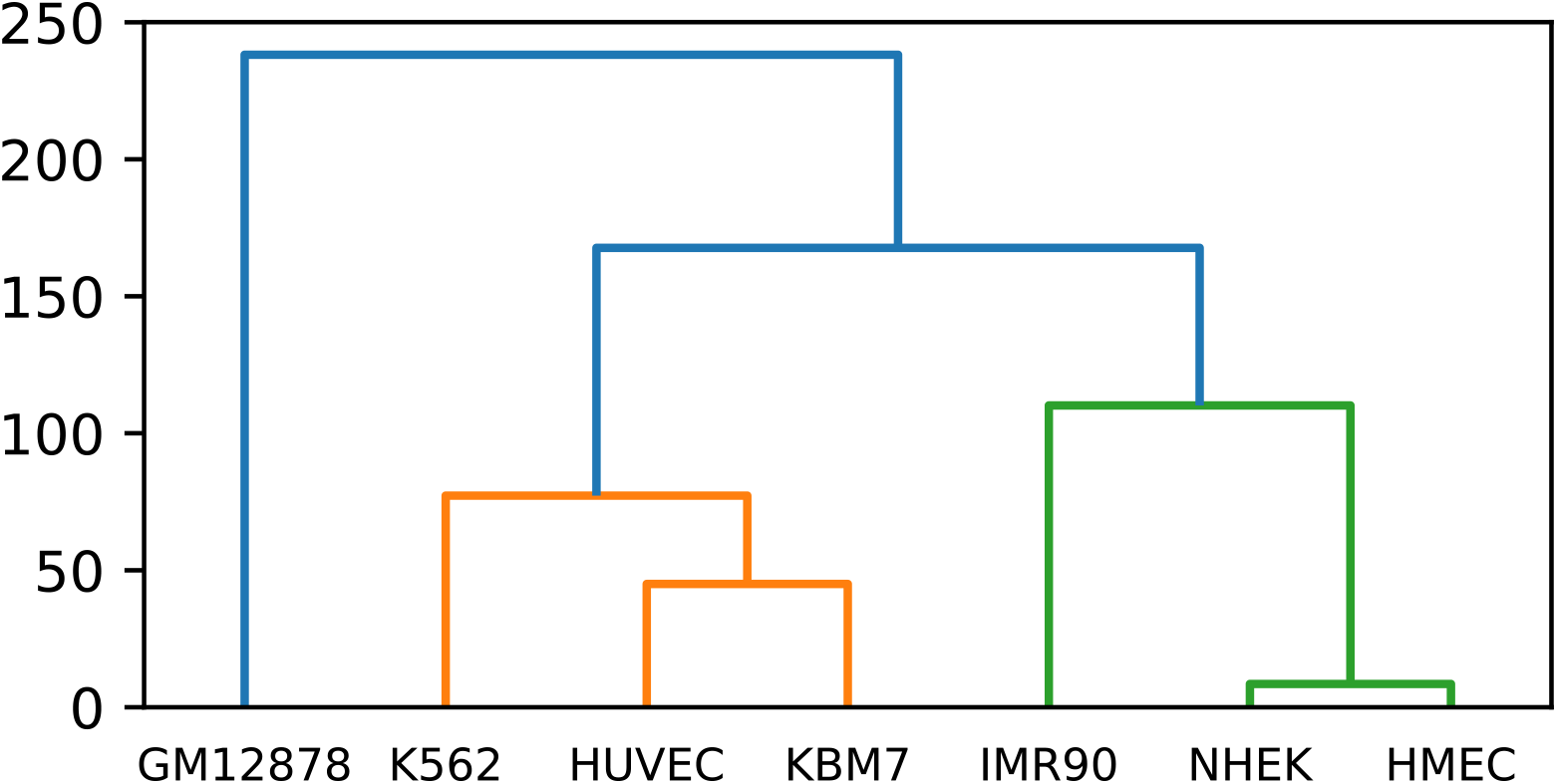
Hierarchical clustering of cell types using the number of *split* TADs.

**Figure S16:**
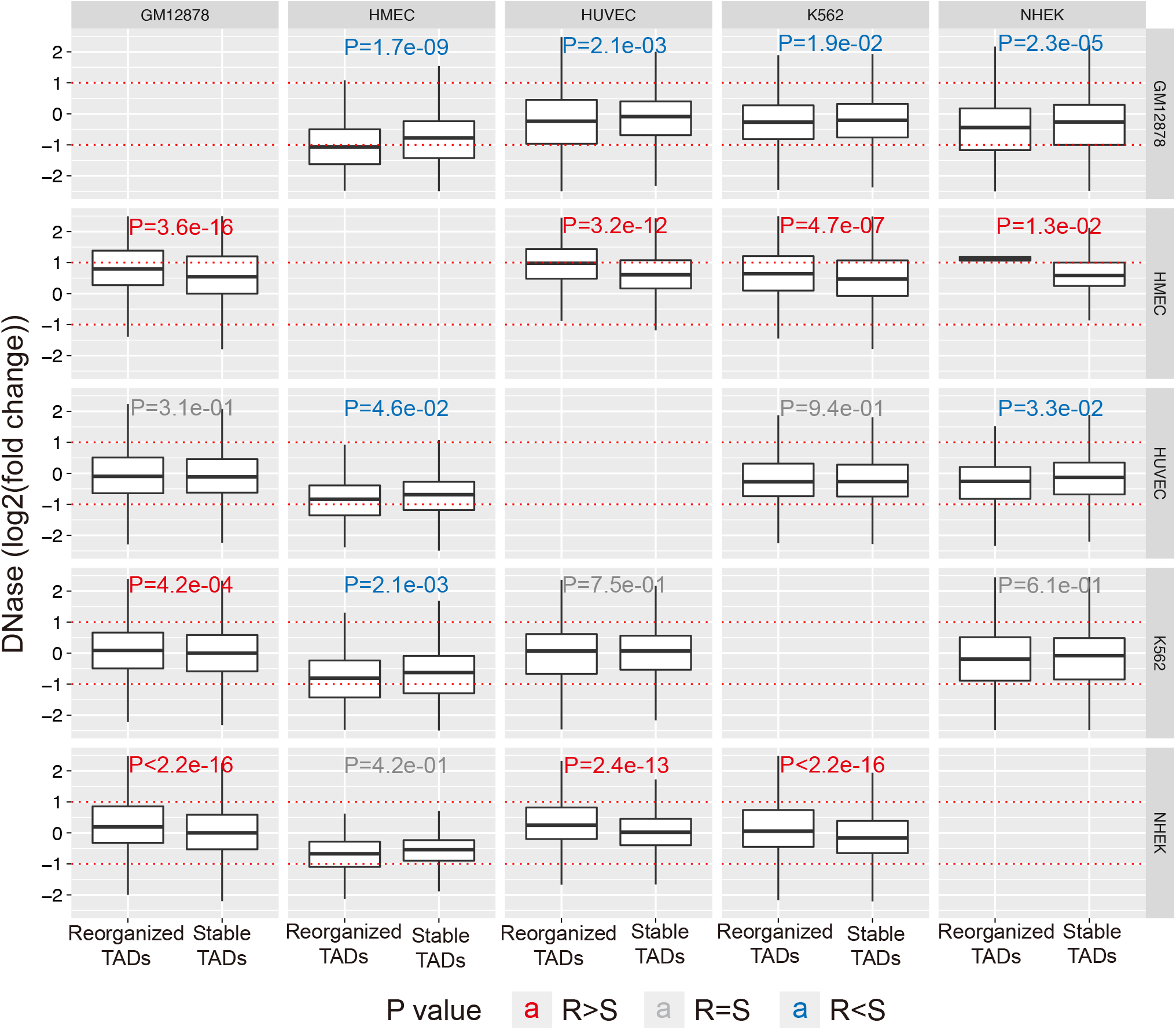
Reorganized TADs tend to gain chromatin accessibility. We describe the analysis for the pair of GM12878 and HMEC cell types (first column and second row). we grouped GM12878 TADs into two groups: 1) TADs that are reorganized in HMEC cell type, called reorganized TADs; 2) the other TADs, called stable TADs. For each GM12878 TAD, we compute the fold-change of DNase peak coverage of the TAD in HMEC cell type to the coverage in GM12878 cell type. Then the fold-change is *log*_2_ transformed, where a positive/negative value means that the TAD gains/losses chromatin accessibility in HMEC. The boxplots show that the reorganized TADs have significantly (*P* = 3.6 ×10^*−*16^) higher coverage of chromatin accessibility than the stable TADs. We repeat the analysis to the other pairs of cell types. Here each column represents the cell type where TADs are called, i.e., condition 1. Each row represents the cell type where TADs are tested whether they are reorganized, i.e., condition 2. The *P* values (*P*) in red and blue colors mean that reorganized TADs have significantly (*P ≤*0.05) higher and lower mean change in chromatin accessibility. While the p values in grey mean that the two groups of TADs have no significant difference in change of chromatin accessibility. *P* values are computed by Mann-Whitney test. IMR90 and KBM7 are excluded because they do not have the ENCODE DNase data at the UCSC Genome Browser. Although reorganized TADs have significantly lower fold-change in chromatin accessibility in some pairs of comparisons, the levels of significance are much smaller than those from the comparisons where reorganized TADs have significantly higher fold-change in chromatin accessibility. In the box plots, the middle line represents the median; the lower and upper lines correspond to the first and third quartiles; and the upper and lower whiskers extend to values no farther than 1.5*×*IQR.

**Figure S17:**
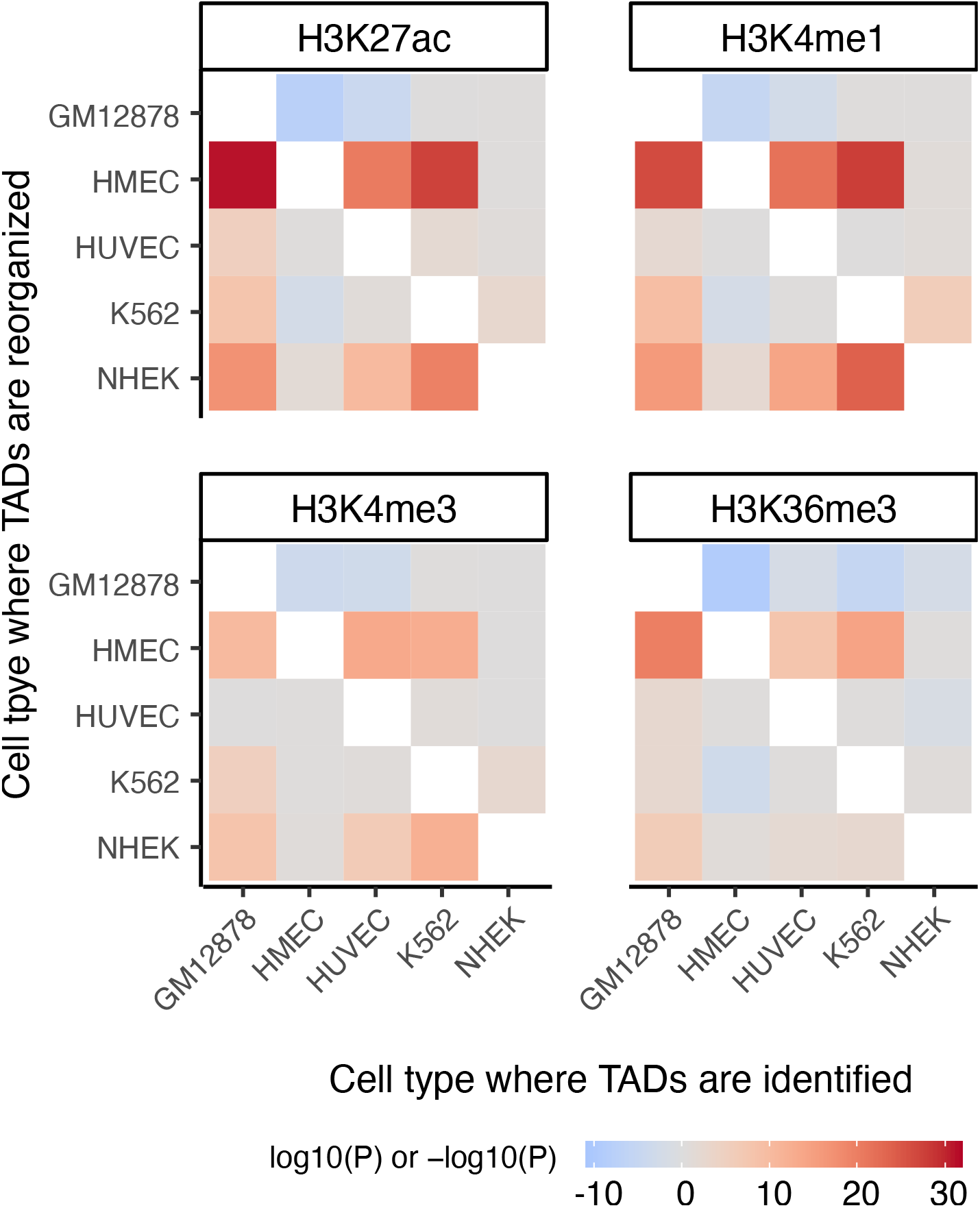
Reorganized TADs tend to gain histone modifications that are signals for enhancers and active transcription. P values are calculated as the same in the case of DNase signals shown in Supplementary Fig. S16. *P* values are transformed by −log_10_(*P*) if reorganized TADs have higher fold-change in signals than the other TADs. Otherwise, *P* values are transformed by log_10_(*P*). Missing off-diagonal elements mean that the corresponding marks do not exist in the corresponding cell types.

**Figure S18:**
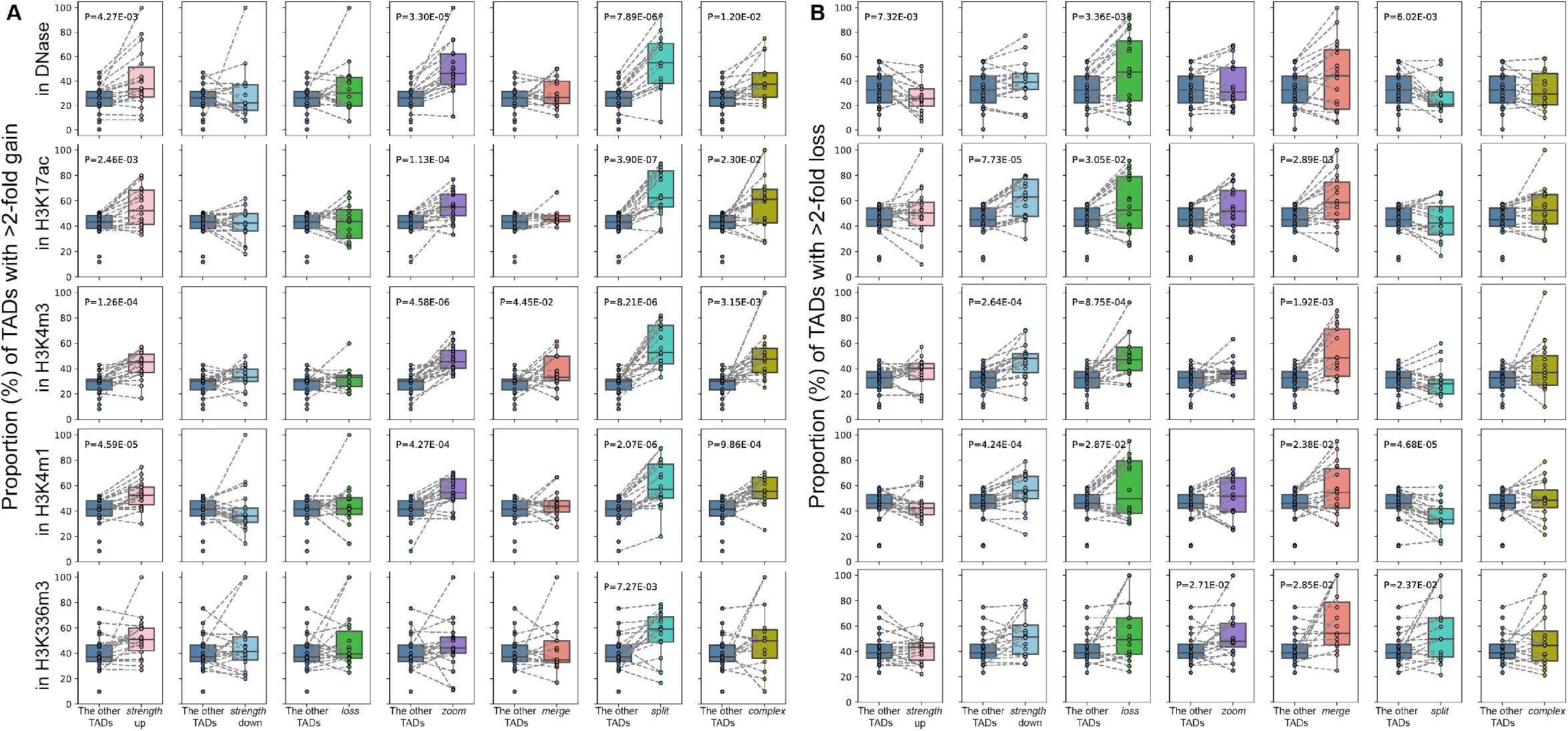
Distinct associations between TAD reorganization subtypes and chromatin accessibility as well as histone modifications. (***A***) Boxplot comparing reorganized TADs with subtypes of reorganized TADs in terms of increasing in chromatin accessibility and histone modification signals. We explain the boxplot for DNase peak coverage (*top left*). The proportion (Y-axis) is computed as follows: within a given cell type pair, for the subset of *strength-change up* TADs, the proportion is computed as *M/N*. Here, *M* is the number of *strength-change up* TADs with at least 2-fold increase in DNase peak coverage, and *N* is the number of *strength-change up* TADs with increase (fold-change *>* 1) in DNase peak coverage. Similarly, the proportion is computed for the other TAD subsets. The proportion represents the fraction of TADs with at least 2-fold increasing in DNase peak coverage among the TADs with increased DNase peak coverage, The connection between two proportions from the same cell type pair comparison is indicated by a dash line. In total, the dataset contains 20 pairs of data points, derived from 20 pairwise comparisons among GM12878, HMEC, HUVEC, K562, and NHEK cell types. Insignificant *P* values are omitted from the boxplots for clarity. (***B***) Boxplot comparing reorganized TADs with subtypes of reorganized TADs in terms of decreasing in chromatin accessibility and histone modification signals.

**Figure S19:**
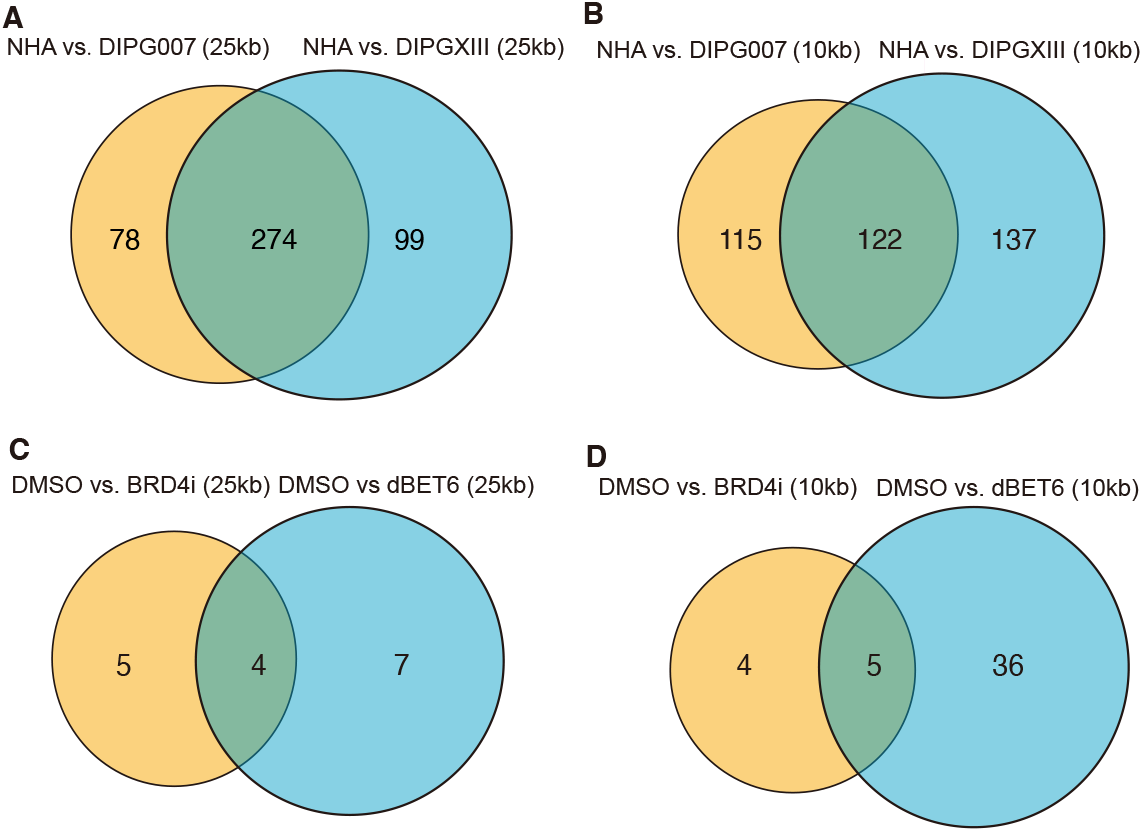
Agreements between reorganized TADs in DIPG cell lines. (***A***) Venn diagram showing the agreement between (1) NHA TADs that are reorganized in DIPG007 and (2) NHA TADs that are reorganized in DIPGXIII. Hi-C resolution is 25 kb. (***B***) Same as (***A***), except for Hi-C resolution at 10 kb. (***C***) Venn diagram showing the agreement between (1) DMSO TADs that are reorganized in BRD4i and (2) DMSO TADs that are reorganized in dBET6. Hi-C resolution is 25 kb. (***D***) Same as (***C***), except for Hi-C resolution at 10 kb. Abbreviation: NHA, normal human astrocytes; DIPG007 and DIPGXIII, pediatric high-grade glioma patient-derived cell lines; DMSO, DIPG007 with dimethyl sulfoxide treatment (used as control treatment); BRD4i, DIPG007 with the BET BRD inhibitor treatment; dBET6, DIPG007 with BRD degrader treatment.

**Figure S20:**
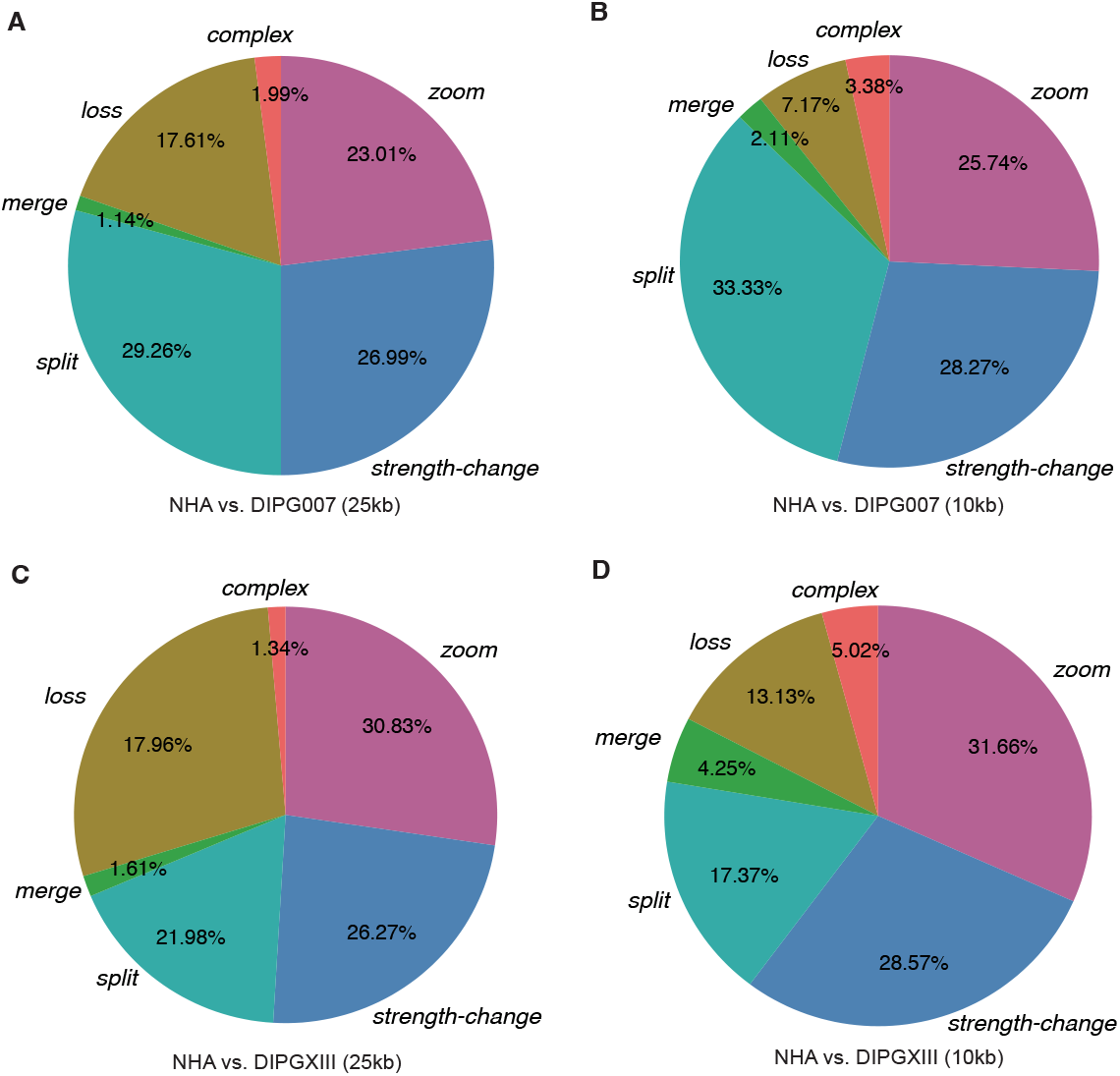
Percentages of subtypes of NHA TADs that are reorganized in DIPG cell lines. (***A***) Pie chart showing the percentages of subtypes of NHA TADs that are reorganized in DIPG007. Hi-C resolution is 25 kb. (***B***) Same as (***A***), except for Hi-C resolution at 10 kb. (***C***) Pie chart showing the percentages of subtypes of NHA TADs that are reorganized in DIPGXIII. Hi-C resolution is 25 kb. (***D***) Same as (***C***), except for Hi-C resolution at 10 kb. Abbreviation: NHA, normal human astrocytes; DIPG007 and DIPGXIII, pediatric high-grade glioma patient-derived cell lines.

**Figure S21:**
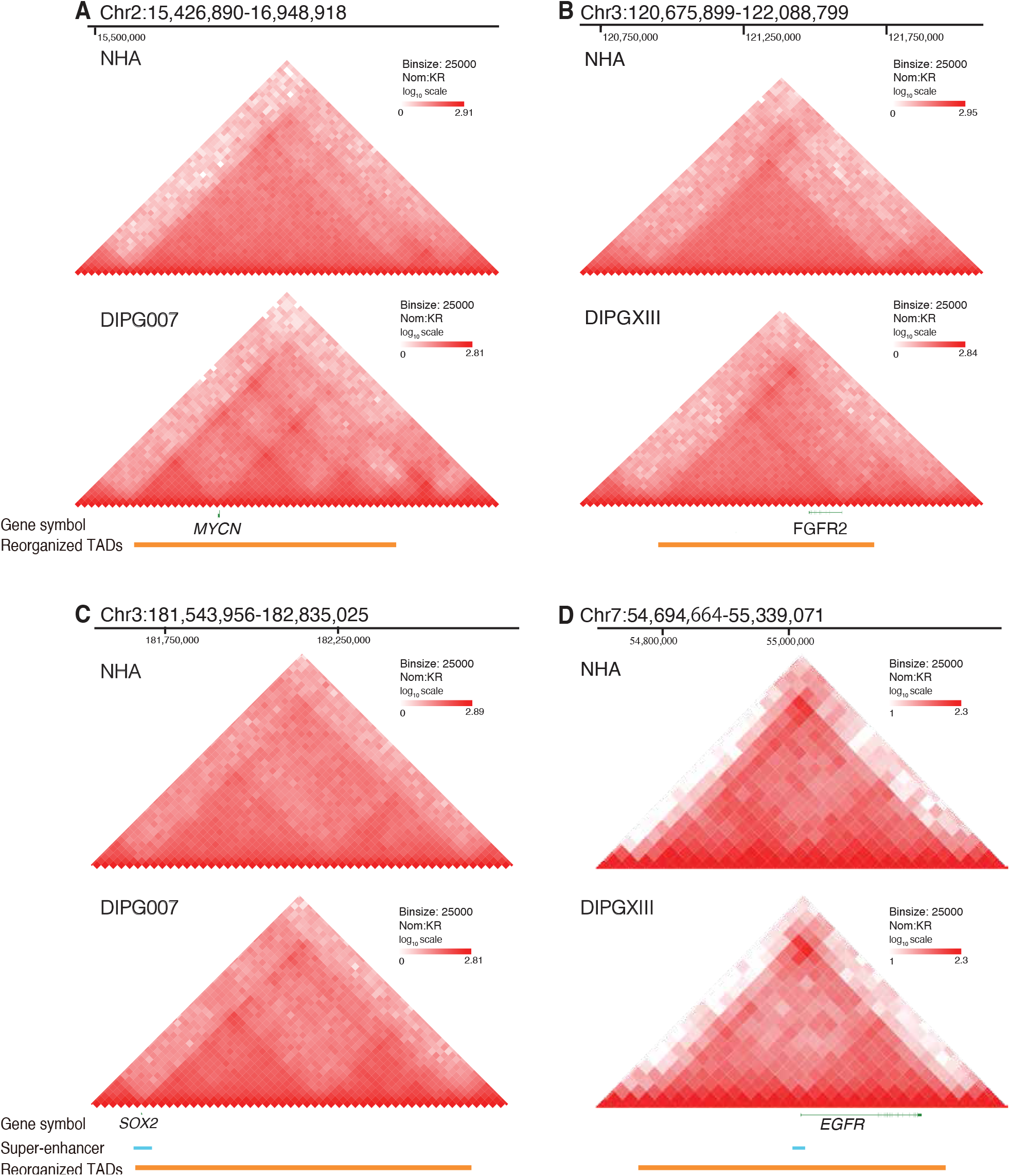
Examples of reorganized TADs in DIPG cell lines that harbor oncogenes and super-enhancers. A NHA TAD reorganized in DIPGXIII. The reorganized TAD harbors oncogene *MYCN*. (***B***) A NHA TAD reorganized in DIPG007. The reorganized TAD harbors oncogene *FGFR2*. (***C***) A NHA TAD reorganized in DIPGXIII. The reorganized TAD harbors both oncogene *SOX2* and a DIPGXIII sup-enhancer. (***D***) A NHA TAD reorganized in DIPG007. The reorganized TAD harbors oncogene *EGFR*. The upstream of the reorganized TAD contains a DIPG007 super-enhancer. Here, a super-enhancer is associated with a reorganized TAD if their 1D genomic distance is within 50 kb.

**Figure S22:**
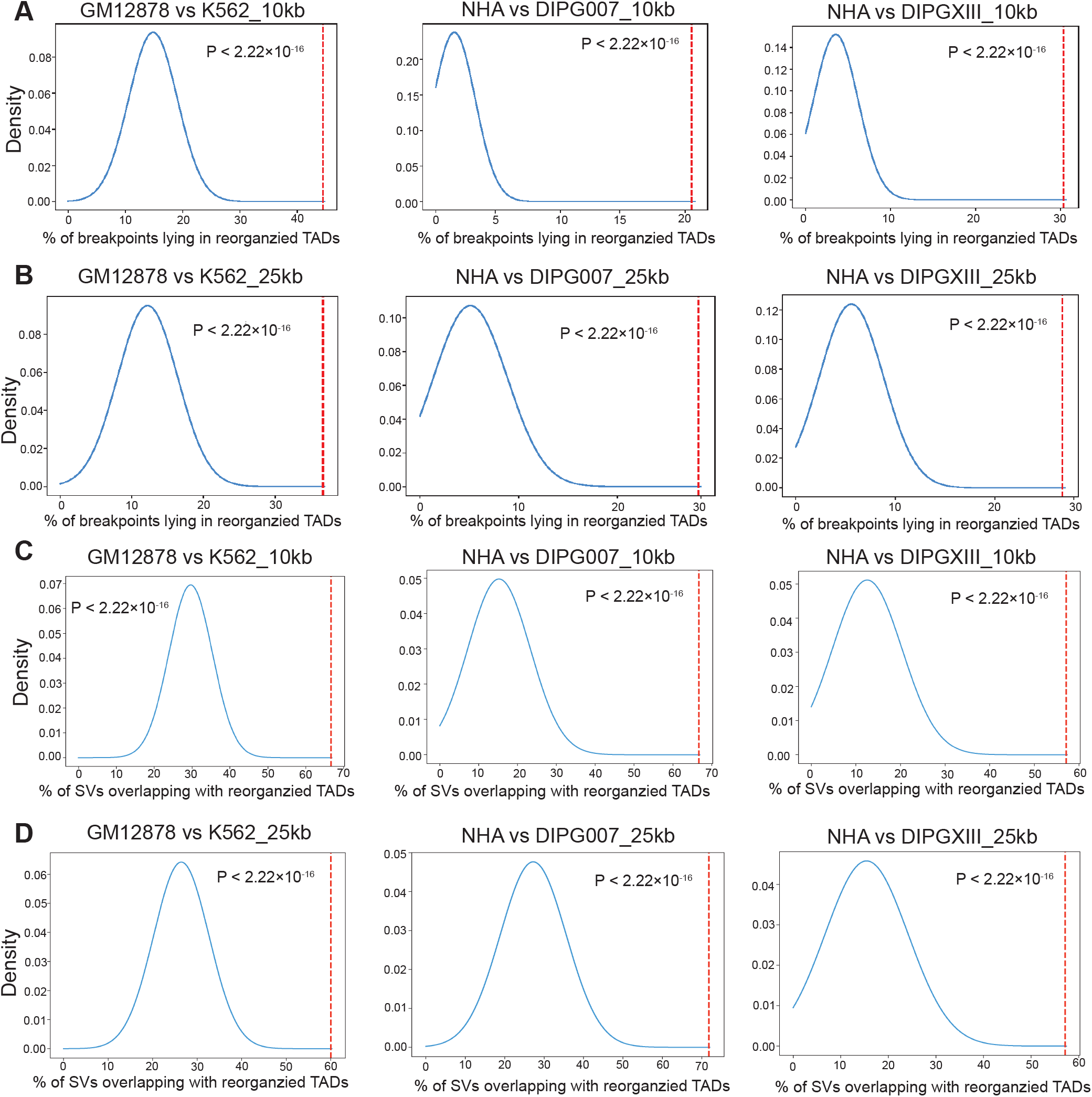
Higher proportions of SVs with reorganized TADs than random cases. (***A-B***) Density curves comparing the proportions (vertical dashed lines) of breakpoints lying in reorganized TADs with the proportions (blue curves) from random cases. The reorganized TADs are GM12878 TADs reorganized in K562, NHA TADs reorganized in DIPG007, and NHA TADs reorganized in DIGPXIII, respectively. Hi-C resolution is chosen at 10 kb (***A***) and 25 kb (***B***). (***C-D***) Density curves comparing the proportions (vertical dashed lines) of the genomic regions of SVs that overlap with reorganized TADs with the proportions (blue curves) from random cases. Hi-C resolution is chosen at 10 kb (***C***) and 25 kb (***D***). To calculate statistical significance, we randomly sample the same number of reorganized TADs and calculate the statistics. The sampling procedure is repeated 10,000 times and empirical *P* value is calculated. SVs with their genomic regions overlappping with reorganized TADs can result from either having their breakpoints lying in a reorganized TAD or containing one or more reorganized TADs (Fig. 4A). Thus, we stratified the simulation based on the two types of associations between SVs and reorganized TADs.

**Figure S23:**
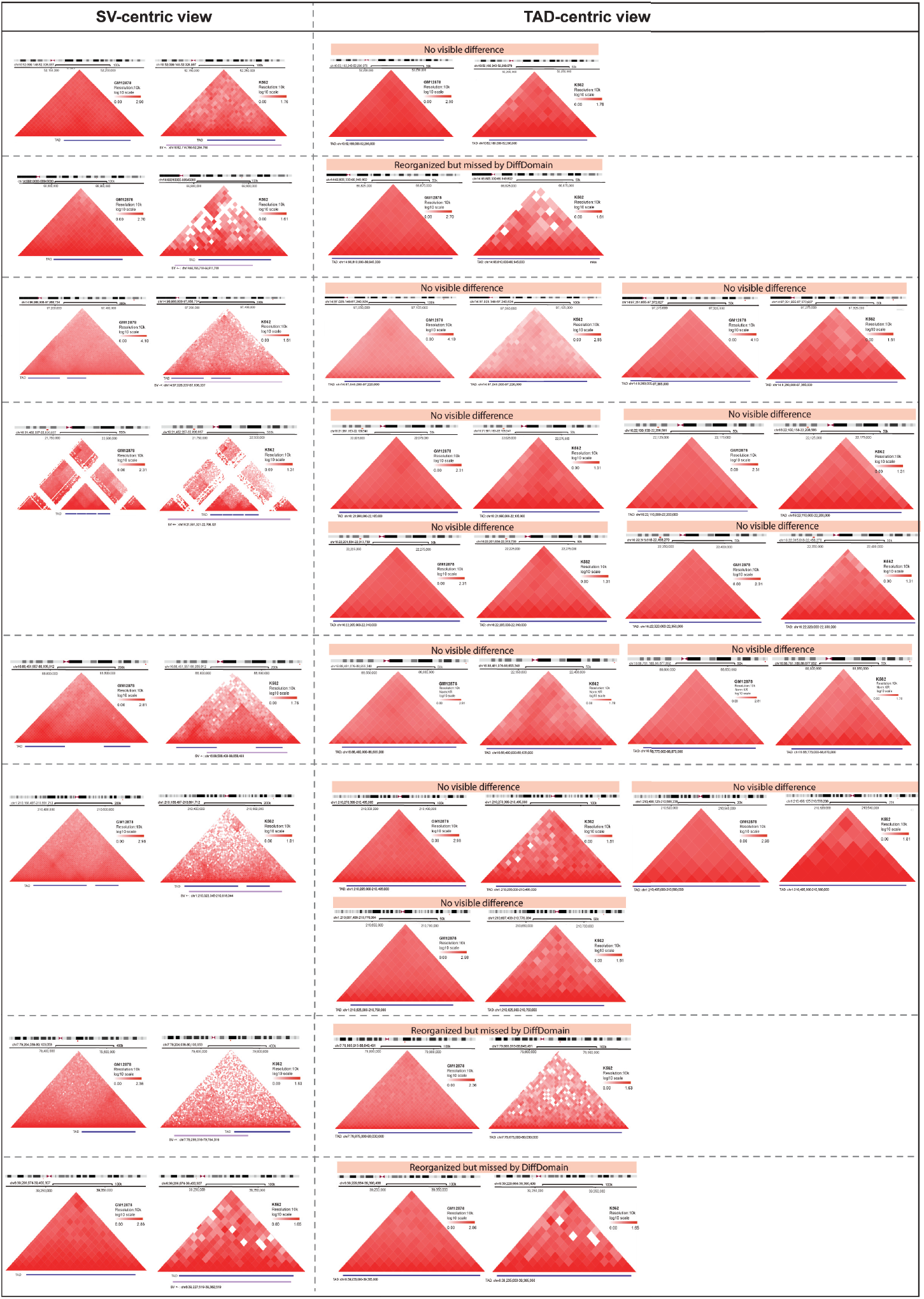
Visual examination on K562 SVs without reorganized TADs. The first two columns are SV-centric visualization, showing Hi-C contact maps from both conditions covering a region containing a specific SV and its associated TADs. First track below the heatmap represents TAD regions associated with a specific SV. Second track represents the specific SV region. Subsequent columns are TAD-centric visualization, showing a zoomed-in view of Hi-C contact maps for each of the TAD region associated with the SV region. Here, only the track representing a TAD region is shown below the heatmaps. The label “no visible difference” represents a manually draw conclusion that the TAD associated with the SV has no clear visual distinction between its two Hi-C contact maps from the two conditions, confirming DiffDomain’s results. The label “reorganized but missed by DiffDomain” represents that a manual assessment identifies visual difference between conditions for the TAD associated with the SV, although DiffDomain did not detect them. Note that, the visualization exclusively visualize SVs lacking reorganized TADs based on DiffDomain results. Among these SVs, 3 SVs have reorganized TADs but are not detected by DiffDomain. The remaining 5 SVs may lack associated TAD reorganization. The visualization is conducted using the Nucleome Browser. Abbreviations: ‘+-’, deletion;’-+’, 5^*1*^ to 3^*1*^ fusion; ‘--’, 5^*1*^ to 5^*1*^ fusion; ‘++’, 3^*1*^ to 3^*1*^ fusion.

**Figure S24:**
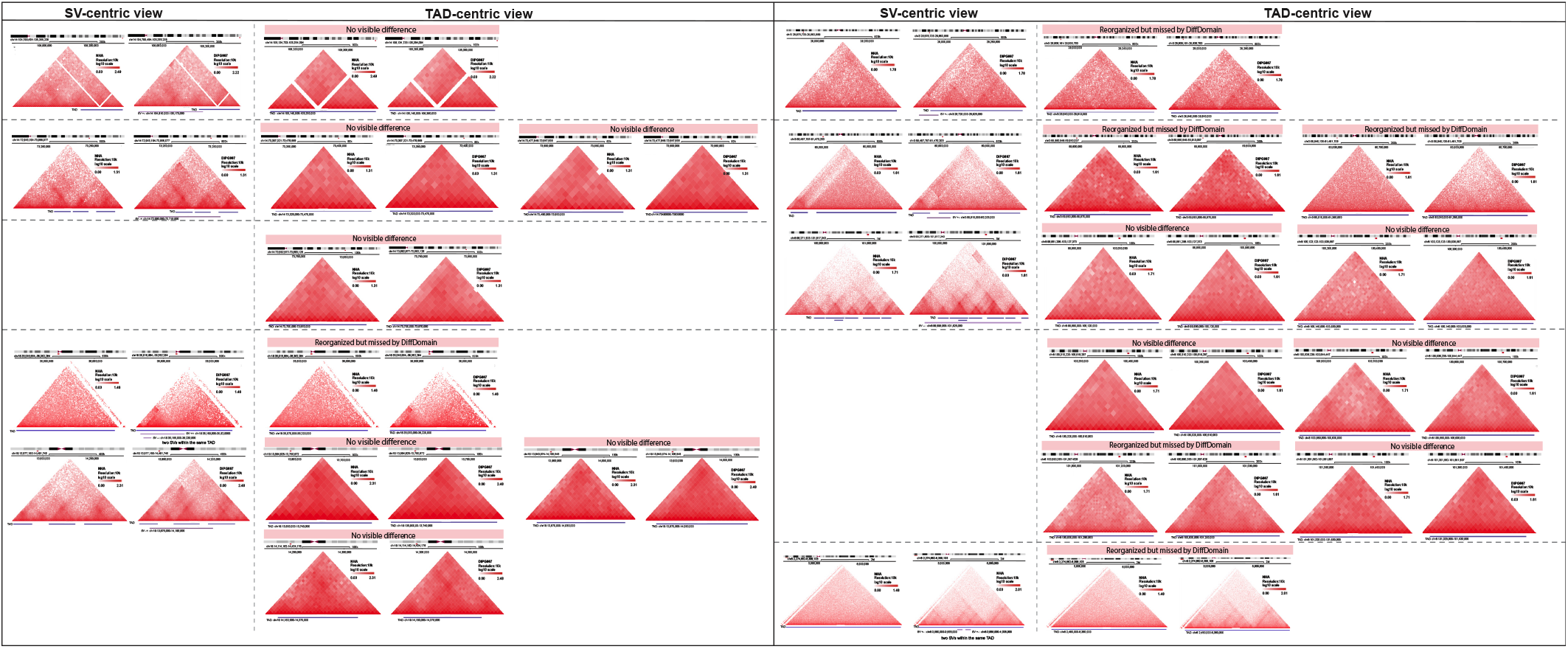
Visual examination on DIPG007 SVs without reorganized TADs. The layout scheme is the same to Supplementary Fig. S23. Note that, the visualization exclusively visualize SVs lacking reorganized TADs, as determined by DiffDomain’s results. Among these SVs, 7 DIPG007 SVs have reorganized TADs but are not detected by DiffDomain. The remaining 3 DIPG007 SVs may lack associated TAD reorganization.

**Figure S25:**
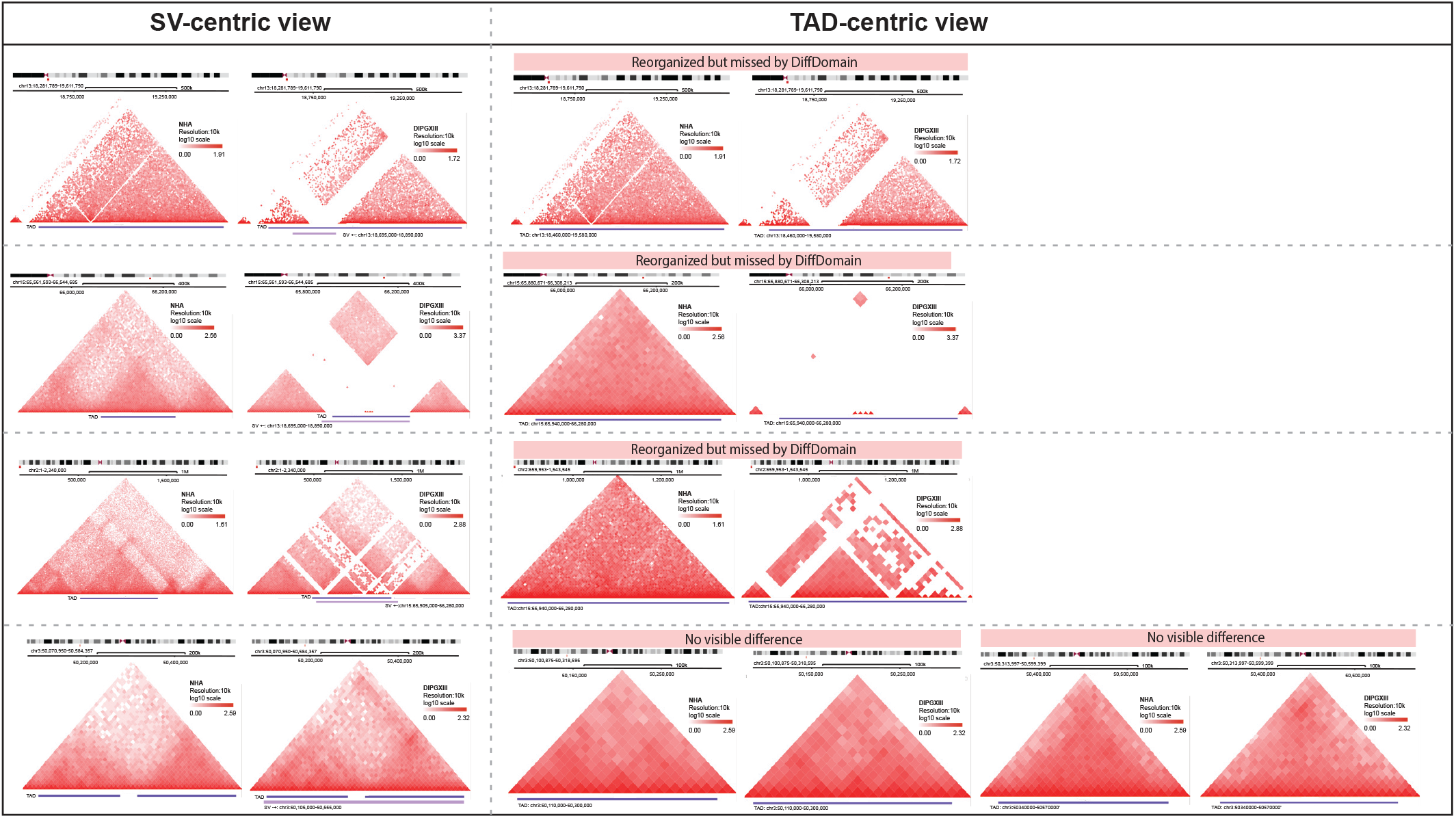
Visual examination on DIPGXIII SVs without reorganized TADs. The layout scheme is the same to Supplementary Fig. S23. Note that, the visualization exclusively visualize SVs lacking reorganized TADs, as determined by DiffDomain’s results. Among these SVs, 3 DIPG007 SVs have reorganized TADs but are not detected by DiffDomain. The remaining 1 DIPG007 SV may lack associated TAD reorganization.

**Figure S26:**
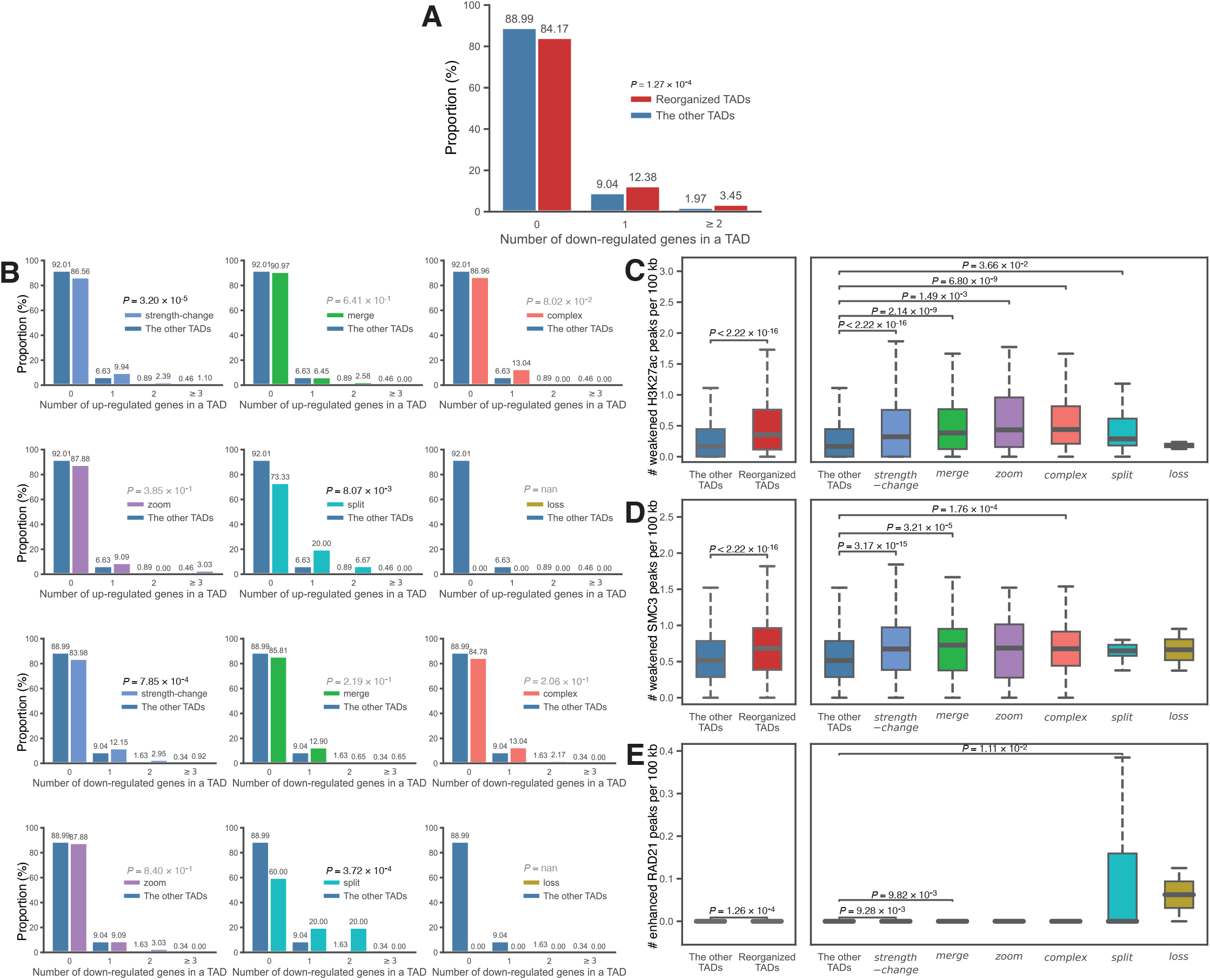
Reorganized TADs after SARS-CoV-2 infection are associated with higher numbers of downregulated genes and weakened peaks of H3K27ac, SMC3, and RAD21. (***A***) Barplot comparing the number of down-regulated genes in the reorganized TADs and the other TADs. (***B***) Barplots comparing the numbers of up- and down-regulated genes between the subtype of reorganized TADs and the other TADs. Boxplots comparing the numbers of weakened H3K27ac peaks (***C***), weakened SMC3 peaks (***D***), and enhanced RAD21 peaks (***E***) per 100 kb. *Left* : comparing reorganized TADs with the other TADs; *right* : comparison stratified by the subtypes of reorganized TADs. Note that the number of enhanced RAD21 peaks called by MAnorm2 is small, thus, the values in (***E***) are close to 0. TADs are called in mock-infected A549-ACE2 cells, reorganized TADs are called in SARS-CoV-2 infected A549-ACE2 cells. Differentially expressed genes are called by DESeq2 [92]. Differential peaks are called by MAnorm2 [93]. Boxplots are drawn by the *seaborn*.*boxplot* function in Python 3. In the box plots, the middle line represents the median; the lower and upper lines correspond to the first and third quartiles; and the upper and lower whiskers extend to values no farther than 1.5×IQR. Data is downloaded from Wang et al. [49].

**Figure S27:**
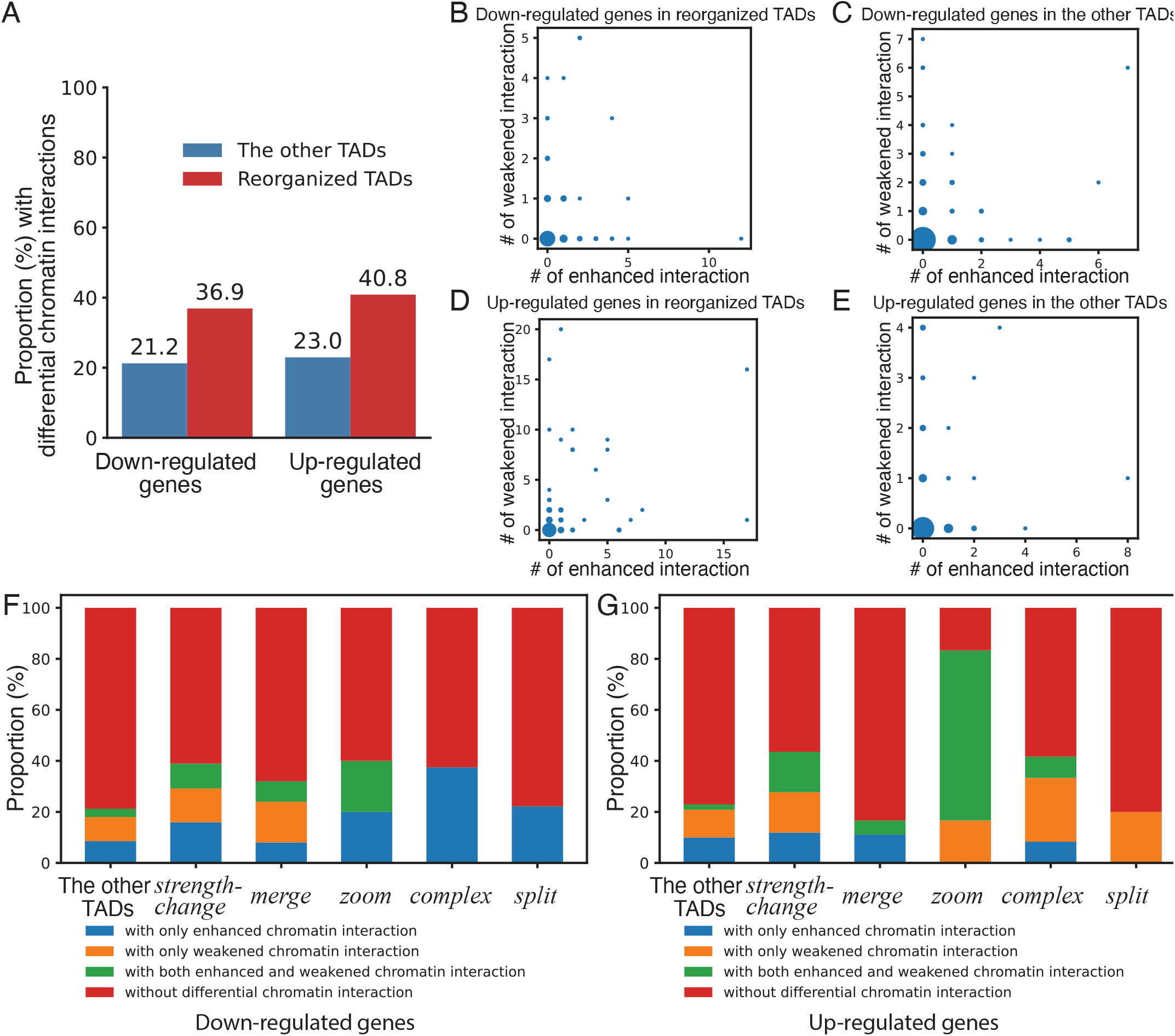
Association between differentially expressed genes and differential chromatin interactions in reorganized TADs after SARS-CoV-2 infection. (***A***) Barplot showing the proportions of down-regulated genes and up-regulated genes that have at least one differential chromatin interactions. The genes are further categorized into two groups: those located in reorganized TADs and those located in the other TADs. (***B***) Scatter plot showing the number of enhanced chromatin interactions (X-axis) against the number of weakened chromatin interactions (Y-axis) for down-regulated genes locating in reorganized TADs. Point size proportions to the number of genes. (***C***) Similar to (***B***) except that the down-regulated genes locate in the other TADs. (***D***) Scatter plot showing the number of enhanced chromatin interactions and the number of weakened chromatin interactions for up-regulated genes located in reorganized TADs. (***E***) Similar to (***D***) except that the up-regulated genes locate in the other TADs. (***F***) Stacked barplot showing the proportion of down-regulated genes with different combinations of differential chromatin interactions: with only enhanced chromatin interaction, with only weakened chromatin interaction, with both enhanced and weakened chromatin interaction, and without differential chromatin interaction. The X-axis represents the other TADs and the subtypes of reorganized TADs that harbor the down-regulatead genes. (***G***) Similar to (***F***), except for up-regulated genes.

**Figure S28:**
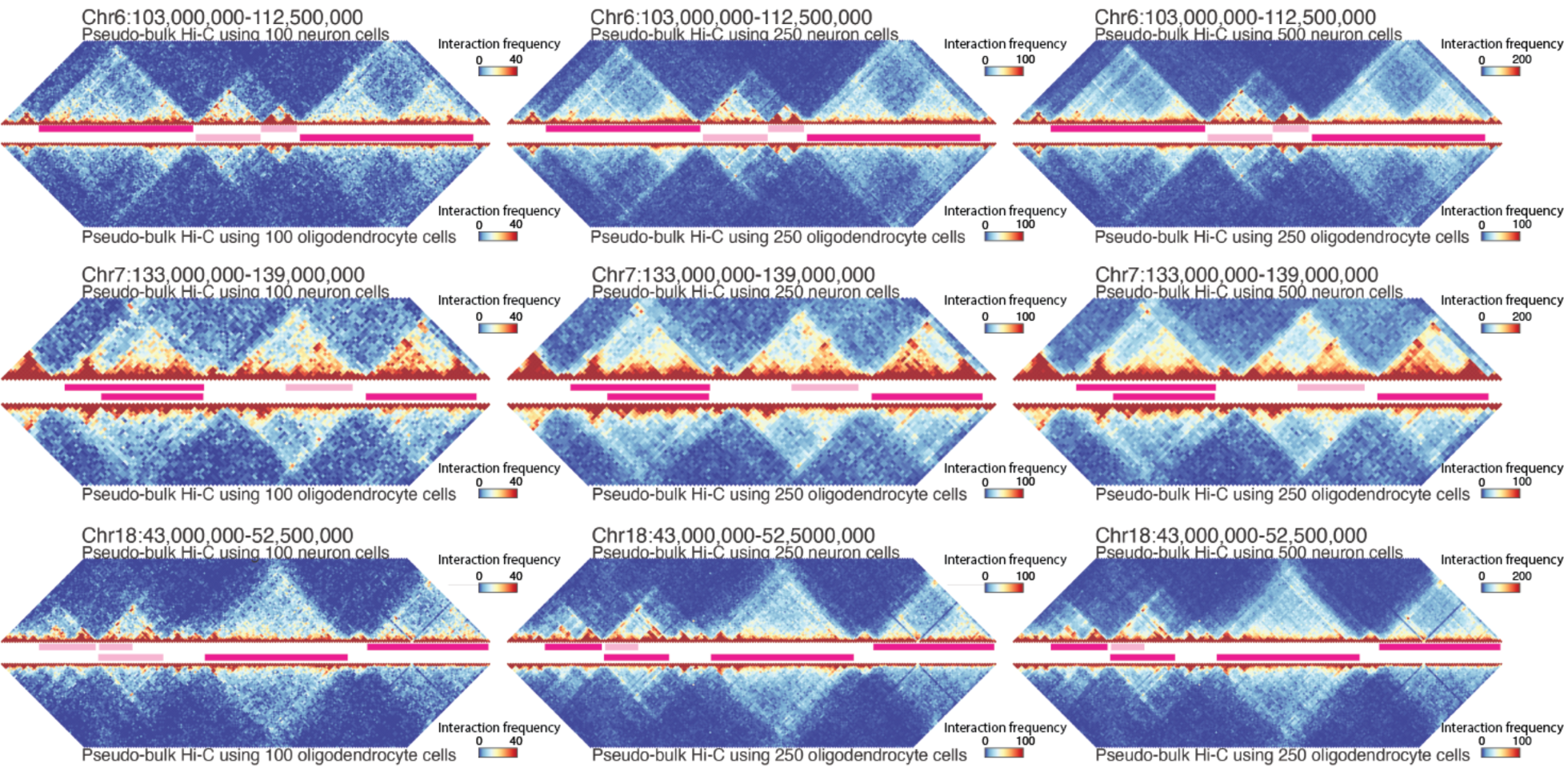
Examples of DiffDomain application to single-cell chromatin contact maps. Visualization of merged Hi-C contact maps and identified reorganized TADs (dark pink horizontal lines) in genomic regions on Chromosome 6 (*Top*), Chromosome 18 (*Middle*), and Chromosome 7 (*Bottom*). The pair of conditions are neurons and oligodendrocytes. TADs are called in neurons. The numbers of sampled cells are (100, 100), (250, 250), (500, 250), respectively.

**Figure S29:**
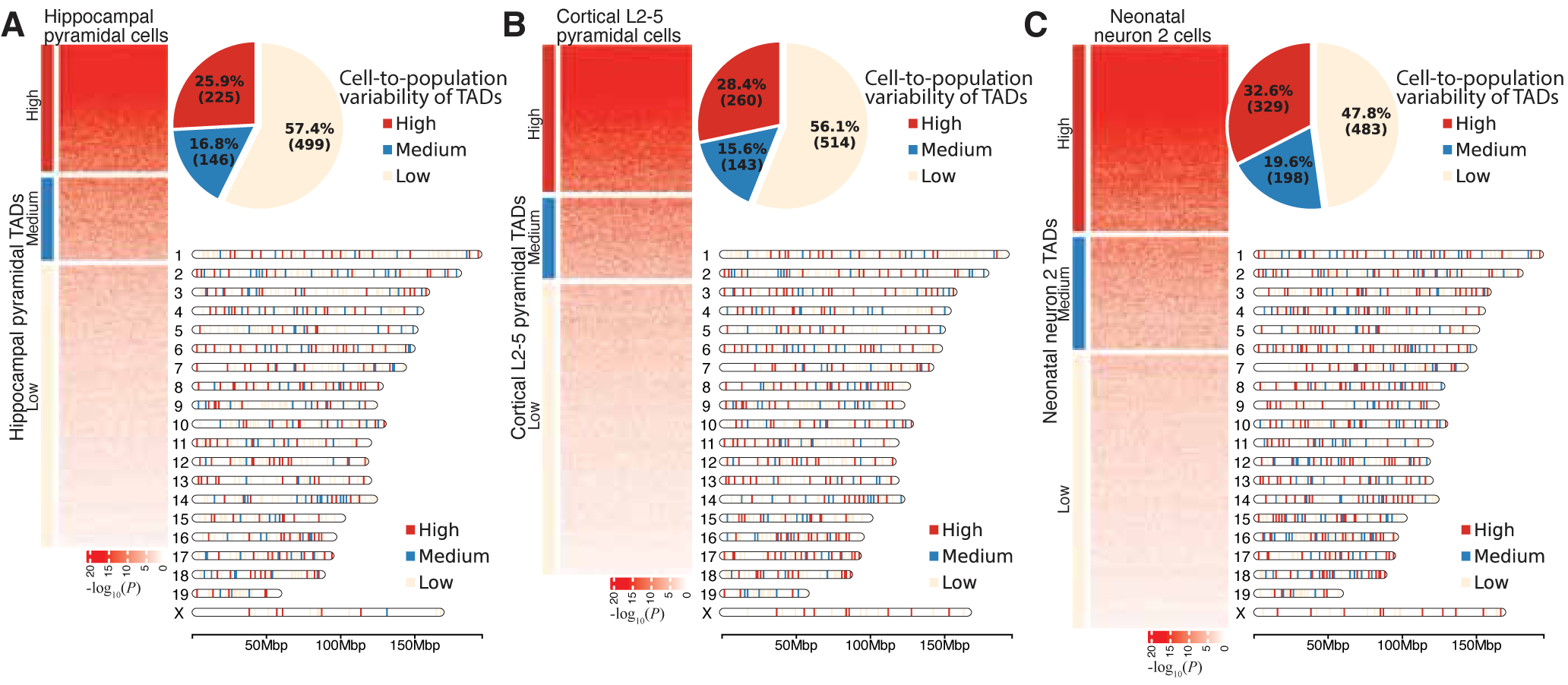
Characterization of differential cell-to-population variability of TADs in three cell types. (***A***) Characterization in hippocampal pyramidal cells. *Left* : Heatmap showing high, medium, and low cell-to-population variational TADs. Columns represent individual hippocampal pyramidal cells. Rows represent individual TADs. Values in the heatmap represent the -log_10_(*P*). *P* value is computed by DiffDomain when comparing scHi-C contact map of a TAD (row) in an individual cell (column) to the pseudo-bulk Hi-C contact map that are created using all scHi-C data from the cell type. Classification of TADs is done by hierarchical clustering. *Top right* : Pie chart showing the percentages of the high, medium, and low cell-to-population variational TADs. *Bottom right* : Chromosome map showing the genomic locations of high, medium, and low cell-to-population variational TADs.(***B***) Characterization in cortical L2-5 pyramidal cells. (***C***) Characterization in neonatal neuron 2 cells. These analyses are the repetition of the cell-to-population analysis neonatal neuron 1 cell type (Fig. 6F, G, I) to these three cell types.

**Figure S30:**
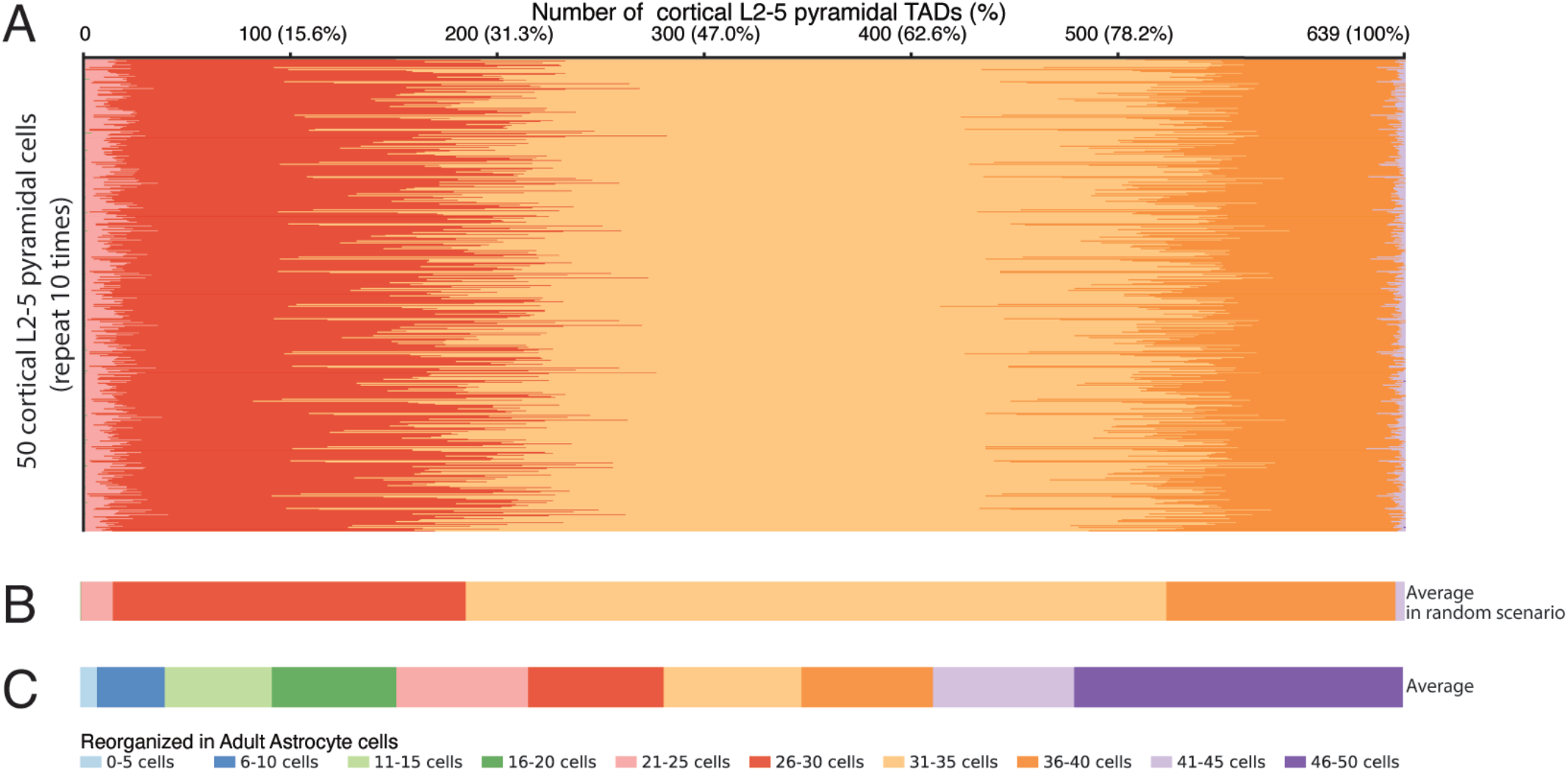
Cell-to-cell variability in random scenarios. (***A***) Stacked barplots showing the number (the percentage) of the cortical L2-5 pyramidal TADs that are reorganized in a varied number (0 to 50) of adult astrocytes in random scenarios. The layout is the same as Fig. 6K. In the simulation, equal-numbered reorganized TADs are randomly assigned in pairwise comparisons. Rows represent 50 cortical L2-5 pyramidal cells with 10 times repetition per cell. (***B***) Stacked barplot representing the average number (the percentage) in the simulation in (***A***). (***C***) Stacked bar graph representing the average number (the percentage) in real cases (also shown in Fig. 6K). This figure clearly shows that cortical L2-5 pyramidal TADs are mostly reorganized in 26-40 cells in random scenarios (***A-B***), which are much different from the patterns in real case (***C***).

